# AKT1-phosphorylated TERT assembles a FOXO3–MYC transcriptional complex that drives ERphagy and proteostasis in post-mitotic RPE

**DOI:** 10.1101/2025.11.24.690135

**Authors:** Vishnu Suresh Babu, Sayan Ghosh, Sridhar Bammidi, Ryu Jiwon, Ruchi Sharma, Dominique Meyer, Stacey Hose, Padmanabhan P. Pattabiraman, José-Alain Sahel, Nathan J. Kidley, Natércia F. Braz, Martin J. Slater, Stuart Lang, Arkasubhra Ghosh, James T. Handa, Ji Yi, Srinivasa Sripathi, Kannan Rangaramanujam, Kapil Bharti, Debasish Sinha

**Author notes:** Corresponding author: Debasish Sinha, The Johns Hopkins University, Wilmer Eye Institute, 400 N Broadway, Smith Bldg Room 6039, Baltimore, MD 21231, USA.

## Abstract

The retinal pigment epithelium (RPE) sustains lifelong proteostasis under chronic stress, yet how post-mitotic cells activate autophagy when canonical kinase pathways suppress it remains unresolved. Here we demonstrate that AKT1, conventionally regarded as an autophagy suppressor, drives autophagy through a non-canonical transcriptional mechanism triggered by isoform imbalance. In RPE and age-related macular degeneration (AMD) models, AKT2 hyperactivation destabilizes mTORC2 and engages S6K-mediated IRS-1 inhibition, creating a feedforward autophagic arrest. Compensatory AKT1 activation via mTORC2 phosphorylates telomerase reverse transcriptase (TERT) at Serine 824, driving nuclear translocation that is independent of telomere maintenance. In the nucleus, TERT assembles with FOXO3 and MYC into a tripartite transcriptional complex that occupies the *EIF2AK3* promoter, enabling PERK transcriptional activation. This activity converts the unfolded protein response from pro-apoptotic to cytoprotective: PERK–ATF4 signaling drives biogenesis of core autophagy machinery while simultaneously inducing selective ERphagy through the receptors TEX264 and CCPG1, which prevents pathological PERK clustering and preserves tubular ER architecture in diseased RPE. Using *C. elegans* phosphorylation-deficient mutants, mouse models and AMD patient induced pluripotent stem cell-derived retinal pigment epithelium (iPSC-RPE), we establish that AKT1-mediated TERT phosphorylation is an evolutionarily conserved prerequisite for FOXO/DAF-16 nuclear function and lysosomal homeostasis in post-mitotic cells. Pharmacologic targeting of AKT2 with a first-in-class dual-pocket allosteric inhibitor selectively enhances AKT1 compensation, restoring macroautophagic and ERphagy flux across disease models. These findings reveal a kinase-to-transcription axis that reprograms organelle quality control and identify the AKT1–TERT–PERK–ATF4 pathway as a therapeutic target in proteostasis-driven disease.

## Introduction

The retinal pigment epithelium (RPE) is a high-performance metabolic monolayer, essential for the maintenance of adjacent photoreceptors and maintains the structural integrity of the outer blood-retinal barrier[1, 2]. Tasked with the lifelong daily phagocytosis of shed photoreceptor outer segments, the RPE operates under a state of chronic oxidative stress and immense degradative demand[3, 4]. Consequently, the RPE is an ideal cell type to study the consequences of progressive collapse of the autophagy-lysosomal pathway (ALP) on cellular health. Not surprisingly, ALP dysfunction plays a central and early event in RPE aging and age-related macular degeneration (AMD)[5–8]. While the accumulation of proteotoxic waste signals the transition to disease[9–11], the upstream signaling dynamics that govern the switch from healthy proteostatic surveillance to maladaptive organelle failure remain poorly understood[12–15]. Central to this metabolic governance is the mechanistic target of rapamycin (mTOR), which exists in two structurally and functionally distinct complexes: mTORC1, which suppresses autophagy in response to nutrient abundance, and mTORC2, which regulates cytoskeletal organization and promotes pro-survival signaling[16, 17]. Under homeostatic conditions, these complexes exist in a delicate equilibrium[18, 19]. However, in the context of RPE aging and AMD, this balance is disrupted[20, 21].

The PI3K/AKT pathway is the canonical autophagy suppressor[22]. Ironically, mammals express three AKT isoforms with distinct tissue distributions and divergent, often opposing, cellular functions[23]. How cells discriminate between AKT isoform signals to produce opposing outcomes in autophagy regulation remains a fundamental, unresolved question. We recently reported an unexpected regulatory relationship between AKT1 and AKT2 in the RPE[24]. Genetic deletion of AKT2 paradoxically triggered compensatory upregulation of AKT1 that enhanced autophagy and protected against cellular dysfunction, directly contradicting the established view that AKT activation suppresses autophagy[25]. We further demonstrated that AKT2 drives RPE degeneration through lysosomal and mitochondrial dysfunction, establishing the two isoforms as opposing regulators of cell fate[26]. The molecular mechanism enabling AKT1 to promote rather than inhibit autophagy remained unknown. Notably, MYC, a transcription factor with established roles in metabolic reprogramming[27] and known physical interactions with both FOXO3 and TERT[28, 29], had never been considered in the context of AKT isoform-specific signaling or autophagy regulation in post-mitotic cells. More broadly, whether this paradox reflects a fundamental principle of stress-response signaling or a cell-type-specific adaptation had not been addressed.

Resolving this paradox requires understanding how AKT1 compensation connects to the transcriptional machinery that regulates autophagy, lysosomal function, and ER quality control. First, the transcription factor FOXO3 is a well-recognized activator of autophagy genes[30, 31]. Under normal conditions, AKT phosphorylates FOXO3 to retain it in the cytoplasm and prevent autophagy induction[32]. However, FOXO3 does not act alone, rather, it functions within multi-protein complexes[32, 33]. The cofactors that determine FOXO3’s transcriptional specificity in the context of AKT isoform compensation has not been identified. Whether MYC could serve as such a cofactor[34], bridging FOXO3 to the broader transcriptional machinery governing organelle quality control has not been explored. Second, telomerase reverse transcriptase (TERT) has been recognized with functions that extend well beyond telomere maintenance[35, 36]. In the nucleus, TERT can regulate transcription in a telomere-independent manner, and relevant to this study, is a known direct substrate of AKT[37]. The consequences of AKT-mediated phosphorylation of TERT in a post-mitotic cell that does not rely on telomerase for replication, is an open question. Third, the PERK–eIF2–ATF4 arm of the Unfolded Protein Response (UPR)[38] can activate both general macroautophagy[39] and selective ERphagy, the targeted lysosomal degradation of damaged endoplasmic reticulum (ER), under ER stress conditions[40]. Whether this selective clearance program is transcriptionally licensed by upstream kinase signaling, and specifically by the AKT isoform balance that regulates RPE survival, had not been investigated. Together, these three open questions pointed toward an unrecognized signaling hierarchy connecting AKT1 compensation, a FOXO3–MYC–TERT nuclear complex, and PERK-driven adaptive proteostasis encompassing both macroautophagy and selective ERphagy.

Here, we define the molecular mechanism linking AKT1 activation to autophagy and ERphagy. Using transcriptomic profiling of the *cryba1* conditional knockout (cKO) RPE, a model that recapitulates the proteostatic defects of human dry AMD[5, 41, 42], alongside RPE-specific *Akt2* knock-in and knockout models[24, 26], human induced pluripotent stem cell-derived RPE (iPSC-RPE) from AMD donors[43], engineered *C. elegans* mutants, and phosphoproteomic approaches, we demonstrate that AKT1 phosphorylates TERT at Serine 824, driving its nuclear translocation[44] and assembly into a previously uncharacterized tripartite complex with FOXO3 and MYC. This complex directly activates PERK, initiating a self-amplifying loop in which PERK–ATF4 signaling drives the biogenesis of autophagy machinery while simultaneously inducing selective ERphagy through the receptors TEX264 and CCPG1[45]. We show that this AKT1–TERT axis is evolutionarily conserved and essential for proteostasis in post-mitotic cells. To leverage these mechanistic insights therapeutically, we employed a structure-based design to develop a first-in-class dual-pocket allosteric AKT2 inhibitor that selectively triggers beneficial AKT1 compensation, restoring autophagy and ERphagy flux in diseased human RPE. These findings establish a previously unrecognized transcriptional mechanism linking isoform-specific kinase signaling to organelle quality control and identify the AKT1–TERT–PERK–ATF4 axis as a therapeutic target for diseases of defective ER quality control and proteostatic failure.

## Results

### A reciprocal AKT1/AKT2 signaling axis governs RPE homeostasis and maladaptation

Lysosomal and autophagic dysfunction is a defining cellular lesion of RPE aging and AMD[5]. Prior work established AKT2 hyperactivation in the RPE as a driver of lysosomal and mitochondrial dysfunction, which severely compromises the autophagy-lysosomal pathway that leads to toxic waste accumulation[26]. While genetic deletion of AKT2 triggers a compensatory and essential AKT1 upregulation to maintain cellular homeostasis[24], the molecular transition between these states remains poorly defined. To identify the upstream signaling events driving lysosomal and autophagy suppression before phenotypic degeneration is established, we performed transcriptomic profiling of *cryba1* conditional knockout (cKO) RPE, a model that endo-phenocopies the proteostatic defects of human dry AMD [5, 41, 42]. RNAseq analysis across the 5- (pre-symptomatic) and 10-month (post-symptomatic) window revealed a striking isoform switch that showcased *Akt2* induction coupled with *Akt1* reduction, alongside elevated expression of mTOR components including *Rptor*, *Rictor* and *Mlst8* (Fig. 1A). Gene ontology (GO) enrichment confirmed a maladaptive signature of hyperactive RTK/PI3K/AKT signaling alongside suppressed unfolded protein response (UPR) and macro autophagy (Fig. S1A and B). The negative correlation between *Akt1* and *Akt2* expression shift (R² = −0.6; Fig. S1C) is sustained at the protein level for at least 18 months (Fig. S1D) and this signaling shift precedes overt phenotypic degeneration, such as basal laminar deposits[46], marking a critical window where mTORC1-driven translational programs override autophagic surveillance (Fig. S1E, F).

**Figure 1.**
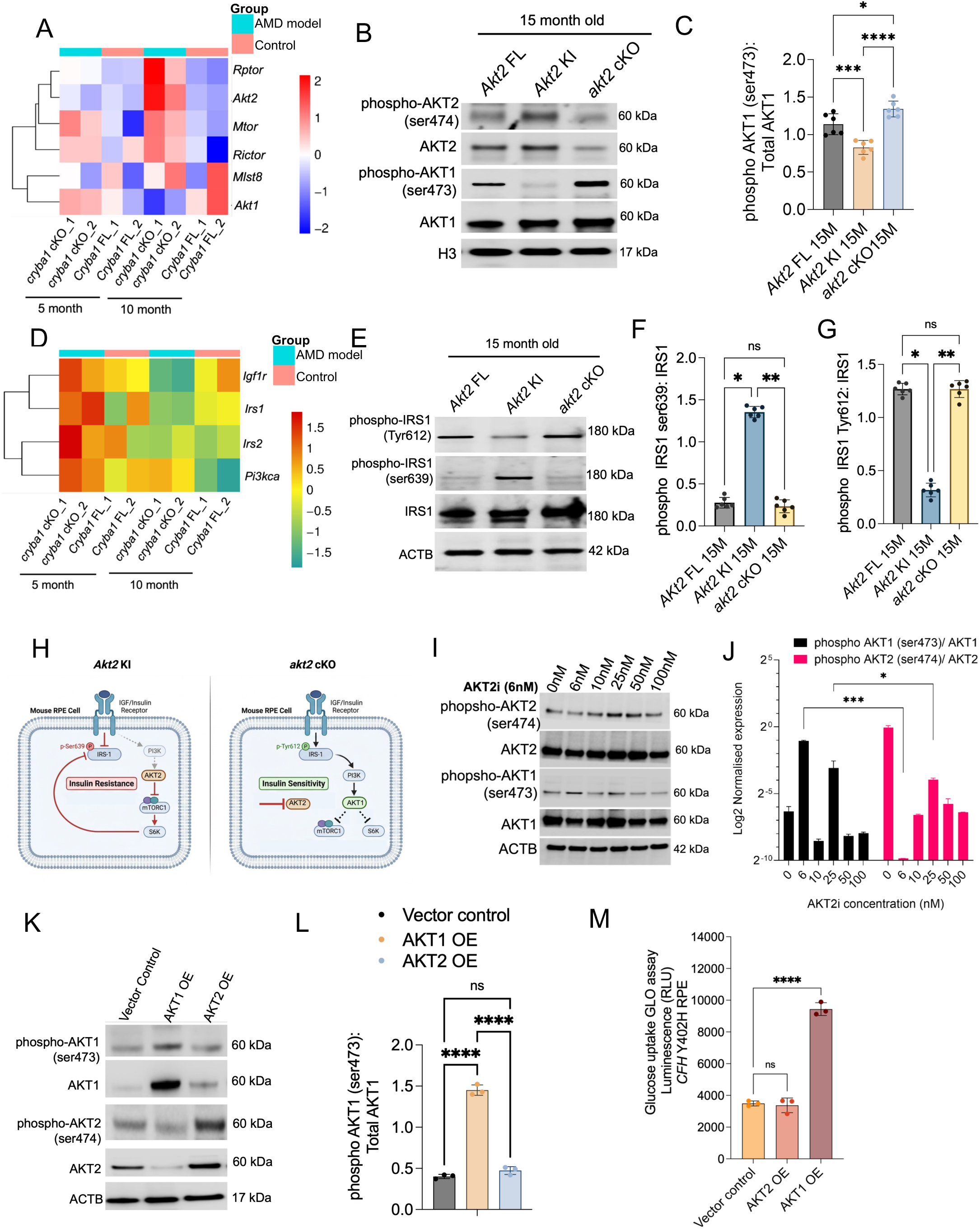
AKT2 hyperactivation drives RPE insulin resistance via IRS-1 inhibitory feedback. (**A**) Heatmaps showing differential mRNA expression of mTOR-AKT pathway components in RPE tissues from *cryba1* cKO (AMD model) and control mice at 5 and 10 months of age. Data represent relative expression (Z-score); n = 4 animals per group. (**B**) Representative immunoblots of phospho-AKT2 (Ser474) and phospho-AKT1 (Ser473) levels in RPE lysates from 15-month-old *Akt2* flox (FL), *Akt2* knock-in (KI), and *akt2* conditional knockout (cKO) mice. Histone H3 (H3) serves as a loading control. (**C**) Densitometry quantification of the immunoblots in (B). Data are mean ± SD; n = 6 independent biological replicates. (**D**) Heatmaps showing differential mRNA expression of IGFR-IRS-PI3K pathway components in RPE tissues from *cryba1* cKO and control mice at 5 and 10 months of age. Data represent relative expression (Z-score). (**E**) Representative immunoblots of inhibitory phospho-IRS1 (Ser639) and activating phospho-IRS1 (Tyr612) levels in RPE lysates from 15-month-old *Akt2* FL, *Akt2* KI, and *akt2* cKO mice. ACTB serves as a loading control. (**F**) Densitometry quantification of phospho-IRS1 (Ser639) immunoblots shown in (**E**). (**G**) Densitometry quantification of activating phospho-IRS1 (Tyr612) immunoblots shown in (**E**). (**H**) Schematic models illustrating the transition from insulin resistance in Akt2 KI RPE (left) to insulin sensitivity in *akt2* cKO RPE (right). AKT2 hyperactivation triggers a negative feedback loop via RPS6KB1/S6K to phosphorylate IRS1 at Ser639, suppressing PI3K-AKT1 signaling. (**I**) Dose-response western blots of AKT1 and AKT2 phosphorylation in human iPSC-RPE cells treated with increasing concentrations (0 to 100 nM) of the AKT2-selective inhibitor CCT128930 for 24 h. Note the preferential increase of AKT1 phosphorylation at the 6 nM dose with subsequent reduction in AKT2 phosphorylation. (**J**) Densitometry quantification of phospho-AKT1 (Ser473) and phospho-AKT2 (Ser474) levels across the indicated concentrations of AKT2 inhibitor. (**K**) Representative immunoblot analysis of human iPSC-RPE *CFH* ^Y402H^ cells overexpressing (OE) AKT1 or AKT2 compared to vector control. ACTB serves as a loading control. (**L**) Densitometry analysis of phospho-AKT1 to total AKT1 levels in control, AKT1 OE and AKT2 OE conditions in iPSC RPE cells (**M**) Glucose uptake GLO luminescence assay in *CFH* ^Y402H^ RPE cells. AKT2 OE and vector control impair glucose uptake, while AKT1 OE significantly enhances glucose uptake. For all graphs, data are represented as mean ± SD. Statistical significance was determined by one-way ANOVA with Tukey’s post-hoc test; *p < 0.05, **p < 0.01, ***p < 0.001, ****p < 0.0001. ns, non-significant.

Consistent with this shift, the RPE from *Akt2* knock-in (KI) mice showed markedly elevated p-AKT2 (Ser474) with reciprocal suppression of p-AKT1 (Ser473) relative to litter mate controls (Fig 1B, C) and differential regulation of the IGFR1/IRS-1/PI3K axis (Fig. 1D). The feedforward mechanism underlying autophagy arrest operates through the mTORC1 downstream kinase S6K, which phosphorylates IRS-1 at the inhibitory Ser639 site[47] (Fig. 1E, F). This steric inhibition prevents IRS-1 from interacting with the insulin receptor, thereby blocking PI3K-AKT1 activation[48], a pathway we show is essential for maintaining autophagic flux and glucose homeostasis (Fig. 1E, F). Elevated inhibitory p-IRS-1 (Ser639) and suppressed activating p-IRS-1 (Tyr612)[49] were confirmed in *Akt2* KI RPE (Fig. 1E, F, G) and both metabolic decompensation and subsequent autophagic suppression were fully reversed in *akt2* cKO RPE (Fig. 1F, G), identifying the AKT2/mTORC1/S6K axis as the primary driver of RPE insulin resistance (Fig. 1H).

To test the therapeutic potential of restoring this imbalance, we used the selective AKT2 inhibitor CCT128930 (AKT2i)[50]. In human iPSC-RPE, a precise 6 nM dose preferentially inhibited AKT2, which disinhibited the IRS1/PI3K/AKT1 pathway (Fig. 1I, J) restoring expression of insulin-sensitizing and pro-survival components in *CFH*^Y402H^ RPE (Fig. S1G). Consistent with this, AKT1 overexpression alone significantly enhanced glucose uptake activity, while AKT2 hyperactivation antagonizes glucose uptake through S6K-mediated feedback inhibition (Fig. 1K, L, M, Fig. S1H). These data establish AKT1 as an essential driver of RPE homeostasis and AKT2 hyperactivation acts as a molecular switch that shunts the cell toward mTORC1-mediated autophagic arrest and degeneration.

### Spatial segregation of AKT isoforms into distinct mTOR complexes drives mTORC1/mTORC2 imbalance

Proteomic analysis of *cryba1* cKO RPE identified altered abundance of mTORC1-trafficiking regulators, including STX7[51], BIN1[52], and FLCN[53] (Fig. S2A), consistent with mTORC1 hyperactivation as a conserved driver of RPE autophagic failure (Fig. S2B). In *CFH* ^Y402H^ iPSC-RPE from three AMD donors compared to three unaffected sibling controls, we observed a transcriptomic signature defined by the suppressed macroautophagy, ER protein processing and lysosomal function, coupled with upregulated inflammatory and apoptotic programs (Fig. S2C), mirroring defects in *Akt2* KI and *cryba1* cKO models and highlights a shared potential mechanistic origin. Pharmacological AKT2 inhibition in three AMD donor cells partially restored the transcriptome, upregulating SNARE-mediated transport and ER protein processing (Fig. S2D, E) while downregulating mTORC1 scaffolding components like *MLST8*, *LAMPTOR4*, *CASTOR1*[54–56] (Fig. 2I). This rebalancing suggests that AKT2 inhibition influences the compensatory AKT1 upregulation required to restore the mTORC1/mTORC2 equilibrium.

**Figure 2.**
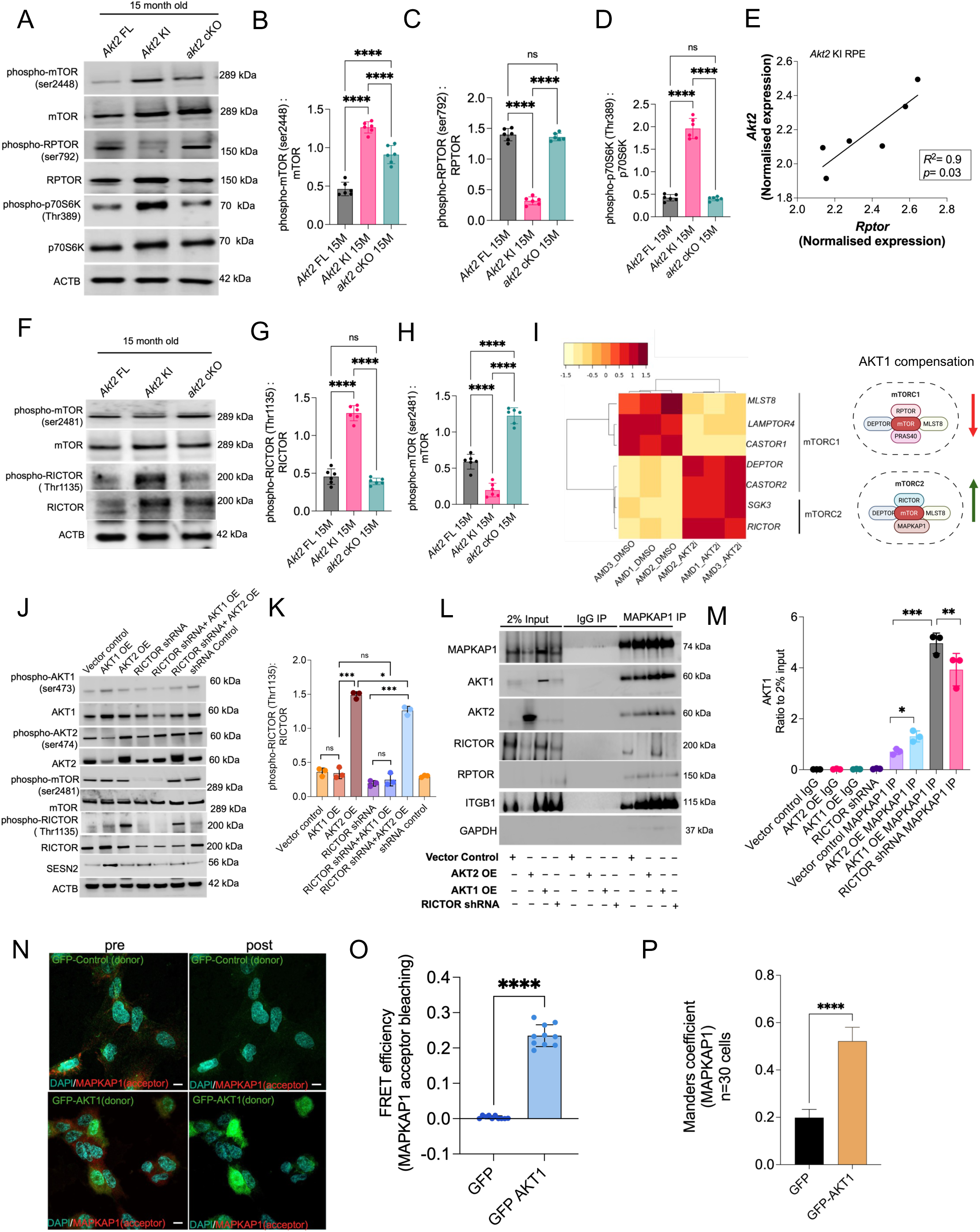
Isoform-specific regulation of mTORC1 and mTORC2 complexes by AKT1 and AKT2 in the RPE. (**A**) Representative immunoblot analysis of mTORC1 activation markers, including phospho-mTOR (Ser2448), the mTORC1 downstream target phospho-RPS6KB1/p70S6K (Thr389), and the inhibitory marker phospho-RPTOR/RPTOR (Ser792) in RPE lysates from 15-month-old *Akt2* FL, *Akt2* KI, and *akt2* cKO mice. AKT2 hyperactivation in *Akt2* KI significantly correlates with increased mTORC1 downstream signaling. (**B–D**) Densitometry quantification of (**B**) phospho-mTOR (Ser2448), (**C**) phospho-RPTOR (Ser792), and (**D**) phospho-p70S6K (Thr389) immunoblots; n = 6 independent biological replicates per group. (**E**) Correlation analysis of qPCR-validated *Akt2* and *Rptor* expression in *Akt2* KI RPE, normalized to *Actb* controls (R^2^ = 0.8, p = 0.02). (**F**) Representative immunoblot of the mTORC2 activation marker phospho-mTOR (Ser2481) and the inhibitory marker phospho-RICTOR (Thr1135). Note that while *Akt2* KI drives RICTOR phosphorylation at Thr1135, *akt2* cKO enhances the autophosphorylation of mTOR at Ser2481, suggesting a shift toward intact mTORC2 assembly. (**G, H**) Densitometry quantification of (**G**) phospho-RICTOR (Thr1135) and (**H**) phospho-mTOR (Ser2481) immunoblots; n = 6 independent biological replicates per group. (**I**) Heatmap showing differential expression of mTORC1 and mTORC2 regulatory subunits in human iPSC-RPE CFH ^Y402H^ cells treated with DMSO or AKT2i (6 nM). Schematic (right) illustrates the “AKT1 compensation” model where mTORC1 activity is suppressed (red arrow) and mTORC2 activity is enhanced (green arrow). (**J**) Immunoblot analysis of mTOR complex components in human iPSC-RPE CFH ^Y402H^ cells following overexpression (OE) of AKT1 or AKT2, and shRNA-mediated knockdown of RICTOR. Total AKT1 and phospho-AKT1 (Ser473) levels were reduced upon RICTOR knockdown, while AKT2 remained unaffected. AKT2 specifically promotes RICTOR phosphorylation at Thr1135 in a RICTOR-independent manner. AKT1 OE elevates SESN2/Sestrin2 expression, which was reduced upon RICTOR knockdown. ACTB serves as a loading control. (**K**) Densitometry quantification of phospho-RICTOR (Thr1135) immunoblots from (**J**). (**L**) Co-immunoprecipitation of MAPKAP1 in plasma membrane fraction showing strong interaction with AKT1 and RICTOR in AKT1 OE conditions and not AKT2 in AKT2 OE conditions in *CFH* ^Y402H^ RPE cells. The interaction of MAPKAP1 with AKT1 was slightly reduced in RICTOR knockdown condition (**M**) Co-IP quantification showing increased AKT1 interaction with MAPKAP1 in AKT1 OE condition, no significant interaction in control or AKT2 OE and slight reduction in MAPKAP1-AKT1 interaction upon RICTOR knockdown. (**N**) Representative FRET imaging of RPE cells expressing GFP-control or GFP-AKT1 (donor) and stained for mCherry-MAPKAP1/MAPKAP1 (acceptor). (**O**) Quantification of FRET efficiency via acceptor photobleaching of mCherry-MAPKAP1 (n = 10 cells). (**P**) Manders’ co-localization coefficient (n = 30 cells) demonstrates a direct and specific physical interaction between AKT1 and the mTORC2 subunit MAPKAP1. Scale bars: 10μm. Data are represented as mean ± SD. Statistical significance was determined by one-way ANOVA with Tukey’s post-hoc test; *p < 0.05, **p < 0.01, ***p < 0.001, ****p < 0.0001. ns, non-significant.

The biochemical basis for this differential autophagic process lies in the spatial partitioning of the two AKT isoforms. We found that AKT2 hyperactivation at the lysosome drives constitutive mTORC1 signaling, as evidenced by elevated p-mTOR (Ser2448)[57] and p-p70S6K (Thr389)[58], the suppression of the inhibitory p-RPTOR (Ser792)[59] and strong positive correlation of AKT2-RPTOR gene expression (R^2^=0.8, *p*=0.02)(Fig. 2A–E). Conversely, AKT2 hyperactivation destabilizes the pro-autophagic mTORC2 complex through inhibitory phosphorylation of RICTOR at Thr1135[60] (Fig. 2F–H). Epistatic experiments using RICTOR shRNA confirmed the specific dependency of this signaling axis. Knockdown of RICTOR selectively reduced phospho-AKT1 (Ser473) without affecting AKT2, whereas AKT2 overexpression (OE) specifically drove RICTOR phosphorylation at the inhibitory Thr1135 site (Fig. 2J, K) in parallel with elevated mTORC1 specific p-mTOR (Ser2448) and activation of its downstream targets S6K and 4E-BP1[61] (Fig. S2F, G). AKT1 overexpression (OE) increased SESN2/Sestrin2, a potent mTORC1 inhibitor[62] and suppressed the lysosomal amino acid transporter SLC36A4[63] in a RICTOR-dependent manner (Fig. S2F, H), directly linking mTORC2 integrity to lysosomal amino acid sensing that activates mTORC1.

Isoform-specific co-immunoprecipitation confirmed the spatial basis of this rebalancing: mTORC2 specific MAPKAP1 co-IP [64] and RICTOR co-IP[65] from plasma membrane fractions showed high AKT1/RICTOR and low AKT2/RICTOR association (Fig. 2L, M; Fig. S2I, J), while RPTOR co-IP[66] from lysosomal fractions had the exact opposite (Fig. S2K-M). These results demonstrate that AKT1 engages mTORC2 at the plasma membrane while AKT2 drives mTORC1 at the lysosome.

To determine whether AKT1 physically docks onto the mTORC2 complex, Förster Resonance Energy Transfer (FRET) acceptor photobleaching[67] was performed in iPSC-RPE *CFH*^Y402H^ cells. Using GFP-AKT1 as the donor and mCherry-MAPKAP1, high FRET efficiency was observed for GFP-AKT1 upon MAPKAP1 acceptor bleaching (Fig. 2N, O), while cells expressing the GFP-control showed no significant energy transfer, confirming the direct physical interaction between AKT1 and MAPKAP1 and identifying MAPKAP1 as a critical mTORC2 docking node for AKT1 recruitment. Greater co-localization for GFP-AKT1/mCherry-MAPKAP1 versus GFP/mCherry-MAPKAP1-control across 30 cells was confirmed by Manders co-localization analysis (Fig. 2P). The dependence of the specific isoform is absolute, as a reciprocal mCherry-AKT2/magenta-MAPKAP1 FRET experiment showed no significant energy transfer (Fig. S2N, O). These results demonstrate that mTORC2 engagement is exclusive to AKT1 and provide a structural basis for the spatial segregation of AKT isoforms and explains why AKT2 hyperactivation fails to compensate for and actively antagonizes AKT1-mTORC2-mediated cellular survival.

### The AKT1/AKT2 isoform switch controls non-canonical TERT nuclear import

Transcriptome analysis of *cryba1* cKO RPE identified AKT1 and TERT as core downregulated nodes (Fig. S1B). While TERT is traditionally associated with telomere maintenance[68], its role in post-mitotic RPE and its functional relationship with AKT1 is poorly defined. We found that *Akt2* KI RPE, characterized by autophagic arrest, exhibited significantly reduced TERT expression and activity, whereas compensatory AKT1 expression in *akt2* cKO mice was associated with basal level TERT expression and telomerase activity (Fig. S3A-C). In human cells, AKT1 phosphorylates TERT at Ser227 and Ser824 to facilitate its nuclear import[44] (Fig. S3D). Pharmacological AKT2 inhibition (6 nM AKT2i) in iPSC-RPE *CFH*^Y402H^ induced TERT phosphorylation at Ser824 (Fig. S3E, F). Conversely, selective AKT1 inhibition suppressed phospho-TERT (Ser824) in hTERT RPE and triggered compensatory AKT2 upregulation (Fig. S3G–I), confirming the bidirectionality of this signaling axis. Notably, neither AKT1 nor AKT2 compensation altered telomere length in the RPE (Fig. S3J), suggestive that the AKT1-TERT axis serves non-canonical regulatory functions in the post-mitotic RPE rather than traditional telomere maintenance.

### AKT1 compensation assembles a nuclear FOXO3–MYC–TERT transcriptional scaffold

To define the kinetics of the AKT-TERT axis, we performed nuclear-cytoplasmic fractionation of *CFH*^Y402H^ iPSC-RPE. Post AKT2i treatment on iPSC RPE cells followed by nuclear-cytoplasmic fractionation, we found that the AKT2 inhibition triggered a robust, time-dependent increase in nuclear phospho-TERT (Ser824) that tracked with p-AKT1 (Ser473), peaking at 6 and 24 hours (Fig. 3A–D). Importantly, TERT translocated to the nucleus alongside FOXO3, a master regulator of the autophagy-lysosomal pathway[30, 31]. In *Akt2* KI RPE, FOXO3 was hyperphosphorylated at the cytoplasmic retention site (Thr32)[69], a state that prevents the induction of autophagy genes (Fig S3K, L). AKT2 inhibition dephosphorylated FOXO3 and its coordinate nuclear translocation with TERT (Fig. 3E, F). Notably, FOXO1 remained phosphorylated and cytoplasmic under these same conditions, demonstrating that AKT1 compensation achieves isoform-selective FOXO regulation (Fig. S3M, N). AKT1 drives assembly of a tripartite FOXO3-MYC-TERT nuclear complex, with MYC serving as the central hub that simultaneously bound both TERT and FOXO3 (Fig. 3G–J). We demonstrated that AKT1 coordinates a hierarchical feed-forward loop through this complex assembly, where FOXO3 binds and activates the *MYC* promoter (Fig. 3K, L) and MYC subsequently occupies the E-box of the *TERT* promoter, driving its transcription (Fig. 3M, N) as demonstrated by CUT&RUN qPCR. This recruitment was functionally specific as enrichment was completely abolished in RICTOR knockdown and TERT knockout (KO) cells, even under AKT1-overexpressing conditions (Fig. 3L, N). These results demonstrate that the AKT1-mTORC2 axis bypasses the mTORC1-mediated autophagic block by stabilizing a nuclear FOXO3–MYC–TERT complex, which sequentially activates the transcriptional programs required to maintain lysosomal function and autophagic process in the aging RPE.

**Figure 3.**
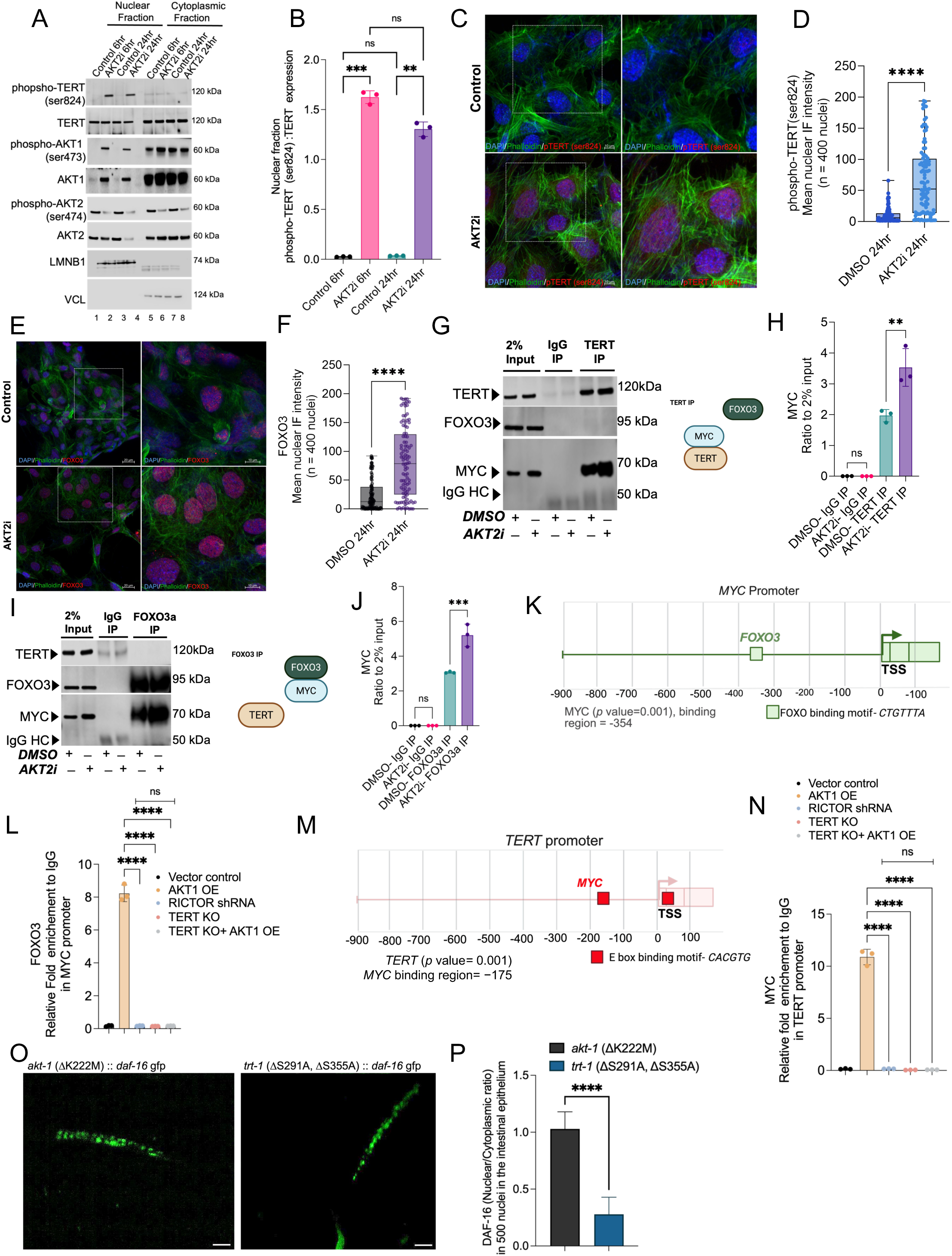
AKT2 inhibition promotes nuclear shuttling and assembly of a TERT-FOXO3-MYC transcriptional complex. (**A, B**) Subcellular fractionation and densitometry quantification in human iPSC-RPE showing time-dependent nuclear accumulation of phospho-TERT (Ser824) following AKT2 inhibition (AKT2i). LMNB1/Lamin B1 and VCL/Vinculin serve as nuclear and cytoplasmic markers, respectively. (**C**) Immunofluorescence showing phospho-TERT (ser824) activation and nuclear entry in *CFH* ^Y402H^ iPSC-RPE upon AKT2i treatment for 24hours. (**D**) Quantification of phospho-TERT nuclear intensity across 400 nuclei. (E, F) Representative immunofluorescence (IF) images (**E**) and quantification (**F**) of FOXO3 (red) nuclear intensity in *CFH* ^Y402H^ iPSC-RPE. AKT2i triggers significant nuclear translocation of FOXO3 (n = 400 nuclei). Nuclei were counterstained with DAPI (blue) and F-actin with phalloidin (green). (**G, H**) Co-immunoprecipitation (Co-IP) using anti-TERT in iPSC-RPE (**G**) and densitometry quantification (**H**). AKT2i enhances the physical association between TERT and MYC, but not FOXO3. Schematic (right) depicts the FOXO3-MYC-TERT assembly. (**I, J**) Reciprocal Co-IP using anti-FOXO3 (**I**) and densitometry quantification (**J**). FOXO3 interacts with MYC but not TERT, suggesting a bridged nuclear complex. Schematic (right) depicts the FOXO3-MYC-TERT assembly. (**K**) JASPAR analysis and showing FOXO3 enrichment at the MYC promoter consensus motif (−354 bp). (**L**) CUT&RUN qPCR validation showing AKT1 OE significantly enhances FOXO3 occupancy at the MYC locus AKT1 OE increases occupancy; enrichment is abolished in RICTOR shRNA, TERT KO, and TERT KO + AKT1 OE cells, confirming binding specificity. (**M**) JASPAR analysis showing MYC enrichment at the E box motif on TERT promoter (−175bp). (N) CUT & RUN qPCR shows that AKT1 OE increases MYC occupancy; enrichment is abolished in RICTOR shRNA, TERT KO and TERT KO + AKT1 OE cells, confirming binding specificity. (**O**) Volumetric analysis showing mean DAF-16/FOXO shuttling intensity of GFP tagged daf-16 in the intestinal epithelium of *akt-1* and *trt-1* variants across 57 slices P) Ratio of nuclear/ cytoplasmic DAF-16 GFP in *akt-1* and *trt-1* variant. For all graphs, data are represented as mean ± SD from at least 3 independent biological replicates. Statistical significance was determined by one-way ANOVA with Tukey’s post-hoc test. Scale bars: 10μm. **p < 0.01, ***p < 0.001, ****p < 0.0001. ns, non-significant

### The AKT1–TERT axis is an evolutionarily conserved regulator of lysosomal homeostasis

To determine whether the non-canonical proteostatic maintenance role of the AKT1–TERT axis is evolutionarily conserved, we utilized *C. elegans* models of post-mitotic cell survival[70]. We generated two specific knock-in strains: an *akt-1* kinase-dead mutant (ΔK222M; homologous to human AKT1 K179M)[71] and *trt-1* phosphorylation-deficient mutants (ΔS291A, ΔS355A; homologous to human TERT Ser227/Ser824 sites)[37] (Fig. S4A). In 15th-generation adults, these mutations drove reciprocal shifts in the autophagy-lysosomal transcriptome (Fig. S4B–E). While *akt-1* kinase-dead worms exhibited compensatory *akt-2* upregulation (mirroring our mammalian observations), the *trt-1* phosphorylation mutants displayed a catastrophic failure of mTORC1/C2 balance and autophagic machinery (Fig S4F–K) with significantly reduced expression of core autophagy genes *bec-1* (*BECN1*) and *atg-5* (*ATG5*) (Fig. S4I, J).

This transcriptional suppression was accompanied by a marked reduction in functional, acidified lysosomes in the intestinal epithelium as assessed by LysoTracker and LysoSensor imaging (Fig. S4L, M). The *trt-1* variants exhibited a dramatic decline in health span (Fig. S4N) and shortened lifespan (Fig. S4O, P). Mechanistically, this autophagic collapse was driven by the failure of DAF-16 (the *C. elegans* FOXO homolog) to undergo nuclear translocation[72]. DAF-16 nuclear/cytoplasmic shuttling assays confirmed that AKT1-mediated phosphorylation of TERT is required for DAF-16 nuclear intensity and transcriptional activity (Fig. 3O, P). Our findings demonstrate that the phosphorylation of TERT by AKT1 rather than telomerase activity, is a prerequisite for the nuclear recruitment of FOXO/DAF-16, thereby ensuring the maintenance of autophagic activity and lysosomal function across species.

### The FOXO3–MYC–TERT complex rewires the PERK–ATF4 axis to promote adaptive ERphagy and proteostatic resilience

Although sustained activation of the PERK–eIF2α–ATF4 arm of the UPR typically drives CHOP-mediated apoptosis[38], we found that AKT2 inhibition activates this pathway to promote cytoprotective proteostasis rather than cell death. Subcellular fractionation of AKT2i-treated *CFH*^Y402H^ iPSC-RPE revealed a time-dependent increase in nuclear ATF4 and p-eIF2α (Ser51), alongside elevated p-PERK (Thr980) and the chaperone BiP (Fig. 4A; Fig. S5A, B). Crucially, the pro-apoptotic factor CHOP was suppressed, while ERN1 activation and nuclear NRF2 levels were significantly increased (Fig. S5C–M), suggesting a shift from terminal UPR to an adaptive proteostatic state. This redirection is orchestrated by the physical engagement of the FOXO3–MYC–TERT complex with the ER stress machinery. Co-IP showed that TERT associates specifically with ATF4, while FOXO3 associates with both PERK and ATF4 (Fig. 4B, C), identifying FOXO3 as a stress responsive scaffold that bridges the nuclear complex to the ER-resident kinase. We confirmed the direct regulation of the UPR by the FOXO3–MYC–TERT complex through CUT&RUN-qPCR, which showed significant FOXO3 occupancy at the *EIF2AK3*/ PERK promoter (Fig. 4D, E). This enrichment was strictly dependent on mTORC2 integrity and TERT availability, as both RICTOR knockdown and TERT knockout completely abolished FOXO3 occupancy in *EIF2AK3* promoter despite AKT1 overexpression. This provides an explanation for how the nuclear transcriptional program retroactively tunes the activation of a cytoplasmic ER sensor.

**Figure 4.**
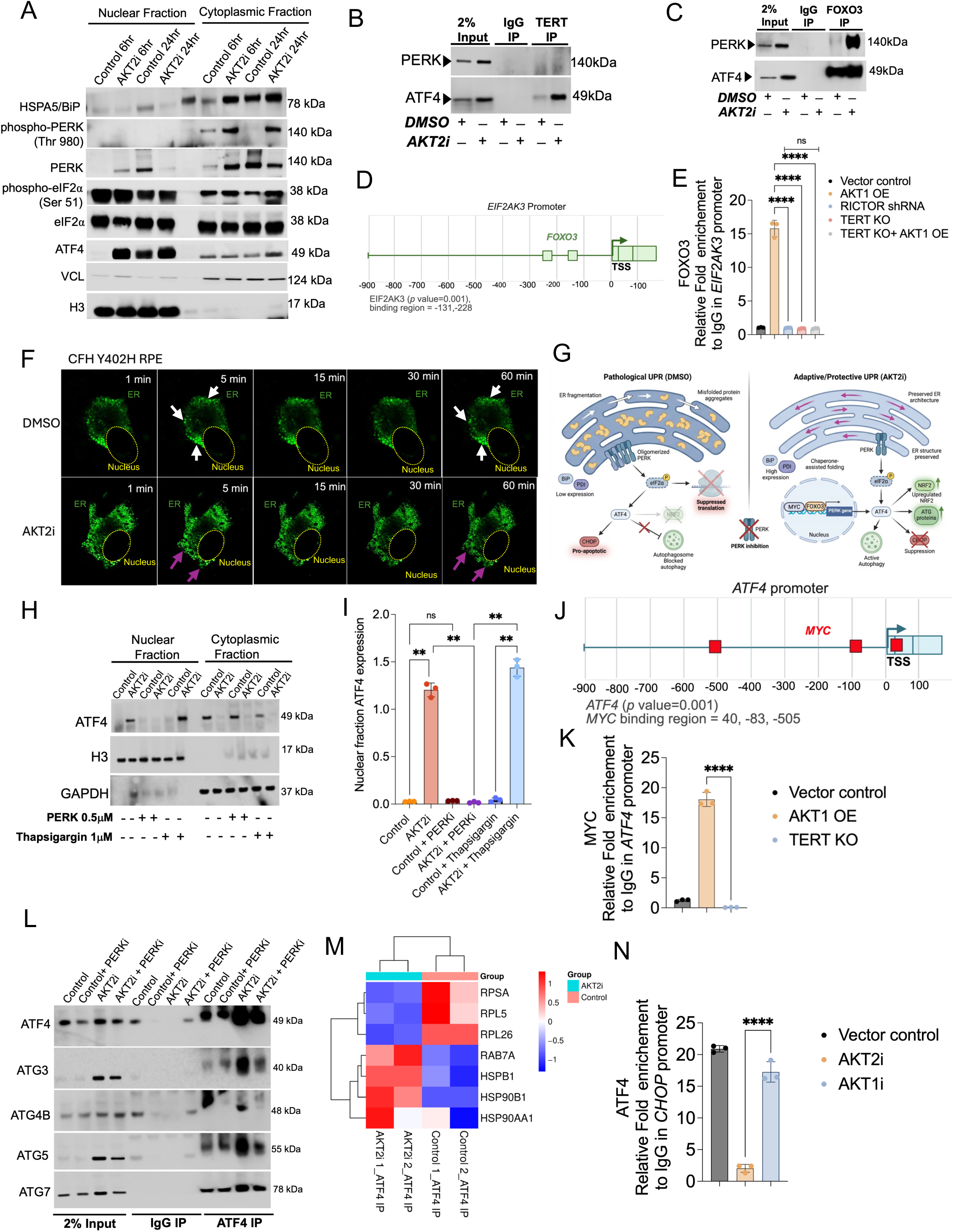
The AKT2-TERT-FOXO3 complex regulates the EIF2AK3-ATF4 stress response axis. (**A**) Subcellular fractionation and western blot analysis of human iPSC-RPE *CFH* ^Y402H^ cells treated with DMSO (control) or AKT2 inhibitor (AKT2i) for 6 and 24 h. AKT2 inhibition leads to a time-dependent increase in nuclear ATF4 and phosphorylated EIF2S1/eIF2α(Ser51), alongside increased levels of HSPA5/BiP and phospho-EIF2AK3/PERK (Thr980) in the cytoplasm. VCL and H3 serve as cytoplasmic and nuclear markers, respectively. (**B, C**) Co-immunoprecipitation (Co-IP) assays in RPE cells using TERT (**B**) or FOXO3 (**C**) antibodies. Western blots demonstrate increased physical association between TERT and ATF4, but not EIF2AK3/PERK, upon AKT2 inhibition in the TERT IP. Conversely, increased physical association is observed between FOXO3, EIF2AK3/PERK, and ATF4 in the FOXO3 IP, suggesting a stress-responsive protein scaffold. (**D**) JASPAR analysis and showing FOXO3 enrichment at the EIF2AK3 promoter consensus motif (−131 bp, −228 bp). (**E**) CUT&RUN qPCR validation showing AKT1 OE significantly enhances FOXO3 occupancy at the EIF2AK3 locus at -131bp. AKT1 OE increases occupancy; enrichment is abolished in RICTOR shRNA, TERT KO, and TERT KO + AKT1 OE cells, confirming binding specificity. (**F**) Live-cell imaging of PERK(LD)-HOTag3-EGFP in iPSC-RPE CFH Y402H cells. DMSO-treated cells show EIF2AK3 oligomerization and ER fragmentation (white arrows), indicating unfolded protein clusters. AKT2i prevents clustering, maintaining a tubular ER network and promoting the clearance of unfolded proteins (magenta arrows). Scale bar: 10μm. (**G**) Schematic showing pathological UPR and adaptive UPR mediated by PERK-eiF2α-ATF4. AKT2i treatment through increased FOXO3 occupancy at EIF2AK3/PERK promoter mediates an adaptive UPR for cell survival. (**H**) Western blot and densitometry quantification of nuclear ATF4 following treatment with AKT2i, the EIF2AK3/PERK inhibitor GSK2606414 (0.5 μM for 24 h), and the ER stress inducer thapsigargin (1 μM for 12 h). The nuclear translocation of ATF4 induced by AKT2i is significantly attenuated by EIF2AK3/PERK inhibition, identifying EIF2AK3 as a mandatory upstream mediator. (**I**) Quantification showing ATF4 expression in the nuclear fractions upon AKT2i and PERKi conditions. (**J**) JASPAR analysis and showing MYC binding sites at the promoter and TSS of ATF4 (40 bp, −83 bp, −505 bp). (**K**) CUT&RUN qPCR validation showing AKT1 OE significantly enhances MYC occupancy at the ATF4 locus at -83 bp. AKT1 OE increases occupancy; enrichment is abolished in TERT KO, confirming binding specificity. (**L**) Co-IP showing increased interaction of ATF4 with autophagy-related proteins (ATG3, ATG4B, ATG5, and ATG7) upon AKT2 inhibition, which is blocked by PERKi. (**M**) Co-IP/proteomics analysis reveals that AKT2i increases the physical interaction of ATF4 with molecular chaperones (HSP90B1, HSP90AA1) and the mitophagy regulator RAB7A, while decreasing interactions with ribosomal proteins (RPSA, RPL5, RPL26). (**N**) CUT&RUN qPCR shows significant reduction in ATF4 occupancy at the DDIT3 (CHOP) promoter at −505 bp from TSS, upon AKT2i. ATF4 occupancy at DDIT3 promoter was high in control and AKT1i conditions. Data represent mean ± SD from n = 3 independent experiments. Statistical significance was determined by one-way ANOVA; ****p < 0.0001. ns, non-significant.

The functional consequence of this rewiring was visualized via live-cell imaging of PERK(LD)-HOTag3-EGFP[73]. In the iPSC RPE from AMD donors, PERK spontaneously oligomerized into toxic clusters associated with ER fragmentation (white arrows, Fig. 4F), a hallmark of autophagic failure. Conversely, AKT2 inhibition prevented PERK clustering, maintained a tubular ER network and accelerated the clearance of unfolded protein aggregates (magenta arrows; Fig. 4F), consistent with productive ERphagy. The pathological versus adaptive UPR state, are illustrated schematically in Fig. 4G. PERK inhibition using GSK2606414, (PERK inhibitor -PERKi) completely abrogated the ATF4 nuclear translocation and adaptive program, even when the nuclear complex remained intact (Fig. 4H, I). These data establish PERK as the obligate relay between the FOXO3-MYC-TERT transcriptional complex and proteostatic output.

### TERT-scaffolded ATF4 physically recruits the core autophagy machinery

To map the genomic targets of the FOXO3–MYC–TERT complex downstream of PERK-ATF4[74], we mapped the genomic occupancy of its components using CUT&RUN-qPCR. We identified significant enrichment of MYC at the *ATF4* promoter specifically under AKT1 OE conditions (Fig. 4J, K) while, this recruitment was completely abrogated in TERT KO RPE cells, confirming TERT’s essential role as a non-canonical nuclear scaffold (Fig. 4K). Beyond transcription, we discovered that ATF4 facilitates a structural bridge to the autophagy machinery. ATF4-IP recovered core autophagic components, including ATG3, ATG4B, ATG5, and ATG7 (Fig. 4L). This physical interaction was abolished by PERK inhibition, placing assembly of the autophagy cascade directly downstream of the PERK-ATF4 relay.

Proteomic analysis of the ATF4 interactome confirmed a functional re-wiring of the RPE stress response toward selective clearance. AKT2 inhibition (AKT2i) shifted ATF4 association away from ribosomal proteins (RPSA, RPL5, RPL26)[75, 76], consistent with the attenuation of mTORC1 linked global translation and toward molecular chaperones (HSP90B1, HSP90AA1) and the endo-lysosomal GTPase RAB7a (Fig. 4M). Given that RAB7a is a master regulator of autophagosome-lysosome fusion[77], these data suggest that ATF4 physically directs misfolded cargo to the degradation machinery. ATF4 also directly coordinates *CHOP* transcription (Fig. 4N) identifying it as the critical scaffold that restores autophagic flux and proteostasis in the diseased RPE.

### Convergent ATF4 transcription and PRKAA1–ULK1 signaling drive dual-axis autophagy induction

While the ATF4-dependent upregulation of the autophagy machinery is necessary for proteostasis [78], it is often insufficient to drive flux without a proximal initiating signal. We found that AKT2 inhibition (AKT2i) in *CFH*^Y402H^ iPSC-RPE activated PRKAA1 (p-Thr172) and the downstream autophagy-initiating kinase ULK1 at its activating Ser555 site is time dependent [79] (Fig. 5A–C). Concurrently, the inhibitory mTORC1-dependent phosphorylation of ULK1 at Ser757[80] was reduced, generating a robust biochemical signal for autophagy initiation. This biochemical activation was matched by an increase in autophagic core machinery components, including ATG3, ATG4B, ATG7 and ATG13 [81](Fig. 5D–F). To determine if the FOXO3–MYC–TERT complex directly orchestrates the biogenesis of the autophagic machinery, we performed CUT&RUN-qPCR to map genomic occupancy at core *ATG4B* loci. We identified significant enrichment of both MYC and ATF4 at the *ATG4B* promoter specifically under AKT1 OE conditions (Fig. 5G-I). Crucially, this recruitment was completely abrogated in TERT KO RPE cells (Fig. 5H, I), confirming that TERT scaffold integrity is a prerequisite for efficient MYC and ATF4 chromatin recruitment in *ATG4B* promoter for autophagy induction.

**Figure 5.**
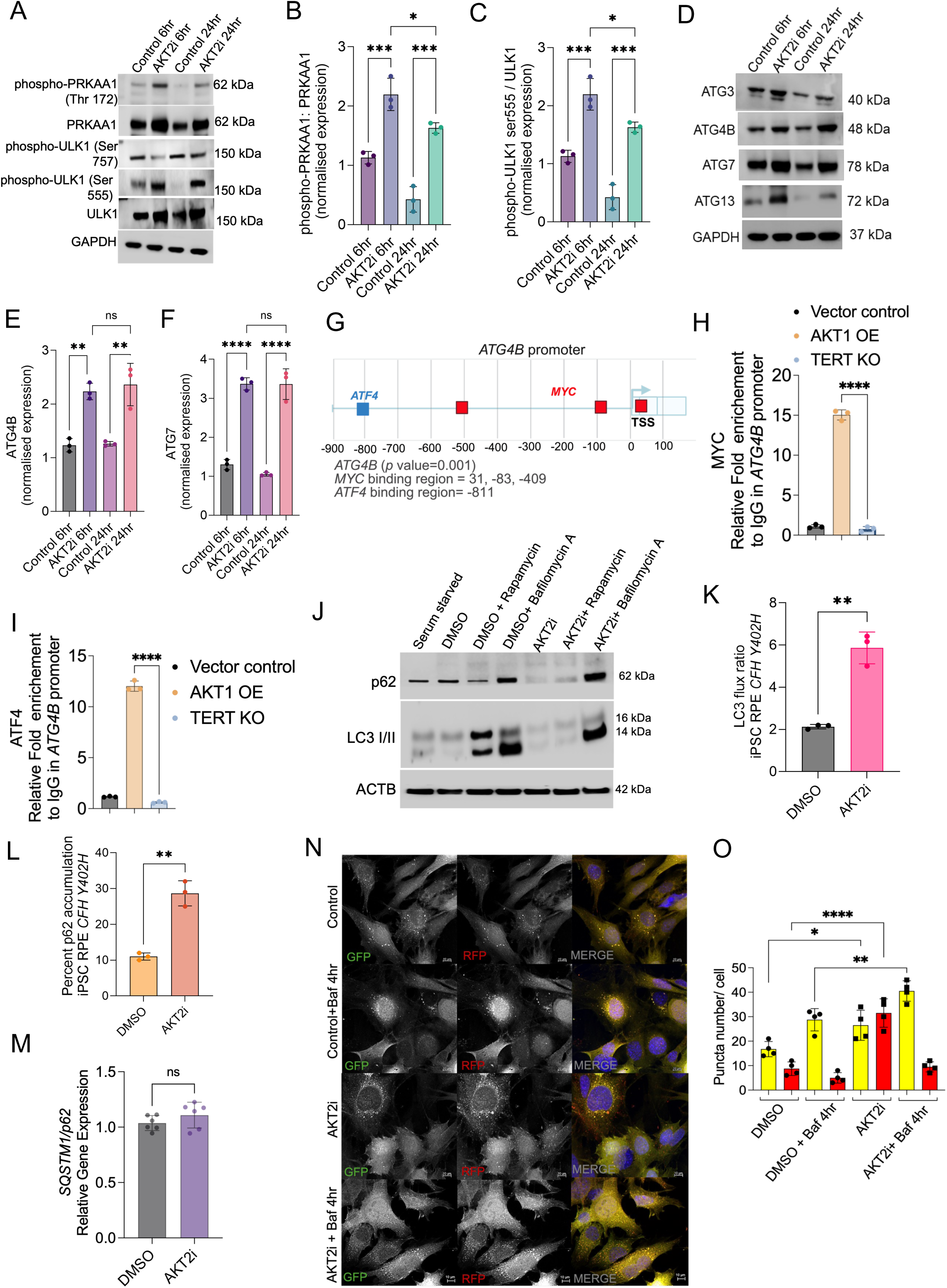
AKT2 inhibition rescues autophagic flux and mitigates proteotoxic stress via the PRKAA1-ULK1 axis. (**A**) Western blot analysis of human iPSC-RPE *CFH* ^Y402H^ cells treated with DMSO (control) or AKT2 inhibitor (AKT2i) for 6 and 24 h. AKT2 inhibition leads to the activation of the energy sensor PRKAA1/PRKAA1 (phospho-Thr172) and its downstream effector ULK1 (phospho-Ser555), while reducing the inhibitory phosphorylation of ULK1 at Ser757. (**B, C**) Densitometry quantification of (**B**) phospho-PRKAA1/PRKAA1α and (**C**) phospho-ULK1 (Ser555) in iPSC-RPE *CFH* ^Y402H^ cells treated with DMSO or AKT2i for 6 and 24 h; n = 3 independent biological replicates. (**D**) Western blot analysis of ATG proteins in iPSC-RPE *CFH* ^Y402H^ cells treated with DMSO (control) or AKT2 inhibitor (AKT2i) for 6 and 24 h. AKT2 inhibition leads to the activation of ATG3, ATG4B, ATG7 and ATG13 at 6 and 24 hours. (**E, F**) Densitometry quantification of (**E**) ATG4B and (**F**) ATG7 in iPSC-RPE *CFH* ^Y402H^ cells treated with DMSO or AKT2i for 6 and 24 h; n = 3 independent biological replicates. (**G**) JASPAR analysis and showing MYC (31 bp, −83 bp, −409 bp) and ATF4 (−811 bp) binding sites at the ATG4B promoter. (**H**) CUT&RUN qPCR shows significant enrichment of MYC at −83 bp site of the ATG4B promoter in AKT1 OE condition which was abolished in TERT KO condition. (**I**) CUT&RUN qPCR shows significant enrichment of ATF4 at −811 bp site of the ATG4B promoter in AKT1 OE condition which was abolished in TERT KO condition. (**J–L**) Representative western blots (**J**) and quantification of the MAP1LC3B/LC3-II flux ratio (**K**) and SQSTM1/p62 accumulation (**L**) in iPSC-RPE cells. Cells were treated with DMSO or AKT2i in the presence or absence of BafA1 or rapamycin. The significant increase in MAP1LC3B-II levels upon BafA1 treatment in AKT2i-treated cells confirms an increase in autophagic flux rather than a block in lysosomal degradation. (**M**) qPCR showing *SQSTM1* mRNA expression levels to distinguish between transcriptional induction and autophagic clearance. (**N**) Representative confocal images of human RPE cells expressing a tandem mRFP-GFP-MAP1LC3B reporter. Yellow puncta (GFP+RFP+) represent autophagosomes, while red-only puncta (GFP–/RFP+) represent acidic autolysosomes. (**O**) AKT2i treatment significantly increases the number of red puncta, and the addition of BafA1 reveals a robust increase in total autophagic vacuole formation compared to controls. Scale bar: 10 μm. Data represent mean ± SD from n = 3 independent experiments. Statistical significance was determined by one-way ANOVA with Tukey’s post-hoc test; *p < 0.05, **p < 0.01, *** <p < 0.001, ****p < 0.0001.

To confirm that this induction results in productive degradation rather than a block in clearance, we performed autophagic flux assays. The addition of Bafilomycin A1 (BafA1)[82] to AKT2i-treated cells produced significantly greater LC3-II accumulation than BafA1 alone, confirming accelerated flux (Fig. 5J, K). While SQSTM1/p62 protein levels increased upon BafA1 in AKT2i condition (Fig. 5L), *SQSTM1* qPCR confirmed autophagic clearance failure rather than transcriptional induction (Fig. 5M)[83]. The tandem mRFP-GFP-LC3B reporter provided direct visual confirmation [26]. AKT2i significantly increased red-only autolysosome puncta (GFP /RFP+), and BafA1 addition revealed robust accumulation of yellow autophagosomes (GFP+/RFP+), which confirms that autophagosomes form and fuse with lysosomes in a complete, productive cycle (Fig. 5N, O). These data, together with the ATF4 driven ERphagy receptor program, depicted schematically in Fig. S5N, demonstrate that AKT1-mTORC2 activation simultaneously supplies both the transcriptional licensing and the initiating biochemical signal required for dual axis proteostatic clearance.

### mTORC2-dependent organelle remodeling synchronizes ERphagy flux

Beyond non-selective macroautophagy, we investigated whether the damaged ER serves as a specific substrate for selective degradation via ERphagy. RNA sequencing of AKT2i-treated iPSC-RPE revealed the significant induction of the ERphagy receptors *TEX264* and *CCPG1*, alongside the ER membrane translocon component *SEC61G* (Fig. 6A)[45, 84]. CUT&RUN-qPCR revealed distinct regulatory requirements for each receptor. We found that ATF4 directly occupies the *CCPG1* promoter in AKT1 OE cells and this recruitment was completely abrogated upon *RICTOR* knockdown or *TERT* deletion (Fig. 6B, C), identifying *CCPG1* as a high-fidelity target of the mTORC2-AKT1-TERT nuclear scaffold. In contrast, ATF4 occupancy at the *TEX264* promoter was significantly lower and only marginally affected by *TERT* KO, with no dependency on RICTOR/mTORC2 (Fig. 6D, E). ATF4 knockout iPSC RPE *CFH* ^Y402H^ cells show significant loss of CCPG1, which was increased in AKT1 OE conditions (Fig 6F,G) These data indicate that while the AKT1-TERT-ATF4 axis is the primary driver of *CCPG1*-mediated ERphagy under UPR stress[85], while *TEX264* induction may be sustained through alternative signaling to maintain basal ERphagy[86], even when the primary survival axis is compromised.

**Figure 6.**
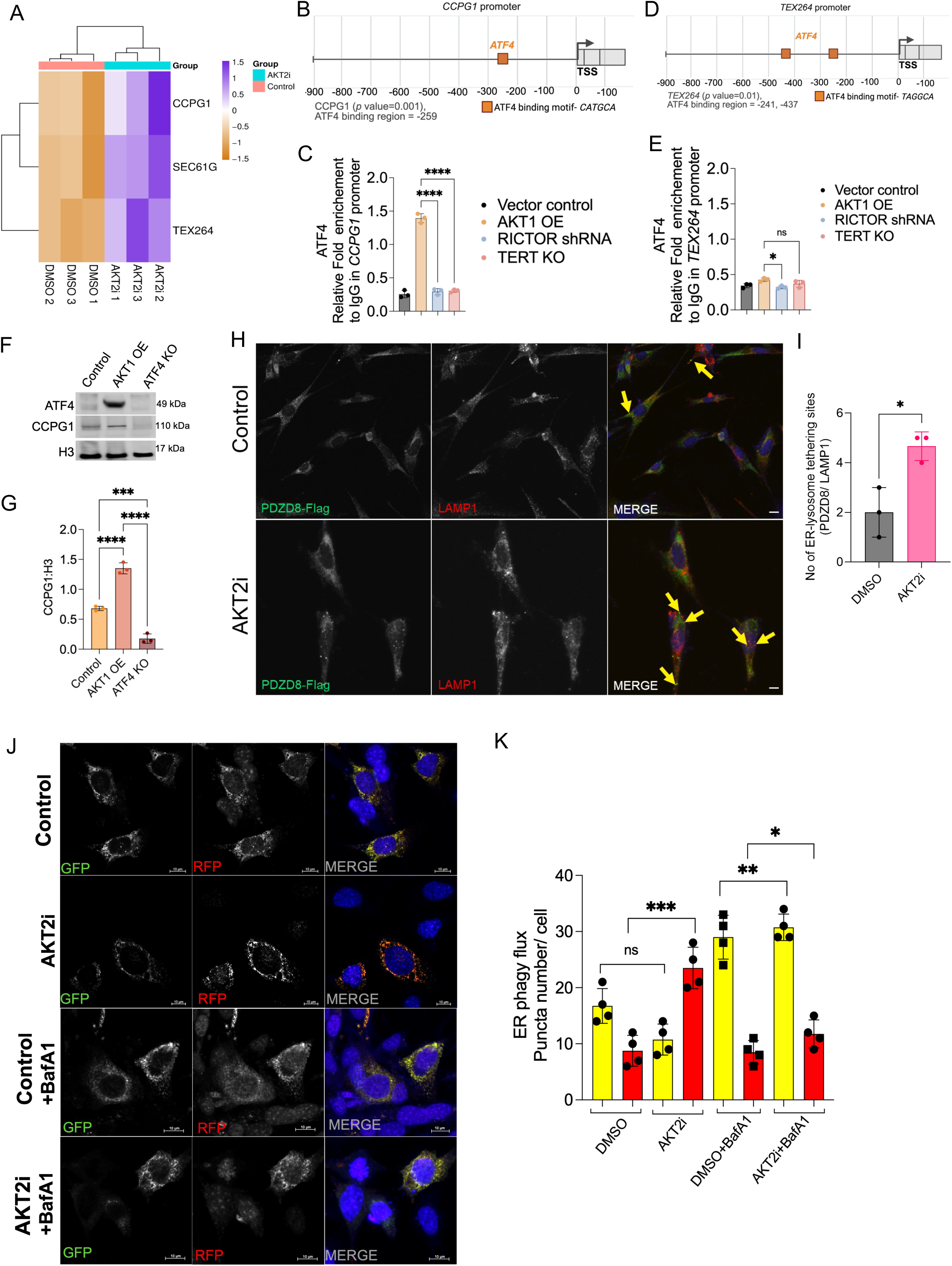
AKT2 inhibition drives ERphagy via ATF4-mediated induction of TEX264 and CCPG1. (**A**) Heatmap showing differential mRNA expression of ERphagy receptors (*CCPG1*, *SEC61G*, *TEX264*) in human iPSC-RPE *CFH* ^Y402H^ cells treated with DMSO or AKT2 inhibitor (AKT2i). AKT2 inhibition significantly upregulates receptors required for the selective clearance of the ER. (**B**) JASPAR analysis and showing ATF4 (−259 bp) binding site at CCPG1 promoter. (**C**) CUT&RUN qPCR shows significant enrichment of ATF4 at CCPG1 promoter at −259 bp upon AKT1 OE which was abolished in TERT KO and RICTOR shRNA conditions. (**D**) JASPAR analysis and showing ATF4 (−241, −437 bp) binding site at TEX264 promoter. (**E**) CUT&RUN qPCR analysis demonstrate weak occupancy of ATF4 in TEX264 promoter upon AKT1 OE, which shows significant reduction upon TERT KO but no significance with RICTOR knockdown condition. Representative confocal images (**F**) and quantification (**G**) of ER-lysosome tethering sites. RPE cells were transfected with PDZD8-FLAG (ER-lysosome tethering protein) and stained for LAMP1. AKT2i treatment significantly increases the number of PDZD8-positive LAMP1 contact sites (yellow arrows) per cell compared to DMSO controls. Scale bar: 10 μm. (**H, I**) Representative images (**H**) and quantification (**I**) of ERphagy flux using a tandem reporter KDEL GFP RFP system in RPE cells treated with DMSO or AKT2i and BafA1 conditions. Yellow puncta (GFP+RFP+) represent ERphagosomes, while red-only puncta (GFP–/RFP+) represent acidified ERphagolysosomes. AKT2i significantly enhances ERphagy flux. Data represent mean ± SD from n = 3 independent experiments. Statistical significance was determined by Student’s t-test or one-way ANOVA; *p < 0.05, **p < 0.01, ***p < 0.001.

Efficient ERphagy necessitates the physical orchestration of ER–lysosome contact sites along with receptor induction, to facilitate membrane engulfment[87, 88]. AKT2 inhibition significantly increased ER- lysosome contact site frequency, quantified by PDZD8-Flag/LAMP1 tethering[89] (Fig. 6H, I). This spatial remodeling was functionally linked to productive ERphagy flux, confirmed by a tandem GFP-RFP-KDEL ER reporter[90]. AKT2 inhibition significantly increased the accumulation of acidified, red-only ERphagolysosomes (GFP− / RFP+), indicating successful ER cargo delivery and degradation (Fig. 6J, K). Collectively, these results establish that the mTORC2-AKT1 driven FOXO3–MYC–TERT transcriptional complex coordinates both transcriptional and structural remodeling required for ERphagy flux, resolving the proteotoxic stress and organelle fragmentation that define RPE degeneration.

### An allosteric AKT2 inhibitor recapitulates the mTORC2–TERT–ERphagy cascade across disease models

Existing ATP-competitive AKT2 inhibitors such as CCT128930 achieve isoform selectivity only within a narrow dose window, limiting translational potential [50]. Structural analysis of the AKT2 DFG-out conformation identified two allosteric pockets at the pleckstrin homology–catalytic domain interface (pocket 1 near Trp80; pocket 2 near Tyr327) [91] (Fig. S6A, B). Trehalose [26,92] served as a thermodynamically validated docking template, and virtual screening of the dual-pocket pharmacophore identified 14 dual-pocket binders from 972 candidates, triaged by Blaze similarity, electrostatic complementarity, and docking score. In three independent *CFH*^Y402H^ iPSC-RPE lines, the resulting compound nAKT2i achieved sustained p-AKT2 (Ser474) suppression while consistently inducing compensatory p-AKT1 (Ser473) and p-TERT (Ser824) (Fig. 7A, B; Fig. S6C–E), providing a 20-fold broader therapeutic window than orthosteric inhibitors.

**Figure 7.**
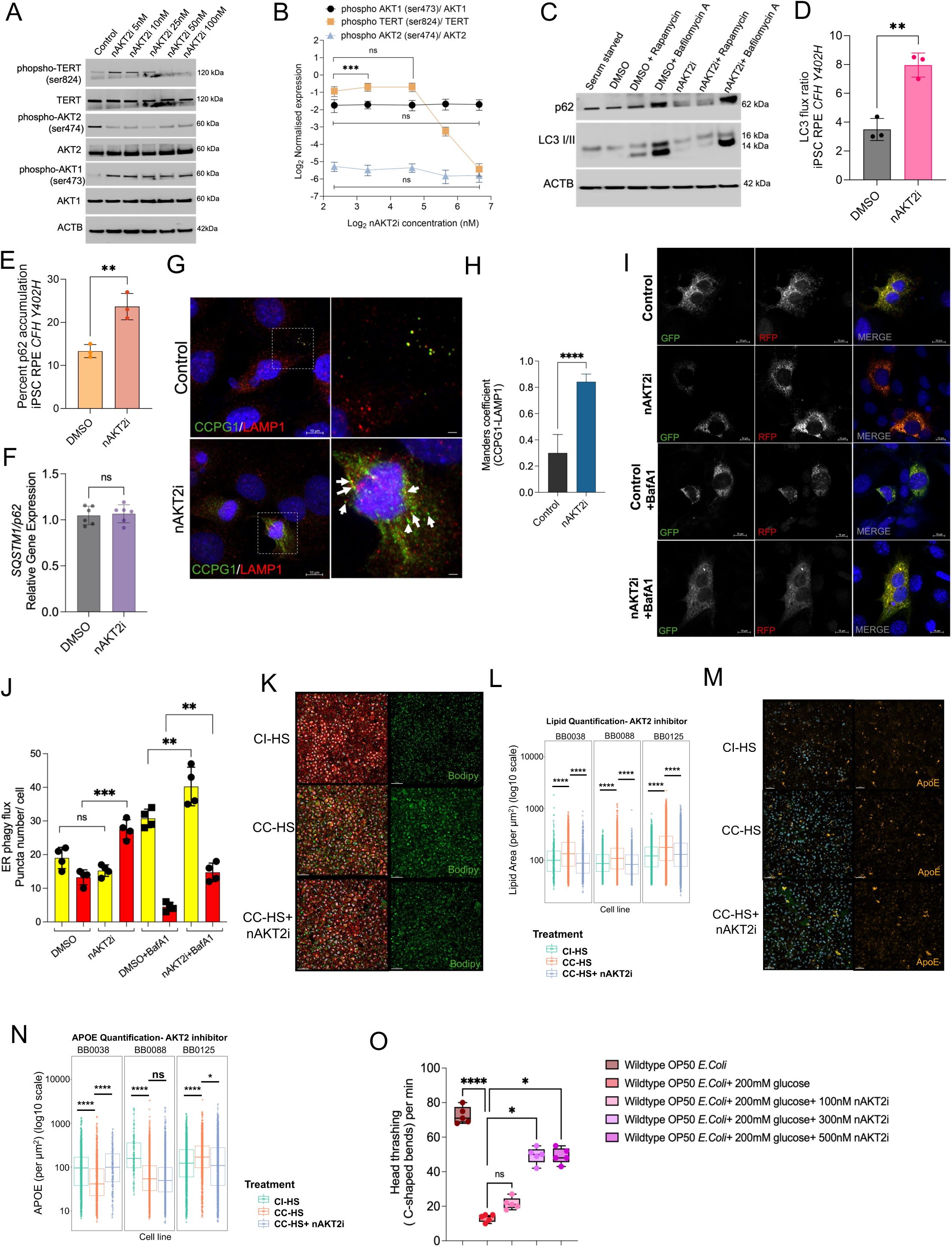
Selective AKT2 inhibition modulates mTORC2 signaling, rescues insulin sensitivity, and reduces lipid accumulation. (**A**) Dose-response analysis of a novel AKT2 inhibitor (nAKT2i) in three independent human iPSC-RPE *CFH* ^Y402H^ lines, showing sustained suppression of phospho-AKT2 (Ser474) over a broad concentration range (5–100 nM) for up to 48 h, with a compensatory upregulation of phospho-AKT1 (Ser473). (**B**) Line graphs showing densitometry analysis of phospho-AKT1 (Ser473), phospho-AKT2 (Ser474), and phospho-TERT (Ser824) levels in three independent iPSC-RPE lines upon nAKT2i treatment. (**C-E**) Representative western blots (**C**) and quantification of the MAP1LC3B/LC3-II flux ratio (**D**) and SQSTM1/p62 accumulation (**E**) in human iPSC-RPE cells. Cells were treated with DMSO or nAKT2i in the presence or absence of bafilomycin A1 (BafA1) or rapamycin. The significant increase in MAP1LC3B-II levels upon BafA1 treatment in nAKT2i-treated cells (relative to DMSO) confirms an increase in autophagic flux. (**F**) qPCR showing *SQSTM1* mRNA expression levels to distinguish between transcriptional induction and autophagic clearance. (**G, H**) Representative confocal images (**G**) and Manders’ co-localization coefficient analysis (**H**) of the ERphagy receptor CCPG1 and the lysosomal marker LAMP1 upon nAKT2i treatment. CCPG1/LAMP1 co-localization was increased upon nAKT2i treatment. (**I, J**) Representative images (**I**) and puncta quantification (**J**) of ERphagy flux in human RPE cells treated with nAKT2i, including BafA1 and rapamycin treatment panels. (**K**) Representative fluorescence images of human RPE cells stained with BODIPY (green) to visualize lipid droplets under CI-HS (control), CC-HS (AMD-risk), and CC-HS + nAKT2i conditions. (**L**) Quantification of lipid droplet area across three independent iPSC-RPE cell lines carrying the *CFH* Y402H polymorphism (BB0038, BB0088, BB0125). nAKT2i treatment significantly reduces the elevated lipid accumulation observed in CC-HS conditions. (**M, N**) Immunofluorescence imaging for APOE (orange) (**M**) and quantification (**N**) across three independent RPE cell lines carrying the *CFH* polymorphism (BB0038, BB0088, BB0125). (**O**) Head thrashing (C-shaped bend) assay performed in *C. elegans*. High glucose-fed worms exhibited a significant reduction in the frequency of C-shaped bends per minute compared with normally fed controls, indicating impaired muscle responsiveness. Treatment with 100 nM nAKT2i produced no significant change, whereas higher doses (300 nM and 500 nM) of nAKT2i significantly improved muscle responsiveness, as indicated by the increased frequency of C-shaped bends (n=5 per condition). Data represent mean ± SD from n = 3 individual biological replicates per group. Statistical significance was determined by two-way ANOVA with Tukey’s post-hoc test; *p < 0.05, ** p<0.01, ****p < 0.0001; ns, non-significant. Scale bars: 10 μm.

In a complement-competent human serum (CC-HS) disease-in-a-dish model of AMD [93], nAKT2i fully recapitulated the adaptive UPR signature and autophagic rescue program: PERK/ERN1 activation and BiP/PDI induction with CHOP suppression (Fig. S6F–K), clearance of drusen forming ApoE aggregates[91], and CCPG1 ERphagy receptor induction (Fig. S6L–O). Both macroautophagic and ERphagy flux were restored, confirmed by tandem reporter assays and CCPG1/LAMP1 co-localization showing productive lysosomal delivery and cargo acidification (Fig. 7C–J). These proteostatic improvements resulted in significant lipid droplet and disease-associated lipid mediators/transporter like APOE upon nAKT2i treatment across *in vitro* disease model generated from multiple donor iPSC-RPE lines[26, 92, 93] (Fig. 7K–N).

The metabolic and proteostatic utility of nAKT2i extended beyond the RPE, demonstrating tissue-agnostic AKT isoform rebalancing in hyperglycemic *C. elegans* and insulin-resistant skeletal muscle cells and (Fig. 7O; Fig. S6P). Therapeutic efficacy was validated *in vivo* using *Nuc1* rats (*Cryba1* mutation), a model of progressive retinal degeneration [94]: nAKT2i restored rhodopsin signal[94] and EBP50 apical polarity[95] to near wild-type levels in animals that showed loss of both parameters with vehicle treatment (Fig. S7Q). These findings demonstrate that a single allosteric compound can recapitulate the full mechanistic cascade — mTORC2 engagement, TERT nuclear entry, adaptive UPR, and ERphagy — to arrest proteostatic collapse and structural degeneration across the outer retina and insulin-sensitive tissues

## Discussion

Post-mitotic cells face a fundamental paradox. They must maintain proteostasis for decades under chronic degradative demand, yet the kinase pathways that sense stress canonically suppress the very autophagy machinery required to meet that demand[22]. This study resolves how cells escape this constraint. We show that AKT1, a canonical autophagy suppressor[96], can be repurposed as an autophagy activator through a transcriptional mechanism that bypasses direct phosphorylation of the autophagy machinery. When AKT2 activity declines, compensatory AKT1 activation via mTORC2 phosphorylates TERT at Serine 824 to drive its nuclear entry. In the nucleus, TERT does not act alone. It assembles with FOXO3 and MYC into a tripartite transcriptional complex that directly activates PERK, igniting a self-amplifying loop in which PERK–ATF4 signaling simultaneously drives biogenesis of core autophagy machinery and induces selective ERphagy through the receptors TEX264 and CCPG1[45, 85, 86]. These observations identify the FOXO3–MYC–TERT complex as the molecular switch that converts an isoform-specific kinase signal into a coordinated organelle quality control program.

A central question in AKT biology is how two closely related isoforms, sharing over 80% sequence identity, produce opposite effects on cell survival. Our data provide a structural answer. AKT2 and AKT1 partition into distinct mTOR complexes at different subcellular locations. AKT2 hyperactivation drives constitutive mTORC1 signaling at the lysosome, simultaneously destabilizing mTORC2 through inhibitory phosphorylation of RICTOR at Thr1135[60], which suppresses the PI3K–mTORC2 axis required for AKT1 Ser473 phosphorylation[97] and creates a feedforward loop of autophagic arrest. AKT2 also engages a second suppressive mechanism through S6K-mediated phosphorylation of IRS-1 at the inhibitory Ser639 site, generating insulin resistance that further blocks AKT1 activation[48]. These two complementary mechanisms — mTORC2 destabilization and IRS-1 inactivation — work together to ensure that AKT2 hyperactivation is dominant and self-reinforcing. Selective AKT2 inhibition dismantles both mechanisms simultaneously, disinhibiting IRS-1 and restoring mTORC2 integrity, which together enable robust compensatory AKT1 Ser473 phosphorylation. The spatial segregation of the two isoforms, confirmed by isoform-specific co-immunoprecipitation from plasma membrane and lysosomal fractions, and by FRET analysis showing exclusive AKT1–MAPKAP1 proximity, means that AKT2 inhibition does not simply reduce total AKT activity. Rather, it repositions the entire signaling equilibrium from a mTORC1-dominant, autophagic arrest state toward a mTORC2-dominant, pro-survival state.

The most unexpected finding of this study is the identification of TERT as the nuclear effector through which AKT1 activates transcription. TERT is well-established as a telomerase catalytic subunit. Its nuclear functions beyond telomere maintenance have been recognized but poorly defined. We show that AKT1-mediated phosphorylation of TERT at Ser824 drives nuclear translocation[44] without altering telomere length in post-mitotic RPE, confirming its telomere-independent function[37]. In the nucleus, TERT does not directly bind FOXO3. Instead, MYC serves as the organizing hub, binding both FOXO3 and TERT simultaneously to form the tripartite FOXO3–MYC–TERT complex. The assembly logic follows a strict hierarchy. Upon AKT2 inhibition, the dephosphorylation of FOXO3 permits its nuclear entry alongside TERT. FOXO3 then occupies the MYC promoter, driving MYC transcription[28]. MYC in turn, occupies the TERT E-box, completing a feed-forward loop that amplifies complex assembly[98]. Importantly, FOXO1 remains phosphorylated and cytoplasmic under these conditions, demonstrating isoform-selective FOXO regulation — AKT1 compensation specifically releases FOXO3 while retaining FOXO1 inhibition[99]. This selectivity is critical because it allows autophagy induction without the broad transcriptional changes that global FOXO activation would produce. The complex is functionally essential. RICTOR knockdown or TERT deletion abolishes MYC and ATF4 chromatin occupancy at autophagy gene promoters even when AKT1 is overexpressed, placing TERT scaffold integrity as a prerequisite for the entire downstream program.

A second paradox resolved by this study concerns the PERK–eIF2–ATF4 arm of the UPR[39]. Sustained PERK activation typically drives CHOP-mediated apoptosis[38]. Yet we find that AKT2 inhibition activates PERK in a manner that is cytoprotective rather than lethal. The distinction lies upstream. When the FOXO3–MYC–TERT complex occupies the EIF2AK3 promoter, it drives controlled, transcriptionally licensed PERK activation rather than the spontaneous, stress-overload oligomerization that accompanies proteotoxic failure. Live-cell imaging of the PERK(LD)-HOTag3-EGFP reporter[73] makes this contrast vivid. In diseased RPE without AKT2 inhibition, PERK spontaneously clusters into toxic foci associated with ER fragmentation. AKT2 inhibition prevents this clustering entirely, maintaining a tubular ER network and accelerating aggregate clearance. PERK inhibition abrogates the entire adaptive program — ATF4 fails to translocate, autophagy gene induction is lost and ERphagy is abolished — even when the nuclear complex remains intact. This places PERK as an obligate relay between the transcriptional complex and the proteostatic output. ATF4 then functions as a dual-output integrator. It physically associates with core autophagy proteins ATG3, ATG4B, ATG5 and ATG7, bridging transcriptional induction to direct machinery recruitment. Simultaneously, it drives expression of the ERphagy receptors CCPG1 and TEX264 through distinct regulatory mechanisms[45] — CCPG1 being strictly dependent on the TERT scaffold and mTORC2 integrity, TEX264 being partially independent, providing a fail-safe for basal ERphagy maintenance[86, 100].

The evolutionary data anchor the biological importance of this axis. *C. elegans* share no reliance on telomerase for post-mitotic cell maintenance[101], yet the functional consequences of disrupting this signaling hierarchy are conserved across hundreds of millions of years of evolution. Worms with catalytically inactive *akt-1* showed compensatory *akt-2* upregulation, enhanced lysosomal acidification, and 90% survival at 30 days compared with 71% for wild-type controls. In striking contrast, *trt-1* mutants lacking the AKT1 phosphorylation sites showed lysosomal dysfunction, reduced *bec-1* and *atg-5* expression, progressive health span decline and only 63% survival. The DAF-16 nuclear localization data close the mechanistic loop in the worm. Just as FOXO3 nuclear entry in human RPE depends on TERT availability, DAF-16 nuclear translocation in *C. elegans* requires AKT1-mediated TERT phosphorylation[72]. The pathway topology — AKT1 phosphorylates TERT, TERT bound MYC enables FOXO/DAF-16 nuclear function, FOXO drives lysosomal and autophagic gene expression — is thus an ancient and essential feature of post-mitotic cell survival. This conservation argues that the FOXO3–MYC–TERT axis is not a cell-type-specific adaptation but a fundamental principle of proteostatic regulation in non-dividing cells.

The therapeutic translation of these findings required overcoming a significant pharmacological obstacle. Existing ATP-competitive AKT2 inhibitors like CCT128930 achieve isoform selectivity only within a narrow 6 nM dose window[50], limiting their translational potential. We addressed this by targeting the AKT2 DFG-out conformation, which uniquely exposes a pleckstrin homology–catalytic domain interface absent in the active state. Structure-based virtual screening against two allosteric pockets at this interface, guided by trehalose as a thermodynamically validated template, identified a first-in-class dual-pocket allosteric inhibitor, nAKT2i, with selective AKT2 inhibition maintained across a 5–100 nM range for 48 hours — a 20-fold broader therapeutic window. In three independent *CFH*^Y402H^ iPSC-RPE lines[93], nAKT2i reproduced the full mechanistic cascade: AKT2 suppression, compensatory AKT1 and phospho-TERT induction, adaptive UPR activation with CHOP suppression and restoration of both macroautophagic and ERphagy flux. Under complement-competent human serum conditions mimicking the inflamed, aged subretinal environment of AMD[93], nAKT2i cleared ApoE aggregates[102], induced CCPG1 and normalized ER stress markers. The inhibitor also demonstrated tissue-agnostic activity, rebalancing AKT isoforms in insulin-resistant skeletal muscle cells and in hyperglycemic *C. elegans* and restoring rhodopsin signal and EBP50 apical polarity in *Nuc1* rats *in vivo*[103, 104]. This breadth of efficacy reflects the tissue-agnostic conservation of the underlying mechanism.

This work establishes several principles with implications that extend well beyond retinal disease. First, kinase isoforms that share a catalytic domain can drive opposite cellular fates through differential subcellular partitioning into distinct signaling complexes. Second, a kinase can activate autophagy through transcriptional licensing rather than direct phosphorylation of autophagy initiators, providing a mechanism for sustained, gene expression-based autophagy induction as distinct from rapid post-translational activation. Third, TERT functions as a nuclear scaffold in post-mitotic cells that is entirely independent of the canonical telomere maintenance[35, 36], organizing transcription factor complexes that govern organelle quality control. Fourth, the UPR can be transcriptionally pre-programmed to adopt a cytoprotective rather than apoptotic output, with the FOXO3–MYC–TERT complex serving as the upstream licensing signal. Post-mitotic cells in the brain, heart, kidney, and retina must maintain proteostasis across decades[105] and dysregulated AKT isoform balance is increasingly implicated in neurodegeneration, cardiac failure, and diabetic organ damage. The AKT1–TERT–PERK–ATF4 axis identified here provides a mechanistic blueprint for restoring organelle quality control in these settings. The rescue of complement-induced proteostatic damage in *CFH* ^Y402H^ cells is particularly relevant to diseases where innate immune dysregulation and autophagic failure intersect, including Alzheimer disease[106] and membranous nephropathy[107]. Isoform-selective rebalancing of AKT signaling — activating the beneficial arm rather than suppressing the pathway globally — thus represents a therapeutic strategy with broad applicability across the spectrum of age-related disease

## Materials and Methods

### Antibodies

The primary antibodies Akt1 (2938S), Akt2 (3063S), mTOR (2983S), RPTOR (2280S), RICTOR (2114S), PERK (5683T), p-PERK (3179S), eIF2α (9722S), p-eIF2α (9721S), ATF4 (11815S), BiP (3177T), CHOP (2895T), FOXO3 (12829S), p-FOXO3 (9464S), ACTB (4970S), VCL (13901S) and MYC (5605T) were purchased from Cell Signaling Technology, Inc. TERT (NB100-317) was purchased from Novus Biologicals. P-NRF2 (ab76026) and LAMP1 (ab25245) were purchased from Abcam Inc. PDI (SPA-890) was purchased from Stressgen Biotechnologies. CCPG1 (13861-1-AP) and Histone H3 (17168-1-AP) were purchased from Proteintech Group, Inc., whilepTERT-Ser824 (SAB4504295) was purchased from Sigma Aldrich. The HRP-based secondary antibodies used for western blots were purchased from KPL: anti-rabbit (074–1506) and anti-mouse (074–1806). The fluorescently labelled secondary antibodies used for immunocytochemistry in this study were Alexafluor 488 Donkey anti mouse IgG (A31572) and Alexafluor 555 Donkey anti rabbit IgG (A21202) (Invitrogen, Thermo Fisher Scientific).

### Animals

All mouse studies were performed in compliance with the institutional and national ethical guidelines and approved by the Institutional Animal Care and Use Committee (IACUC) of The Johns Hopkins University School of Medicine. The following transgenic and genetically modified lines were used in this study: *Cryba1* conditional knockout (*cryba1* cKO)[5], *Akt2* knock-in (*Akt2* KI)[108, 109], *Akt2* conditional knockout (*akt2* cKO)[109], and *mLST8* knock-in (*mlst8* KI) mice[110]. All lines were developed and maintained on a uniform C57BL/6J background under pathogen-free conditions. Animals were housed in ventilated cages at 22 ± 1 °C with 50–60 % relative humidity on a 12 h light/12 h dark cycle and provided standard chow and sterile filtered water *ad libitum*. Both male and female animals were used for experiments, and age-matched controls were included for all comparative analyses. All procedures including anesthesia, euthanasia, and enucleation were performed under aseptic conditions to minimize discomfort. RPE sheets or tissue lysates were collected immediately after euthanasia and processed for RNA, protein, or histological analyses following standardized protocols[108, 110].

### *C. elegans* generation and maintenance

*Caenorhabditis elegans* strains were maintained on nematode growth medium (NGM) agar plates seeded with *E. coli* OP50 at 25 °C using standard conditions. Mutant lines were generated by CRISPR–Cas9 mediated genome editing to target key phosphorylation-dependent sites conserved with human AKT1 and TERT. The *akt-1*(ΔK222M) kinase-dead mutant corresponds to human AKT1 K179M, and the *trt-1*(ΔS291A, ΔS355A) double mutant corresponds to human TERT S227A and S824A. Genome-edited lines were established and maintained in the laboratory, and experiments were performed on synchronized L4 or young-adult worms as indicated.

### *C. elegans* phenotypic and functional assays

Synchronized 15th-generation L4 adults (n = 3; approximately 100 worms per condition) were collected for RNA extraction and expression analysis. Lysosomal function was evaluated by dual staining with LysoTracker Red DND-99 and LysoSensor Green DND-189 (L7528 and L7535, Invitrogen, Thermo Fisher Scientific), followed by live imaging of the intestinal epithelium. Worms were mounted on agar pads and anesthetized in 0.1% levamisole and 0.01% tricane in 1X PBS and imaged using OlympusFV 3000 microscope, under identical acquisition settings for all genotypes. Fluorescence co-localization of LysoTracker and LysoSensor signals (yellow puncta) was quantified as a measure of acidic functional lysosomes. Health span was assessed by recording pharyngeal pumping rates on days 1, 7, and 15 of adulthood, and lifespan was monitored for up to 30 days.

### C. elegans whole organism imaging

Whole-organism fluorescent imaging was performed using a custom-built Mesoscopic Oblique Plane Microscope (Meso-OPM) system[111]. Prior to imaging, adult *C. elegans* were anesthetized and mounted in 2% agarose padded slides. Volumetric imaging was performed at 1 Hz temporal resolution, capturing the entire nervous system of *C. elegans* with high resolution (x-y-z: 1.2*1.6*2.2 µm). The Meso-OPM configuration achieves single objective light sheet microscopy optimized to flexibly interchange between imaging speed, resolution, and field of view. Meso-OPM then utilizes a remote focusing system to correct for the off-axis tilt and capture the final image. Here, the imaging set-up consisted of an excitation light sheet formed from a 488nm laser (20mW, Coherent OBIS) galvanometrically scanned across the lateral extent of the sample for fluorophore excitation. We utilized the highest resolution primary objective (OL1: Olympus 20X XLUMPlan FL N,1.0 NA) configuration to capture cellular features. This objective focused the oblique excitation sheet at the sample while simultaneously collecting the fluorescence emission. Two additional matched objectives (OL2 and OL3: Olympus 20X XLUMPlan FL N,1.0 NA) were used downstream for remote focusing and image formation. The fluorescence signal then passed through an emission filter (bandpass 535/70 nm) before being captured by a high-speed sCMOS camera (Teledyne Photometrics Kinetix22). The output volumes were pre-processed to correct for minor positional offsets during scanning, normalize signal intensity by scaling each frame relative to the smoother time-averaged mean, and generate maximum intensity projections for visualization and subsequent analysis.

### Human iPSC-derived RPE culture and maintenance

Human iPSC derived RPE lines were a kind gift from the Henderson Ocular Stem Cell Laboratory (Dr, Srinivasa Sripathi, Retina Foundation of the Southwest, Dallas, TX, USA). The panel included RPE derived from AMD donors carrying the *CFH* Y402H variant (n = 3) and age-matched non-AMD sibling controls (n = 3). The derivation of iPSC cells from human AMD and the sibling control donors and the differentiation of iPSC into RPE was performed as previously described[43]. The iPSCs for ‘disease in a dish model’ developed from donor cells containing *CFH* (Y/Y) for controls, and the risk allele *CFH* (H/H) for AMD were differentiated and maintained as described previously[112]. The iPSC lines were cultured on vitronectin-coated plates in E8 complete medium (A1517001, Thermo Fisher). The recombinant vitronectin (A14701SA, Thermo Fisher) was diluted in DPBS at the concentration of 1:200 to get the final concentration of 2.5mg/ml. The cells were passaged every 4–5 days at 70–80% confluency and were fed every 24 h. Cells were maintained under standard RPE culture conditions at 37°C with 5% CO . All procedures involving human iPSCs were conducted in compliance with institutional biosafety and ethical regulations.

### Human RPE and hTERT-immortalized RPE cell lines

Human RPE cells and human telomerase reverse transcriptase immortalized RPE (hTERT-RPE) cells were used for selected mechanistic and imaging assays, including isoform-specific AKT inhibition, tandem fluorescent autophagy reporter analysis, and phospho-TERT (Ser824) detection. Both cell types were maintained under standard RPE culture conditions at 37°C in a humidified atmosphere of 5% CO . hFRPE cells were cultured in MEMα media supplemented with 5% FBS, growth factors and antibiotics. hTERT RPE is cultured in DMEM F12 media supplemented with 5% FBS and 1% antibiotics (Penstrep).

### Computational identification of novel AKT2 inhibitor

We employed structure-guided virtual screening to identify selective AKT2 allosteric inhibitors. Comparative analysis of activated and inactive AKT2 catalytic domain structures (Asp-Phe-Gly motif in “DFG-in” and “DFG-out” conformations) revealed two druggable pockets at the catalytic–pleckstrin homology domain interface: pocket 1 adjacent to Trp80 and pocket 2 near Tyr327. Trehalose was docked into both sites, and top-scoring poses were evaluated by docking scores, glycosidic torsion angles, and hydrogen-bonding patterns. Selected poses were validated through 150 ns molecular dynamics simulations (AMBER ff14SB[113], OpenFF 2.2.0[114]; explicit solvent, 298 K, 1 bar, 0.150 M). Trajectory clustering and electrostatic complementarity analysis (EC > 0.25) identified distinct trehalose conformations for each pocket. These MD-validated geometries served as independent templates for Blaze screening of approximately 23 million compounds using the AKT2 structure as excluded volume. The top 10,000 hits per pocket underwent multi-criteria triage: field similarity (>0.6), electrostatic complementarity (EC > 0.25), docking score (<−7.8 kcal/mol), and preservation of pocket-specific interactions (pocket 1: Trp80, Leu215, Arg269, Asp293; pocket 2: Tyr18, Asn53, Asp324). Candidates were filtered for either trehalose-like properties (SlogP −3.4 to −1.4; TPSA 149–230 Å^2^; MW 290–390 Da) or drug-like properties (SlogP <3.5; TPSA 50–150 Å^2^; MW 250–410 Da). Chemical liabilities were excluded and chemotype diversity was enforced through ECFP6 clustering[115]. This process yielded 504 pocket 1 candidates, 468 pocket 2 candidates, and 14 predicted dual-pocket binders for experimental validation.

### Pharmacological treatments

Pharmacological treatments were performed on confluent RPE monolayers maintained under standard conditions. An ATP-competitive AKT2 inhibitor at a dose of 6 nM (CCT12890; S2635, Selleckchem) was used to induce compensatory AKT1 activation at 6 h and 24 h, while a novel AKT2 inhibitor (nAKT2i) was applied at 10 nM for selective and sustained AKT2 inhibition up to 48 h. For autophagy flux assays, 100 nM of bafilomycin A1 (S1413, Selleckchem) was applied for 4 h to the cells to inhibit autophagosome–lysosome fusion, and 1 µM of thapsigargin for 12 h (S7895, Selleckchem) was applied as an ER-stress inducer for ERphagy assessment. The PERK inhibitor GSK2606414 (S7307, Selleckchem) was used at 0.5 µM for 24 h to determine the role of the PERK–eIF2α–ATF4 pathway in ERphagy induction. Vehicle-treated controls (DMSO) were included for all conditions, and cells were processed for imaging, RNA, or protein analyses immediately after treatment.

### RNA sequencing (RNA-seq)

RNA sequencing was carried out on RPE cells isolated from *Cryba1* cKO, *Akt2* KI, *Akt2* cKO mice, and on human iPSC-derived RPE obtained from *CFH* ^Y402H^ AMD donors and age-matched sibling controls, both before and after AKT2 inhibitor treatment. For mouse studies, RPE tissue was collected at 5 and 10 months of age corresponding to early and advanced disease stages, and total RNA was extracted using standard procedures previously established in the lab[108–110]. For human studies, confluent cultures were harvested at baseline and following pharmacological treatment. RNA integrity was confirmed prior to sequencing. cDNA libraries were prepared and sent to be sequenced (Novogene, USA) to obtain high-depth transcriptome coverage, and differential expression analyses were performed between experimental and control groups. Data were used for pathway enrichment and gene-network analyses[108–110].

### Quantitative RT-PCR

Total RNA was extracted from *Cryba1* cKO mouse RPE samples and *C. elegans* using a RNeasy Mini Kit (74104, Qiagen) following manufacturer’s instructions. Complementary DNA (cDNA) was synthesized from equal RNA input using SuperScript VILO cDNA preparation kit (11754-050, Invitrogen), and quantitative PCR (RT-qPCR) was performed using gene-specific primers for target genes with *ACTB* as internal control. Relative expression levels were normalized to internal housekeeping controls, and fold changes were calculated using the ΔΔCt method [108–110]. The primer sequences for mouse qPCR and *C. elegans* qPCR are provided in Table 1.

### Western blotting

Western blotting was performed using previously published methods from our laboratory[108–110]. The cell and tissue lysates were prepared in 1× RIPA buffer (20–188, EMD Millipore) containing 0.1% of protease inhibitor cocktail (I3786, Sigma Aldrich) and 0.1% phosphatase inhibitor cocktail (P0044-5ML, Sigma Aldrich). Densitometry was performed to estimate the protein expression relative to the loading control (ACTB, VCL or H3, as applicable) using ImageJ software (National Institute of Health).

### Phospho-proteomic analysis

Phospho-proteomic profiling was performed on RPE isolated from *Cryba1* cKO and control mice at 8 and 24 months of age, corresponding to early and advanced disease stages. Phosphorylated proteins were compared between groups to identify stage-specific differences. Differentially regulated phospho-proteins included molecular chaperones associated with endoplasmic reticulum protein-folding and stress-response pathways, as described in the Results.

### Co-immunoprecipitation

Co-immunoprecipitation (Co-IP) was performed to assess interactions among TERT, FOXO3, MYC, and ATF4 in RPE cell lysates under control or AKT2-inhibited conditions. To further investigate signaling scaffold interactions, plasma membrane fractions were isolated for MAPKAP1 and Rictor IPs, while lysosomal fractions were purified to perform RPTOR IP. Samples were incubated with specific antibodies against TERT, FOXO3, MYC, ATF4, MAPKAP1, RICTOR or RPTOR and protein A/G magnetic beads under gentle agitation. Immunocomplexes were washed, eluted and analyzed by immunoblotting with reciprocal antibodies to confirm protein–protein interactions. Input lysates and specific sub-cellular fractions were run in parallel to verify expression levels and fractionation purity.

### ATF4 immunoprecipitation proteomics

ATF4 immunoprecipitation was performed from RPE cell lysates following AKT2 inhibition, and bound proteins were processed for on-bead digestion as described previously[116]. Briefly, beads were resuspended in 8 M urea, 100 mM Tris-HCl (pH 8.5), reduced with TCEP, alkylated with chloroacetamide, and digested overnight with Trypsin/Lys-C. Peptides were acidified, desalted on C18 columns, and analyzed by nano-LC–MS/MS on an EASY-nLC 1200 coupled to an Exploris 480 mass spectrometer with FAIMS pro interface (Thermo Fisher Scientific). Spectra were processed using Proteome Discoverer 2.5 and searched with SEQUEST HT against the *Homo sapiens* UniProt database, with standard tolerances and dynamic modifications for oxidation and phosphorylation. Peptide and protein false-discovery rates were controlled at 1% using Percolator, and quantification was based on peptide-spectrum match (PSM) counts[116].

### CUT& RUN

Targeted protein-DNA complexes were enriched using the <u>EpiNext™ CUT&RUN Fast Kit</u> (EpigenTek, P-2028) following the manufacturer’s instructions. Briefly, approximately 1x10^6^ cells were lysed to isolate nuclei, which were then incubated with 2 µg of and affinity beads for 90 minutes at room temperature. Chromatin fragmentation and targeted cleavage of unbound DNA were performed in situ using a unique nucleic acid cleavage enzyme mix and Nuclear Digestion Enhancer for 10 minutes. Following stringent magnetic washes, the protected DNA fragments were released via Proteinase K digestion at 60°C and purified using DNA binding beads. The resulting high-resolution DNA was eluted in Elution Buffer and quantified for downstream application. The primers used for CUT& RUN qPCR are provided in Table 2.

### Förster Resonance Energy Transfer (FRET) acceptor photobleaching analysis

FRET via acceptor photobleaching was performed using an Olympus FV4000 confocal microscope equipped with the bleaching control application. iPSC RPE cells were transfected with either GFP-AKT1 (donor) and mCherry-MAPKAP1 (acceptor) or RFP-AKT2 (donor) and Magenta-MAPKAP1 (acceptor). Control groups expressing only the donor fluorophore (GFP or RFP alone) or the donor in the presence of an unbleached acceptor were used to account for spectral bleed-through and incidental donor bleaching. For each measurement, a pre-bleach baseline of donor and acceptor intensities was established, followed by targeted photobleaching of the acceptor ROI for 30 s at 100% laser power (561 nm for mCherry; 594/633 nm for Magenta) to achieve >90% reduction in acceptor fluorescence. Immediate post-bleach images were captured to quantify donor de-quenching. FRET efficiency was calculated using the formula: *E* = 1 - (*Dpost* - *Dpre*) + (*Dpost*); D_post_= Donor fluorescence intensity after acceptor bleaching, D_pre_= Donor fluorescence intensity before acceptor bleaching. All data were background-corrected and normalized against donor-only controls to ensure the specificity of the detected protein-protein interaction.

### Quantitative- telomerase reverse transcriptase amplification protocol (Q-TRAP) Assay

The Q-TRAP assay was performed according to established protocol[117]. Briefly, cells are harvested and lysed to obtain a protein extract containing telomerase. The cell lysate is then mixed with TRAP buffer, deoxynucleotide triphosphates (dNTPs), a telomerase RNA template, and specific PCR primers, followed by a 30-minute incubation at 30°C to allow telomerase to synthesize telomeric repeats. The resulting products are subjected to qPCR, where fluorescence is monitored in real-time to quantify the amplification of the telomeric repeats. Finally, qPCR results are analyzed by comparing the threshold cycle (CT) values against standard curves or controls, providing a precise quantification of telomerase activity in the biological samples.

### Relative telomere length measurement

Relative telomere length was measured in iPSC RPE cells using Relative human telomere length quantification qPCR assay kit (RHTLQ 8908, ScienCell). Briefly, genomic DNA was isolated from human RPE cells using an Qiagen DNA extraction kit, ensuring a concentration of 20-100 ng/µL. The reaction mix is then prepared following the manufacturer’s instructions, combining the qPCR master mix, telomere primer mix, reference gene primer mix, and the diluted genomic DNA template. This prepared mix was dispensed into qPCR plates or tubes, incorporating appropriate controls such as no-template controls (NTC) and positive controls if provided. qPCR was performed with initial denaturation at 95°C for 2-10 min, followed by 40 cycles consisting of denaturation at 95°C for 15-30 s, annealing at 60°C for 30 s, and extension at 72°C for 30 s. After amplification, the qPCR results are analyzed calculating relative telomere length by comparing Ct values, with the formula: Relative Telomere Length = (Telomere Gene Ct - Reference Gene Ct).

### Autophagy flux and ERphagy assessment

Autophagic flux was assessed in human RPE and iPSC RPE cells transduced with a tandem Adenovirus–GFP–RFP–LC3 reporter (2001, Vector Biolabs), which distinguishes autophagosomes (yellow puncta) from autolysosomes (red puncta). Cells were treated with either DMSO (control) or AKT2 inhibitor, in the presence or absence of bafilomycin A1 (100 nM, 4 h), to block autophagosome–lysosome fusion and quantify flux. ERphagy was independently evaluated using a tandem GFP–RFP–KDEL reporter (2001, Vector Biolabs) transfected into hFRPE cells with Lipofectamine 3000 (L3000015, Thermo Fisher Scientific). Following 48 h of expression, cells were treated with DMSO or AKT2 inhibitor to monitor ERphagy progression under basal and inhibited conditions. Confocal images were acquired using identical laser and detector settings, and puncta were quantified in *ImageJ* under uniform thresholds to determine puncta number and fluorescence intensity per cell.

### Assessment of ER - lysosome tethering

ER–lysosome contact sites[89] were visualized to evaluate changes in organelle tethering following AKT2 inhibition. hFRPE cells were transfected with the pCDH–PDZD8–FLAG[118] construct (a kind gift from Xiaojun Tan’s Lab, University of Pittsburgh, Pittsburgh, USA) using Lipofectamine 3000 (L3000015, Thermo Fisher Scientific) according to the manufacturer’s instructions. After 48 h of expression, cells were treated with or without the novel AKT2 inhibitor. PDZD8 localization was analyzed by immunofluorescence alongside the lysosomal marker LAMP1 to identify ER–lysosome contact sites. Confocal images were acquired under identical imaging parameters, and the number of PDZD8–LAMP1 juxtaposed puncta per cell was quantified in ImageJ to assess the extent of ER–lysosome tethering.

### Immunofluorescence and confocal imaging

Cells were fixed, permeabilized, and blocked with serum to prevent non-specific antibody binding, followed by incubation with primary antibodies against phospho-TERT and phospho-FOXO3 to assess nuclear localization, phospho-NRF2 to evaluate ERphagy induction, LAMP1 for lysosomal labeling in the pCDH-PDZD8–FLAG[118] transfection assay to examine ER–lysosome tethering, and LAMP1 and CCPG1 to analyze ERphagy regulation in the presence or absence of AKT2 inhibitor. After primary antibody incubation, cells were treated with Alexa Fluor–conjugated secondary antibodies and Alexa Fluor 488–phalloidin (A12379, Thermo Fisher Scientific) to visualize F-actin. Nuclei were counterstained with DAPI (0100-20, Southern Biotech). Confocal images for immunofluorescence, autophagy, and ERphagy flux assays were acquired using Zeiss LSM 710 2-Photon or Zeiss LSM 880 Axio Examiner microscopes (Zeiss, Switzerland) under identical laser power, gain, and acquisition parameters across all experimental groups. Nuclear translocation of TERT and FOXO3, as well as fluorescence puncta number and intensity, were quantified from optical sections using ImageJ (NIH) with uniform threshold settings to ensure consistent analysis across conditions.

### Statistical Analysis

All statistical analyses were performed as described in the corresponding figure legends. Biological replicates (*n*) are indicated for each experiment. Quantitative data are presented as mean ± SD unless otherwise specified. Statistical significance was determined using two-tailed Student’s *t*-test or one-way ANOVA with appropriate post hoc tests, as indicated in the figure legends. *P* < 0.05 was considered significant.

## Data availability

All data supporting the findings of this study are available within the paper and its Supplementary Information files. The authors will upload RNA sequencing and proteomic data to the GEO NCBI database after the acceptance of the manuscript. Source data for all main and extended figures are provided with this paper. The schematic in Fig.1H, Fig.4G and Fig. S5N is original and were created at BioRender.com and its raw illustration is available at; https://app.biorender.com/illustrations/canvas-beta/69c459fc1842cfa4965b3cb8 (Fig. 1H), https://app.biorender.com/illustrations/canvas-beta/69dd3e6eb65778fe97aa7ce6 (Fig. 4G), https://app.biorender.com/illustrations/canvas-beta/69dda45581dc2a2f5dd69e97 (Fig. S5N).

## Generative AI tools

The authors used Claude Sonnet 4.6 (Anthropic, claude.ai Pro) to assist with language editing and minor text corrections during manuscript preparation. The AI tool was not used to generate, analyze, or interpret any scientific data, and all intellectual content, conclusions, and original writing are solely the work of the authors.

## Acknowledgements

The authors would like to thank J. Samuel Zigler, Jr. for his help during the preparation of the manuscript. This study was supported by NIH 1R01EY031594-01A1 (DS&JTH), Edward N. & Della L. Thome Memorial Foundation Awards Program in Age-Related Macular Degeneration (DS), Maryland Stem Cell Research Fund-Launch Program (DS), Foundation Fighting Blindness- Free Family AMD Research Award (DS), The Johns Hopkins University School of Medicine start-up funds (DS), funds from the Frieda Derdeyn Bambas Professorship in Ophthalmology (DS), NIH K99EY033421 (SG), 5R01EY035412-02 (PPP), P30EY001765 core award from the National Eye Institute, NIH to the Wilmer Eye Institute, The Johns Hopkins University School of Medicine, and unrestricted funds from Research to Prevent Blindness Inc., NY to the Wilmer Eye Institute, The Johns Hopkins University School of Medicine. We acknowledge Robert Scoffin and Krishnakumar Muraleedharan at Cresset for coordinating the inhibitor discovery services provided to the Sinha Laboratory under a paid service agreement, Naresh Rajendran, Adnin Ashrafi for iPSCs, and Avinash Soundararajan, Emma H Doud, Whitney Smith-Kinnaman, and Amber Mosely for the mass spectrometry work, which was done by the Indiana University School of Medicine Center for Proteome Analysis. Acquisition of the IUSM Center for Proteome Analysis instrumentation used for this project was provided by the Indiana University Precision Health Initiative. The proteomics work was supported, in part, by the Indiana Clinical and Translational Sciences Institute (Award Number UL1TR002529 from the National Institutes of Health, National Center for Advancing Translational Sciences, Clinical and Translational Sciences Award) and the P30 IU Simon Comprehensive Cancer Center Support Grant (Award Number P30CA082709 from the National Cancer Institute). We thank Xiaojun Tan for providing the pCDH-PDZD8-flag and pCDH-PDZD8-EGFP plasmids.

## Author Contributions

D.S. conceived, designed, and supervised the study and secured funding. V.S.B. performed most experiments and data analyses. S.G., S.B., and S.H. assisted with cell culture, immunohistochemistry, and biochemical assays, and provided technical support for multiple experiments. R.J., R.S., and K.B. generated iPSC-derived RPE cell lines from non-AMD donors, including CFH-null and isogenic control iRPE cells for the rescue experiments. D.M. and J.Y. performed high-resolution 3D imaging of *C. elegans*. P.P. contributed to mass spectrometry studies. M.J.S., S.L., N.F.B., and N.J.K. contributed to experimental design, molecular modeling, virtual screening, and manuscript preparation for the structure-guided discovery of a selective AKT2 inhibitor. S.S. generated iPSC-derived RPE cell lines from human AMD donors. J.A.S., A.G. J.T.H and K.R. contributed to experimental design and data interpretation. D.S., V.S.B., S.G., S.B., S.H and J.T.H wrote the manuscript. All authors contributed to data interpretation, reviewed and edited the manuscript, and approved the final version.

## Abbreviations

ACTB: actin, beta
AKT1: AKT serine/threonine kinase 1
AKT2: AKT serine/threonine kinase 2
ALP: autophagy-lysosomal pathway
AMD: age-related macular degeneration
ATF4: activating transcription factor 4
ATG3: autophagy related 3
ATG4B: autophagy related 4B cysteine peptidase
ATG7: autophagy related 7
CCPG1: cell cycle progression 1
CFH: Complement factor H
C57BL/6J: C57 black 6J
DMSO: dimethyl sulfoxide
MYC: cellular myelocytomatosis oncogene
EIF2A/eIF2: αeukaryotic translation initiation factor 2 subunit alpha
ER: endoplasmic reticulum
ERphagy: endoplasmic reticulum-selective autophagy
FOXO1: forkhead box O1
FOXO3: forkhead box O3
iPSC-RPE: induced pluripotent stem cell-derived retinal pigment epithelium
IRE-1α: Inositol-requiring enzyme 1α
ITGB1/integrin β1: integrin subunit beta 1
MAP1LC3/LC3: microtubule associated protein 1 light chain 3
mTOR: mechanistic target of rapamycin kinase
mTORC1: mTOR complex 1
mTORC2: mTOR complex 2
PBS: phosphate-buffered saline
PERK/EIF2AK3: eukaryotic translation initiation factor 2 alpha kinase 3
PI3K: phosphoinositide 3-kinase
RICTOR: Rapamycin-insensitive companion of mammalian target of rapamycin
RPE: retinal pigment epithelium
RPTOR: Regulatory-associated protein of mTOR
MAPKAP1: Sty1/Spc1-interacting protein1
SQSTM1/p62: sequestosome 1
TERT: telomerase reverse transcriptase
TEX264: testis expressed 264, autophagy receptor
UPR: unfolded protein response
VCL: vinculin
WT: wild type

