## Supplementary Figures, Tables and Legends for "AKT1-phosphorylated TERT assembles a FOXO3–MYC transcriptional complex that drives ERphagy and proteostasis in post-mitotic RPE"

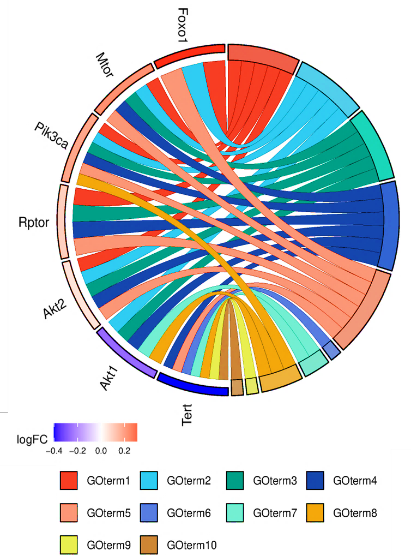

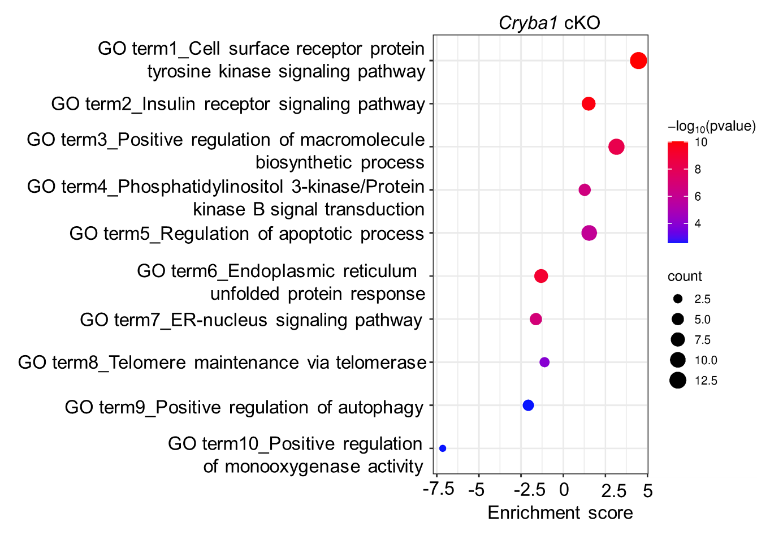

A

B

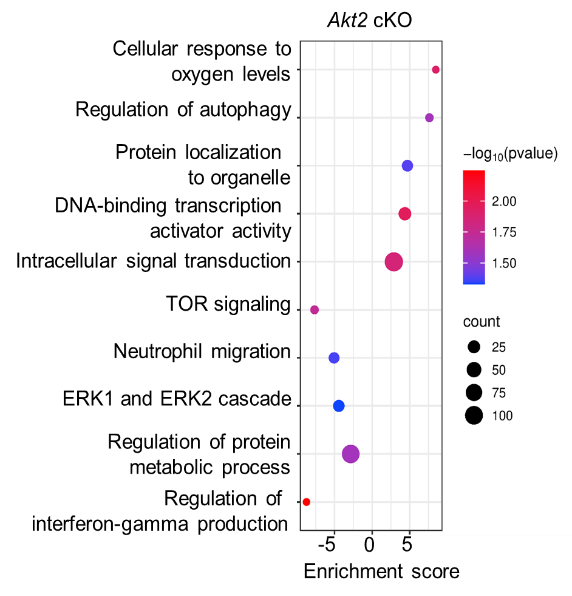

C

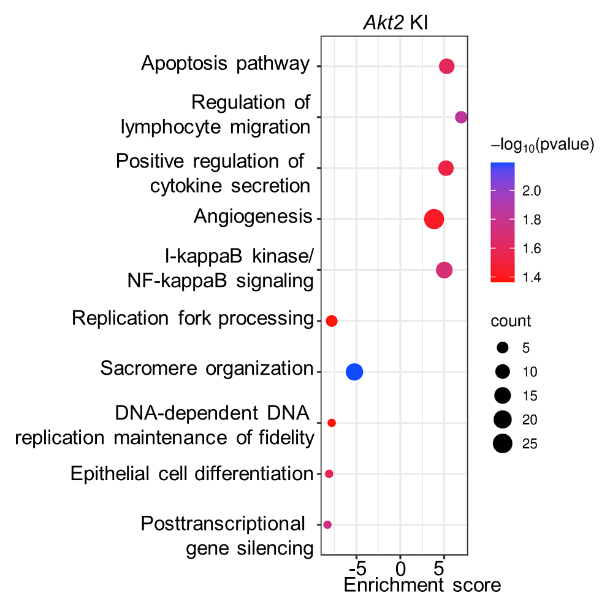

E

F

G

D

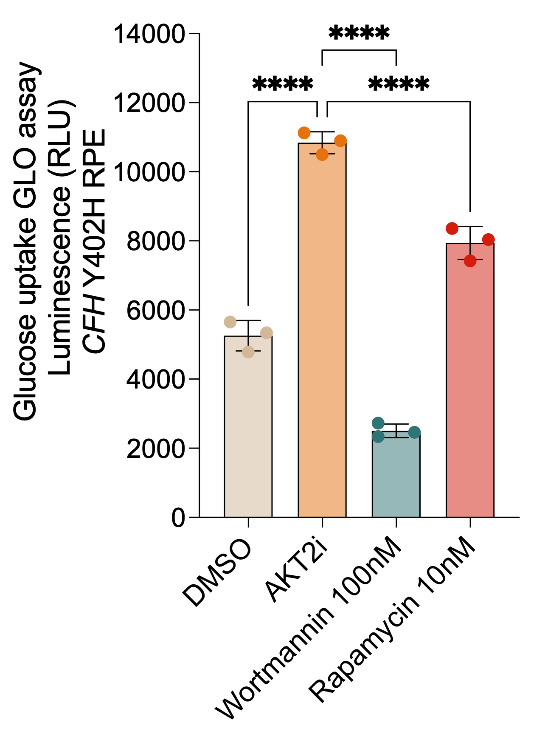

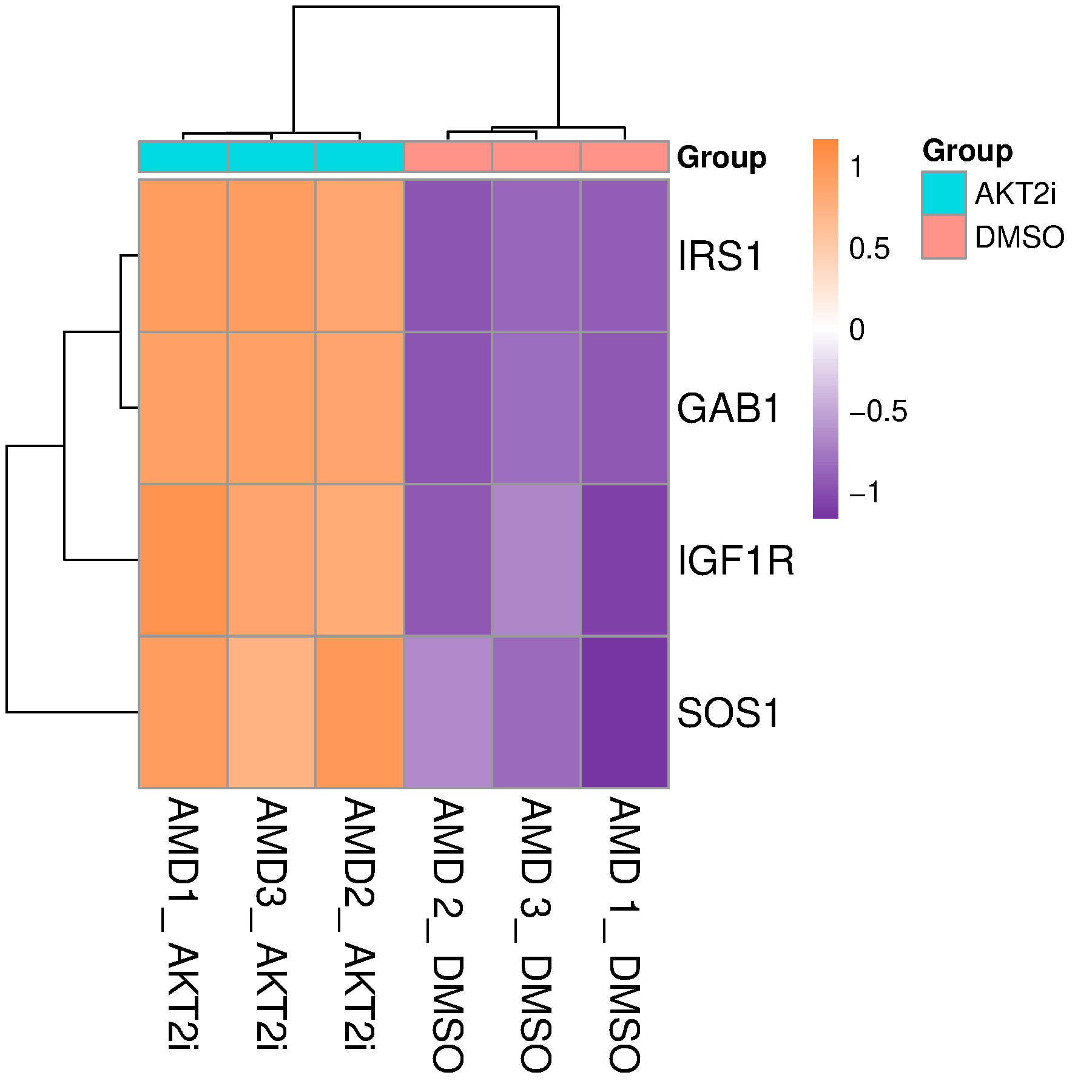

H

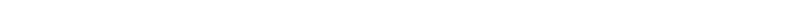

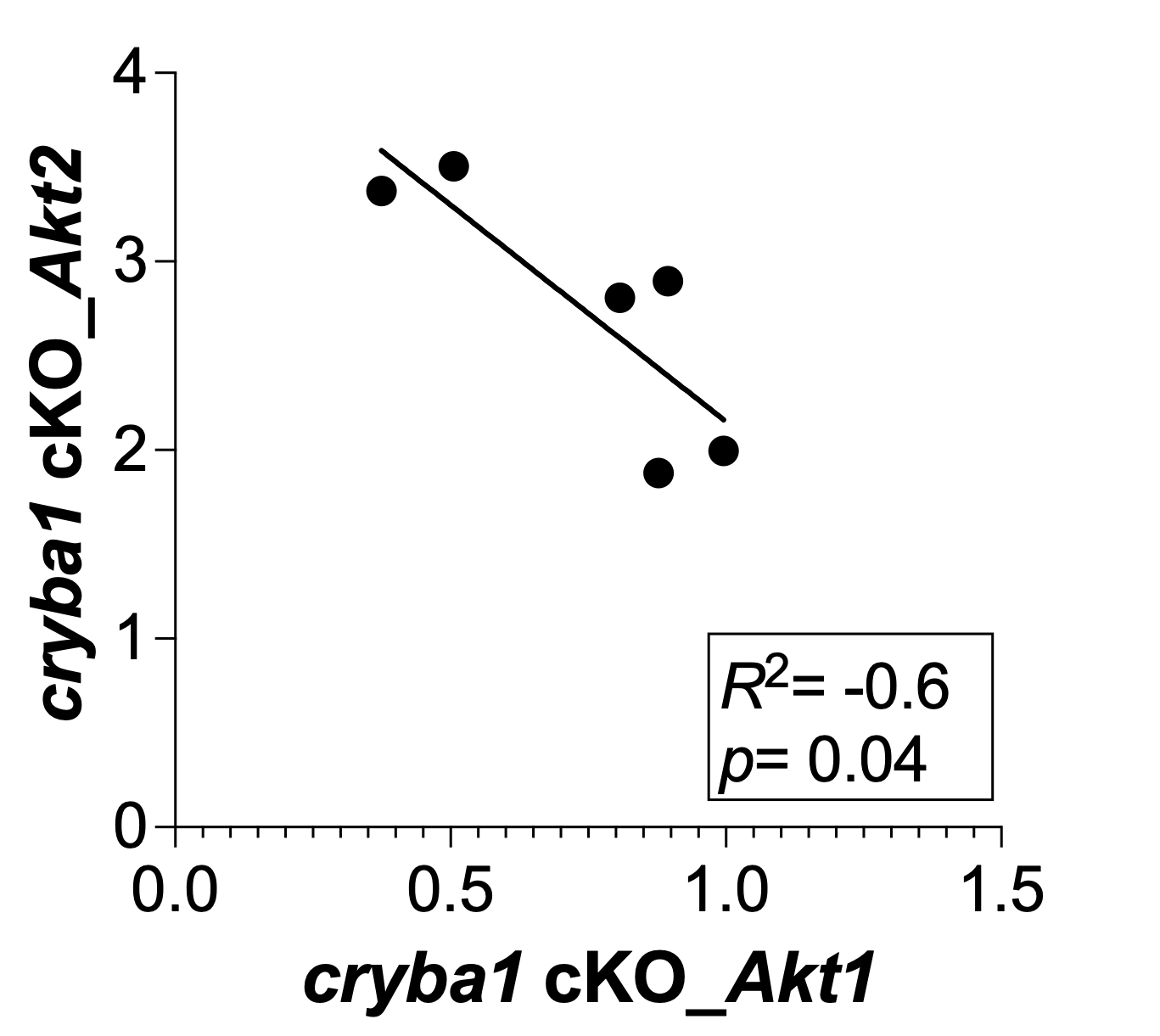

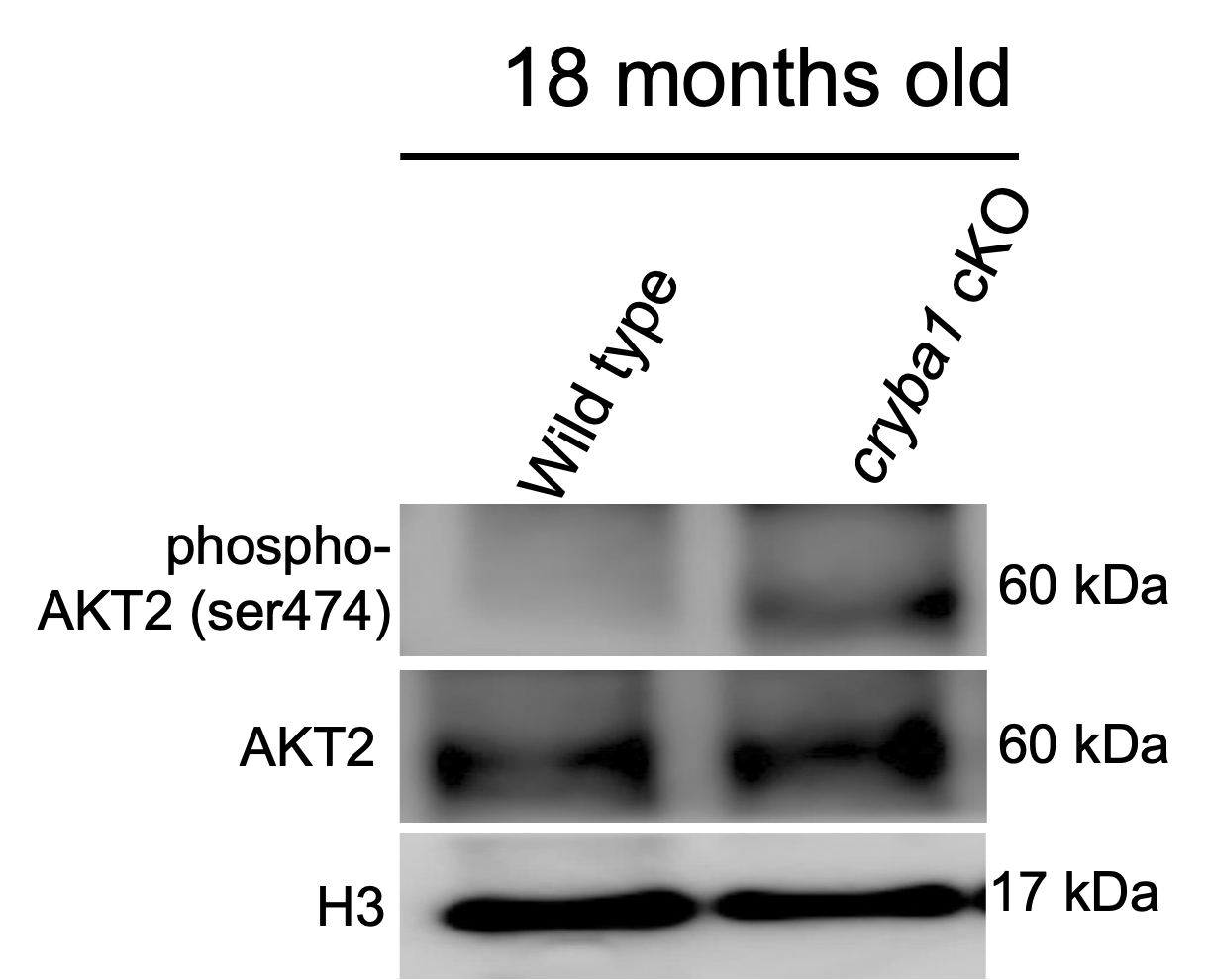

**Figure S1. Reciprocal regulation of AKT1 and AKT2 defines aging pathways in the RPE.** (**A**) Dot plot showing enriched Gene Ontology (GO) terms for differentially expressed genes in *cryba1* cKO RPE. The y-axis represents biological processes; dot size indicates gene count and color represents the -log10 p value. (**B**) Circos plot illustrating the overlap between key metabolic genes (left) and their associated functional categories (right). Ribbons indicate gene-to-term associations for *Mtor*, *Pik3ca*, *Rptor*, *Akt2*, *Akt1*, and *Tert*. (**C**) Scatter plot showing a significant negative correlation between *Akt1* and *Akt2* gene expression (normalized counts). The regression line and 95% confidence interval indicate that loss of *Cryba1* leads to a reciprocal shift in AKT isoform expression. Pearson’s correlation coefficient R^2^ = -0.6, ** *p* =0.04. Each data point represents an individual biological replicate (n = 3). (**D**) Immunoblot analysis of phospho-AKT2, AKT2 and H3 loading control levels in RPE from 18-month-old control and *cryba1* cKO mice. (**E–F**) Enriched signaling pathways and biological processes identified by RNA-seq in *Akt2*-KI (**E**), *akt2*-cKO (**F**). The y-axis represents biological processes; dot size indicates gene count and color represents the -log10 p value. (**G**) Heatmap showing relative gene expression levels (Z-score) of insulin‑signaling components (IRS1, GAB1, IGF1R, SOCS1) across replicates of iPSC‑RPE from AMD subjects treated with DMSO or AKT2 inhibitor (n = 3). (**H**) Glucose uptake was measured by GLO luminescence in *CFH* ^Y402H^ RPE cells. AKT2 inhibitor (AKT2i) significantly increased glucose uptake relative to DMSO, whereas wortmannin completely abolished uptake. Rapamycin partially rescued glucose uptake. Data are represented as mean ± SD. n = 3 biological replicates per group. Statistical significance was determined by one-way ANOVA with Tukey’s post-hoc test; *****p* < 0.0001, ns= not significant.

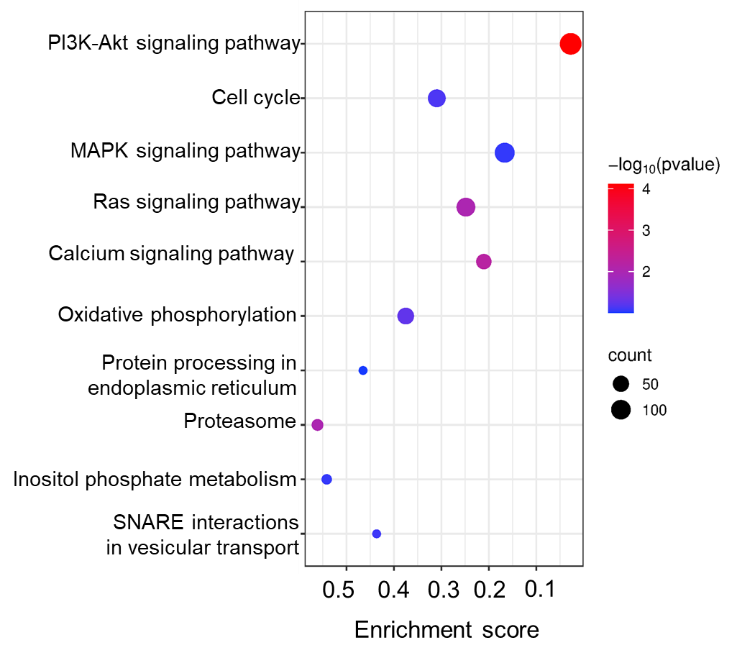

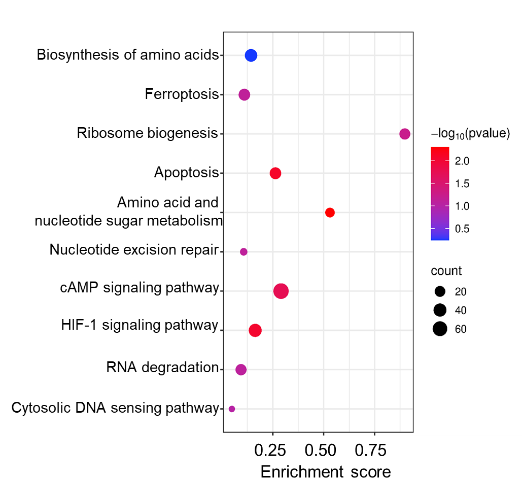

A

C

D

E

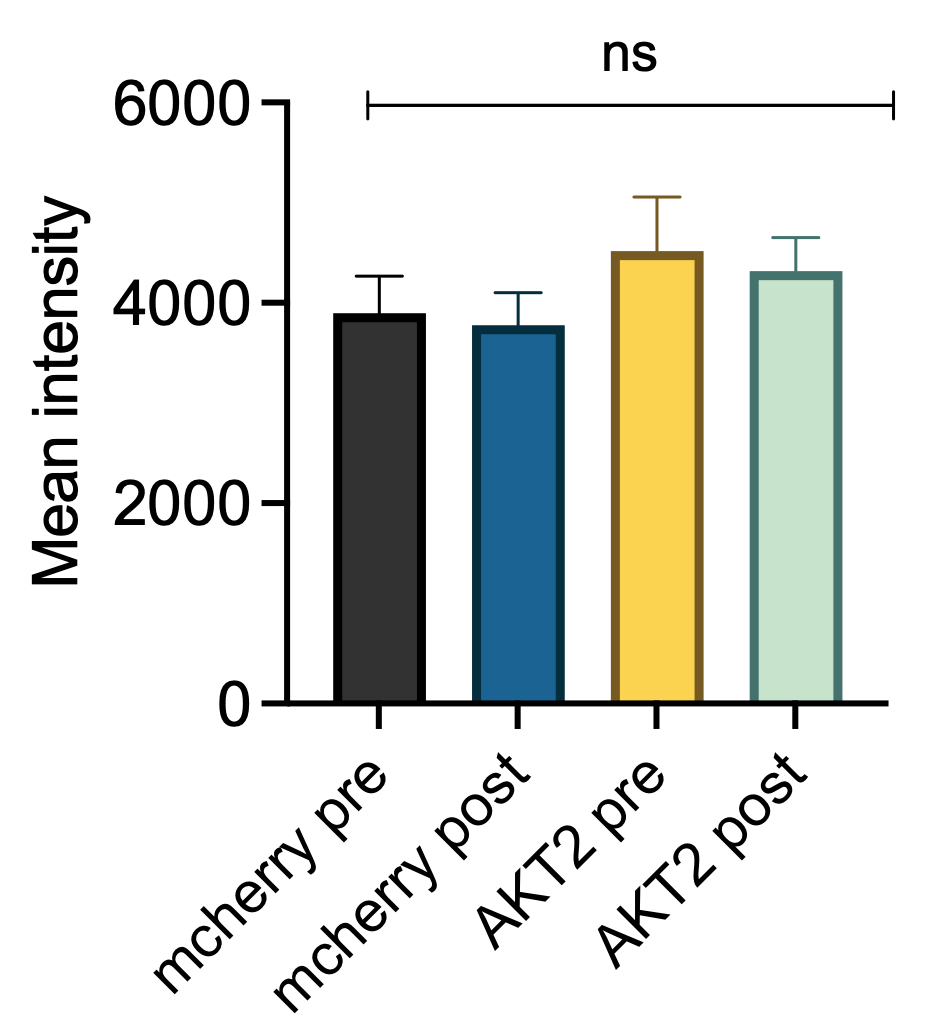

F

G

H

I

K

L

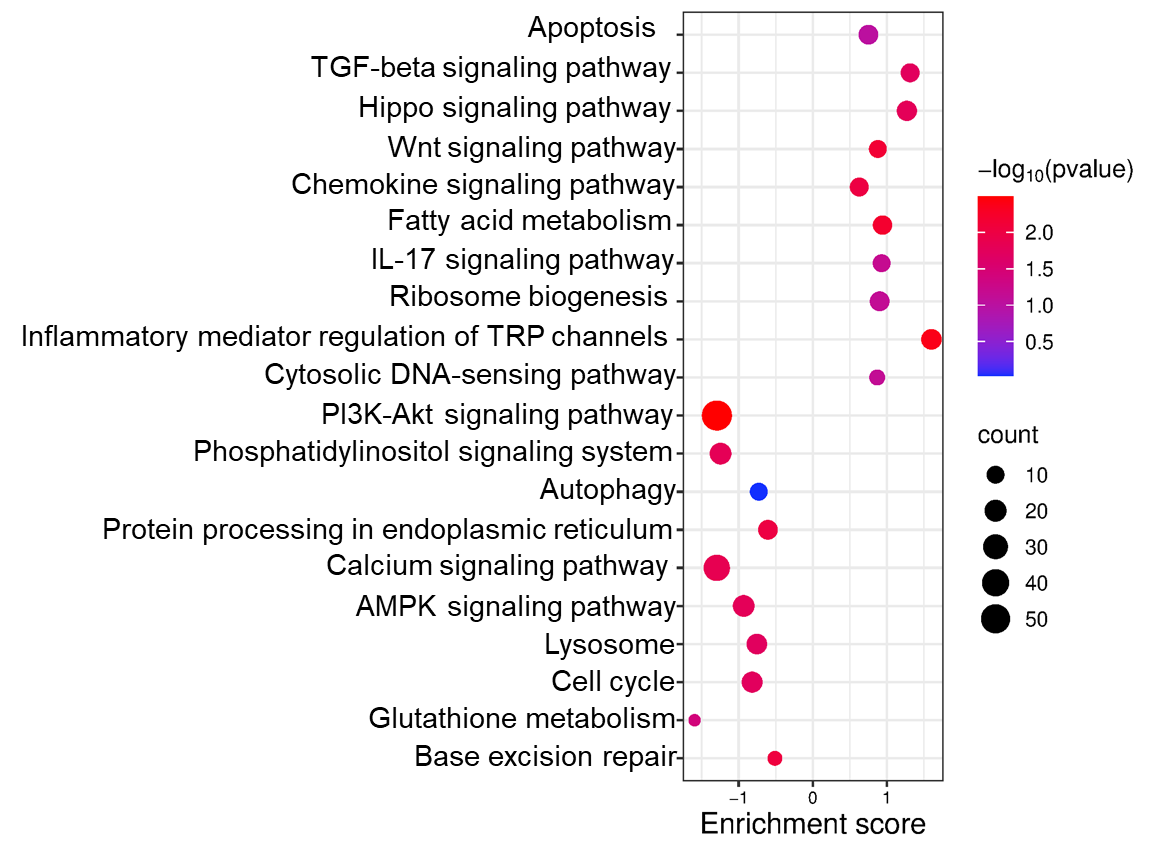

Control RPE vs AMD RPE

Upregulated

Downregulated

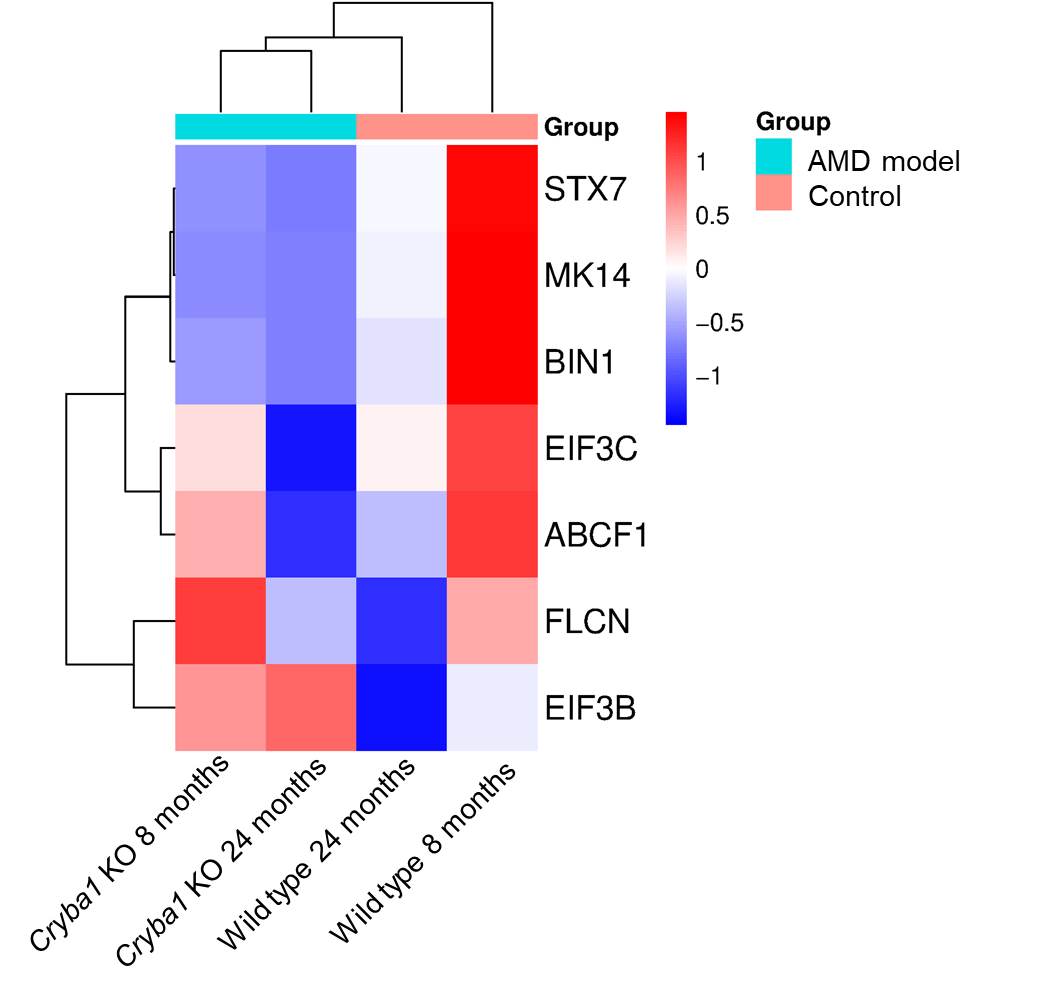

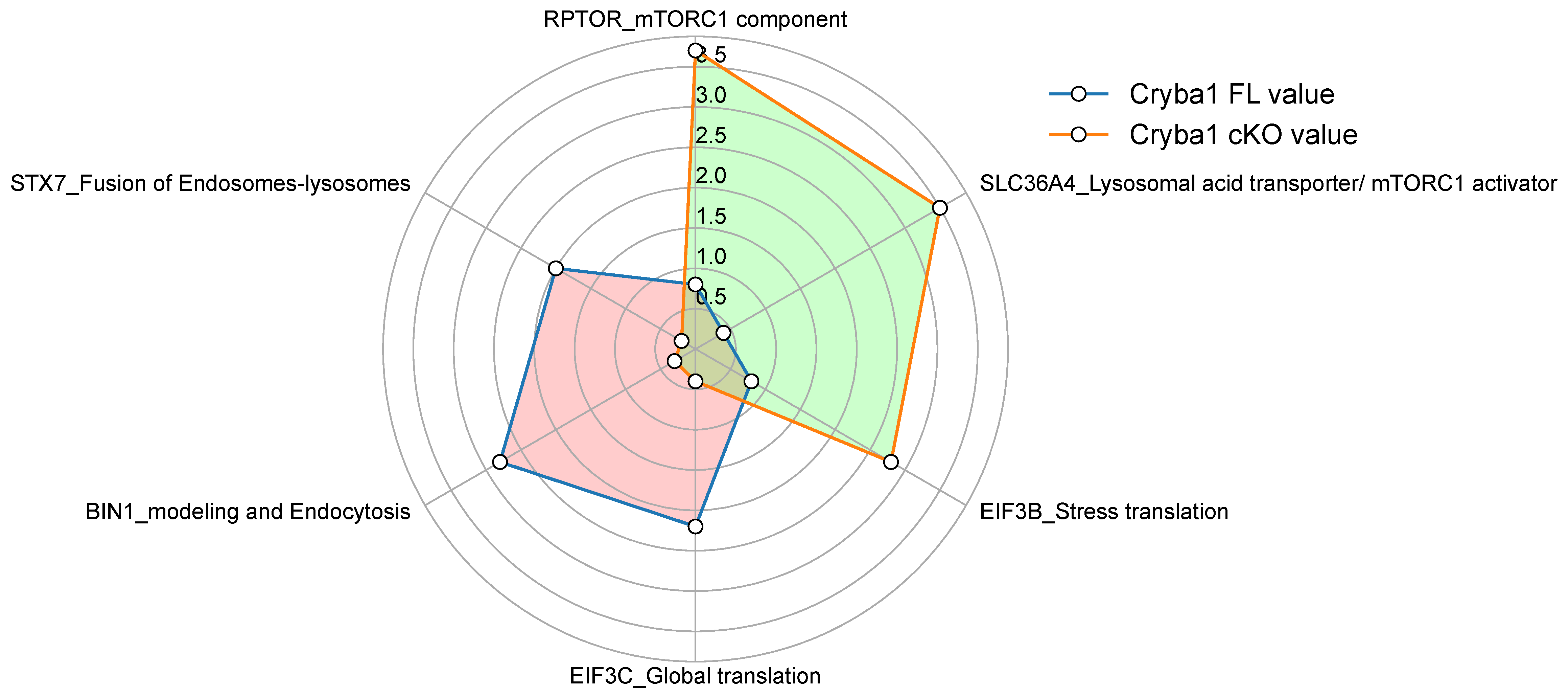

*Cryba1 FL*

*cryba1 cKO*

M

N

B

J

O

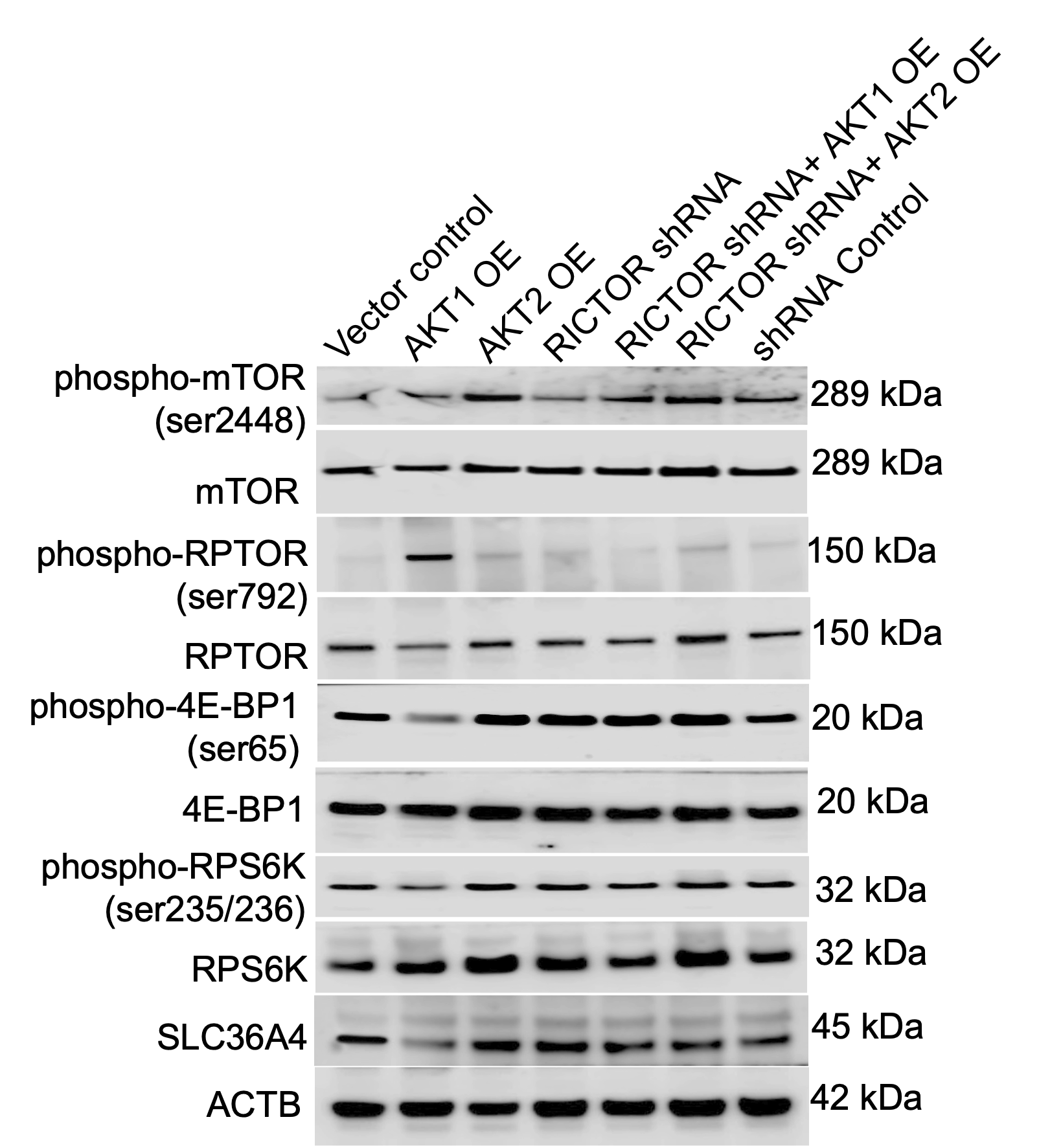

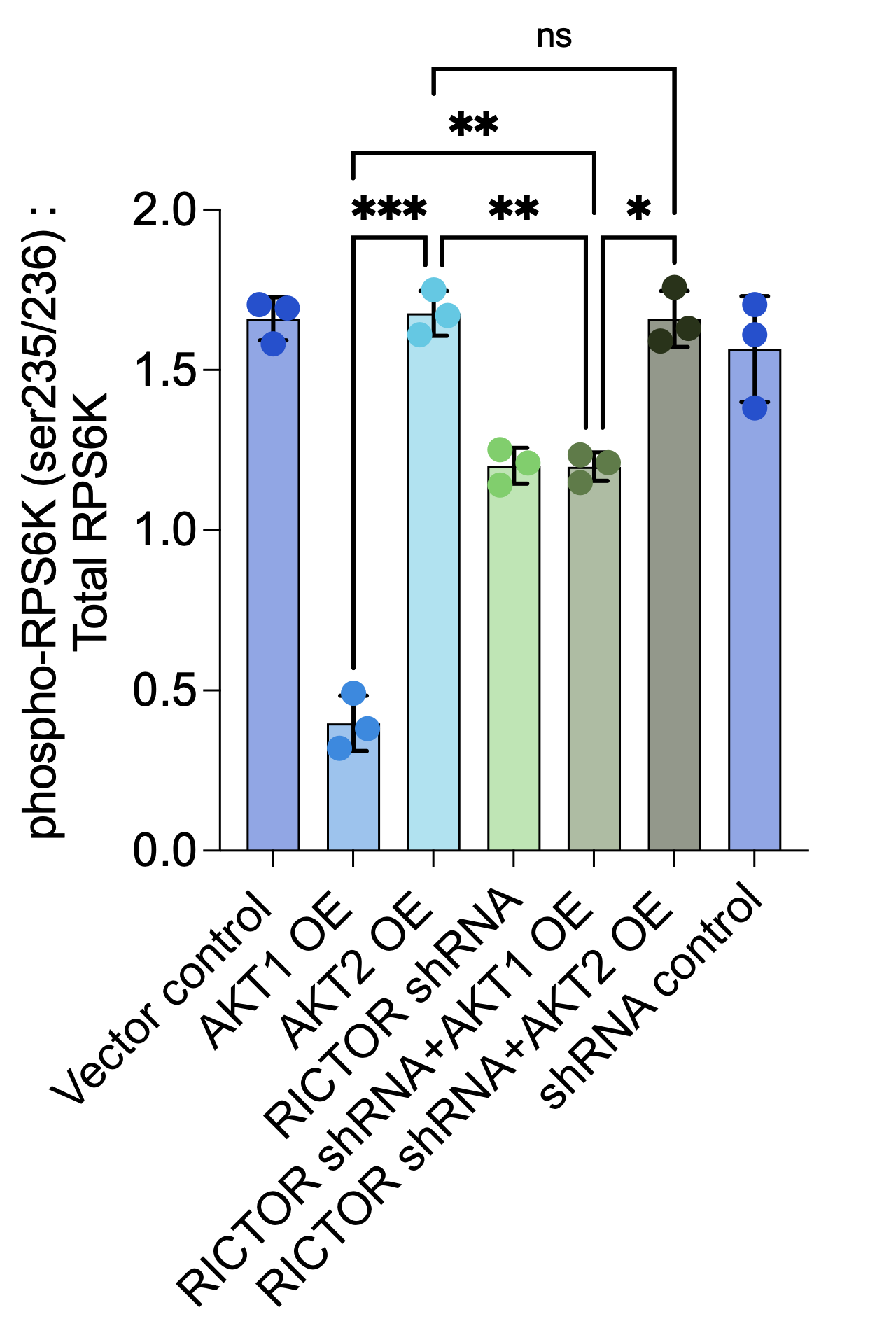

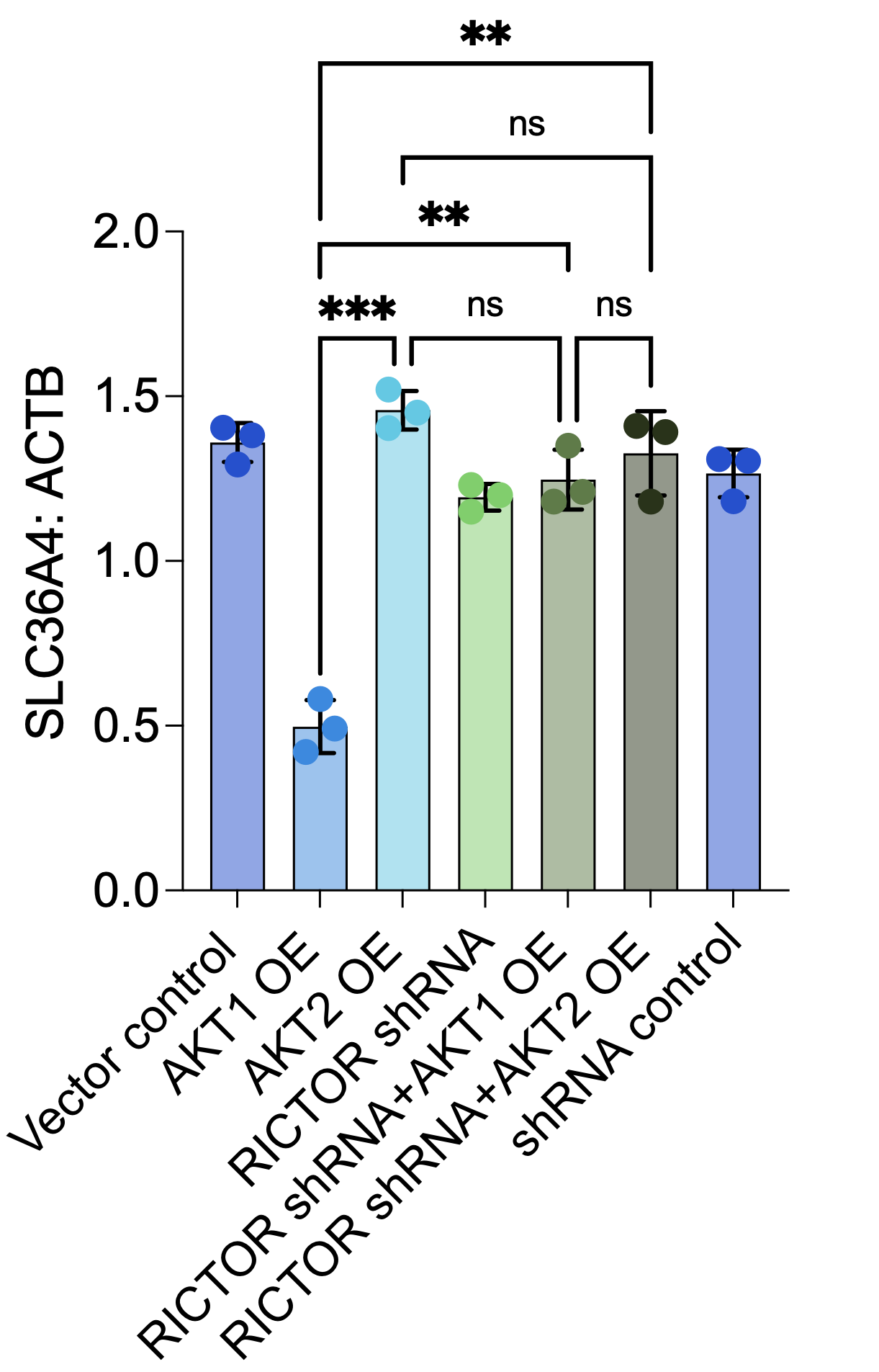

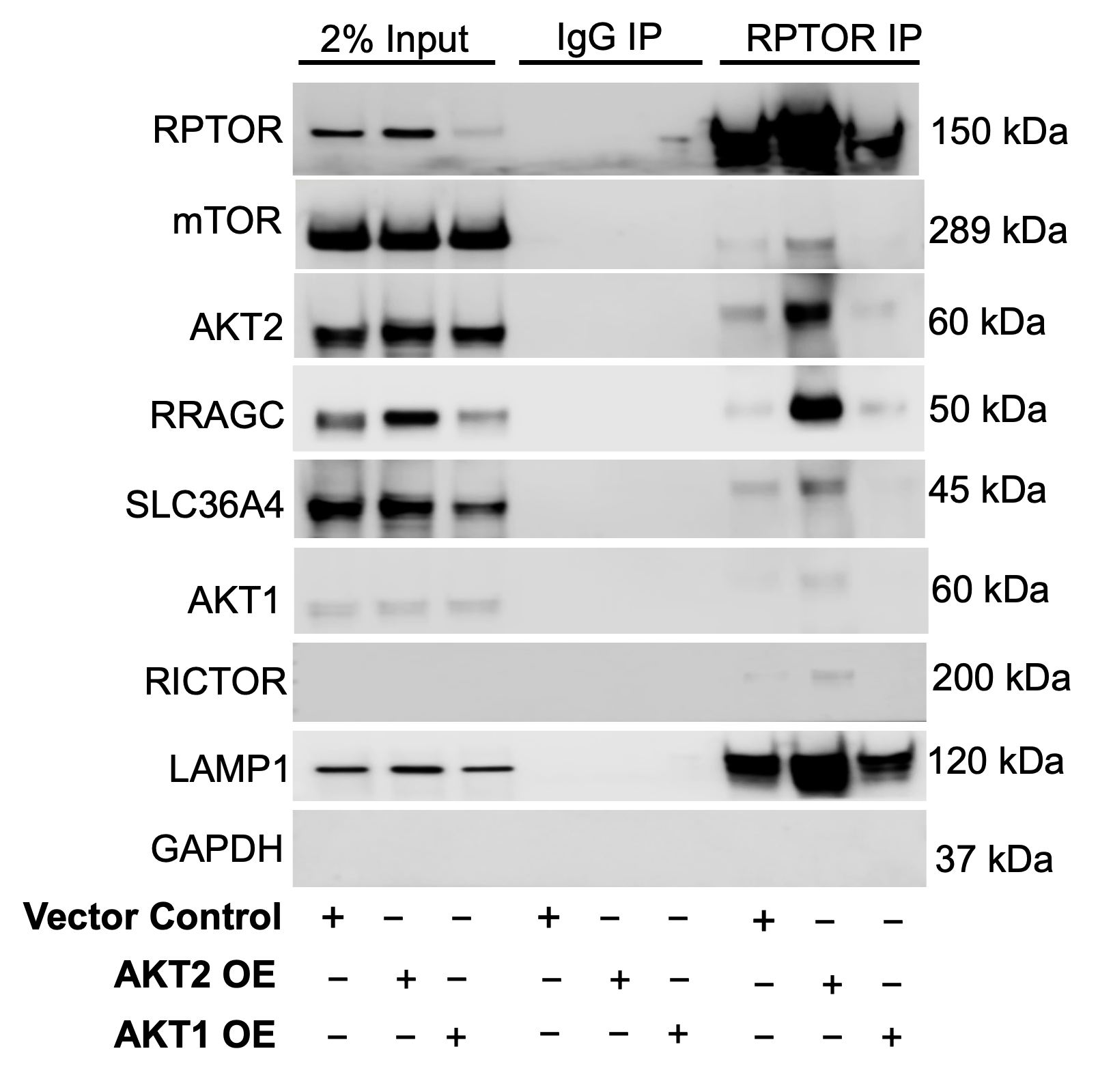

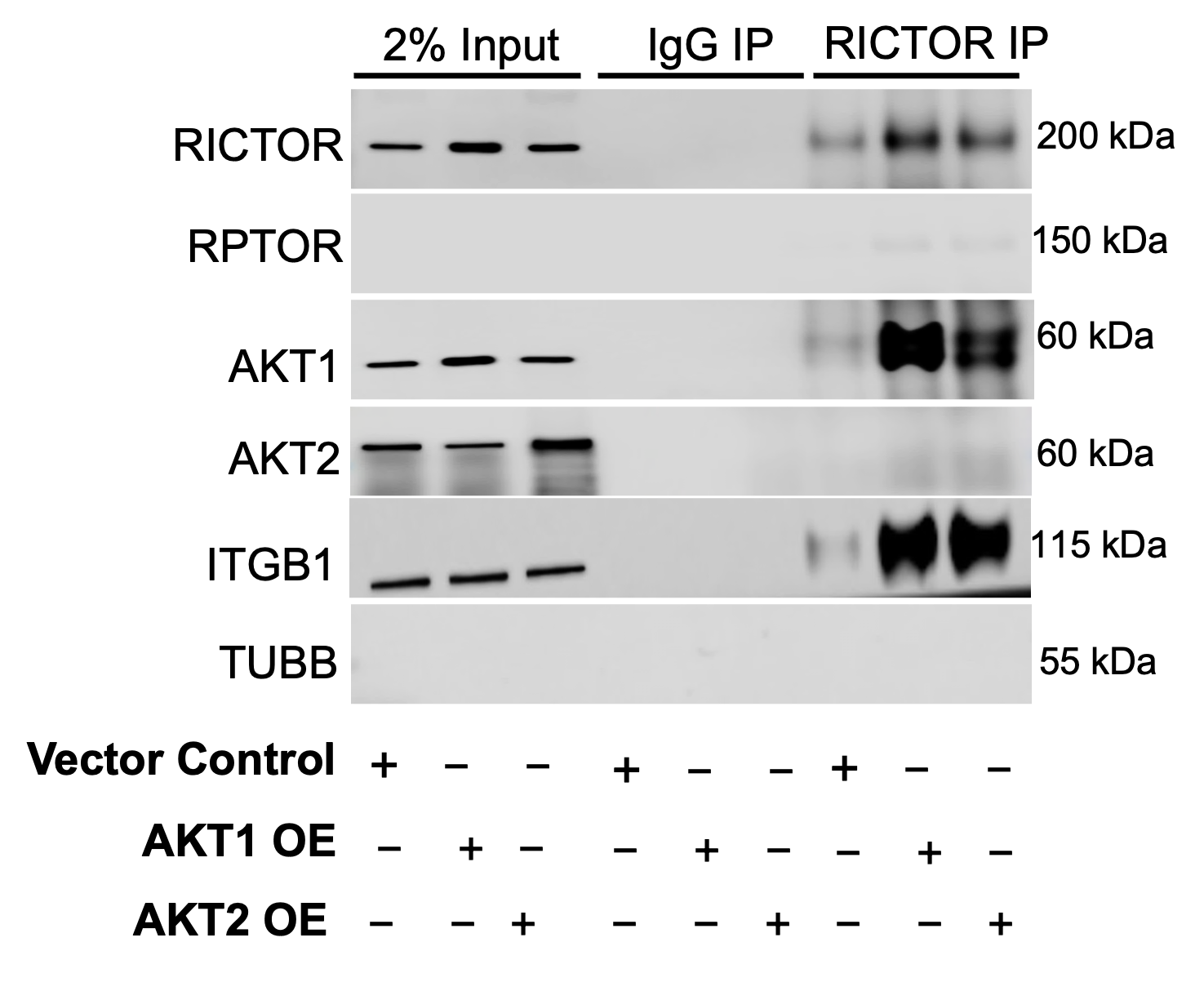

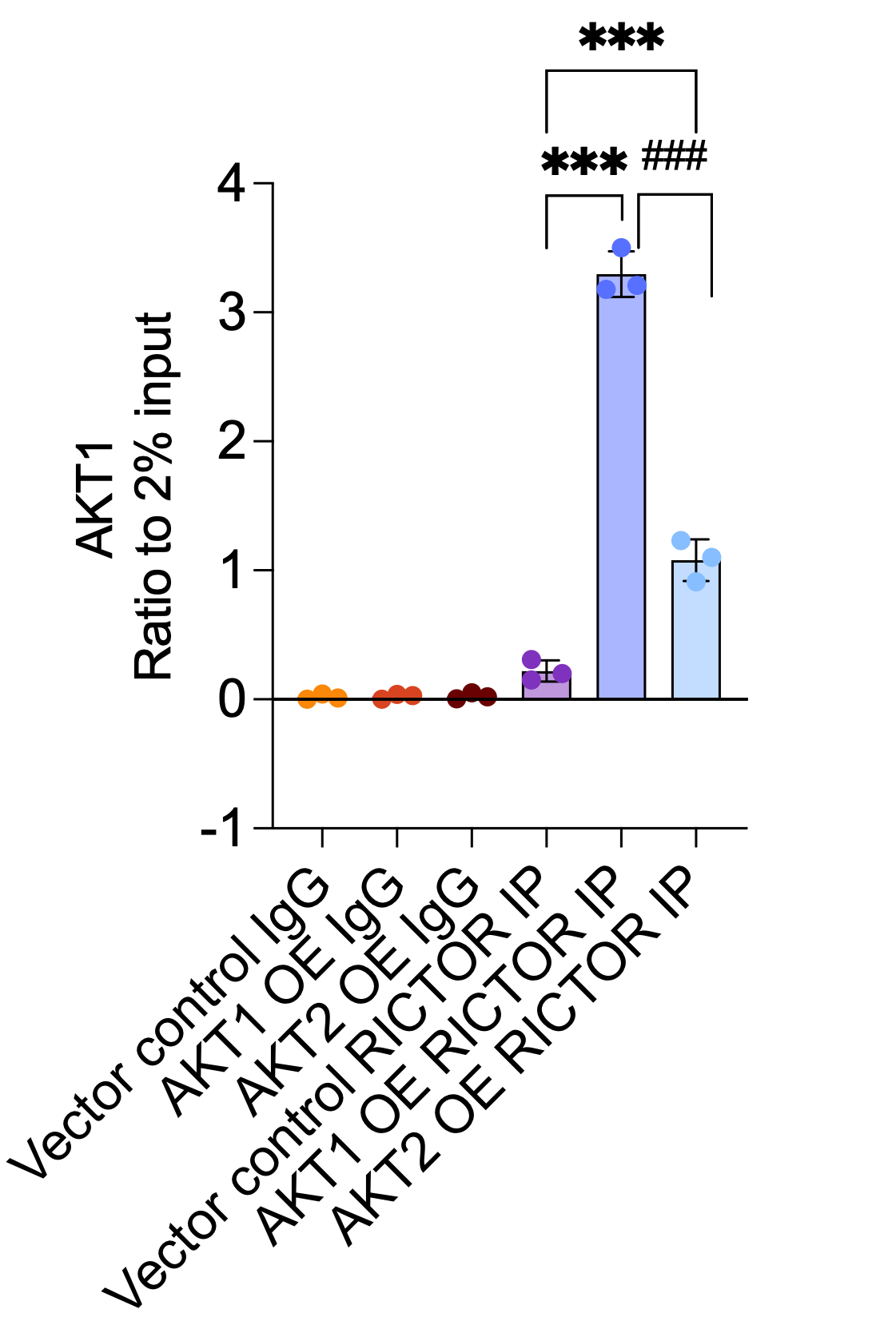

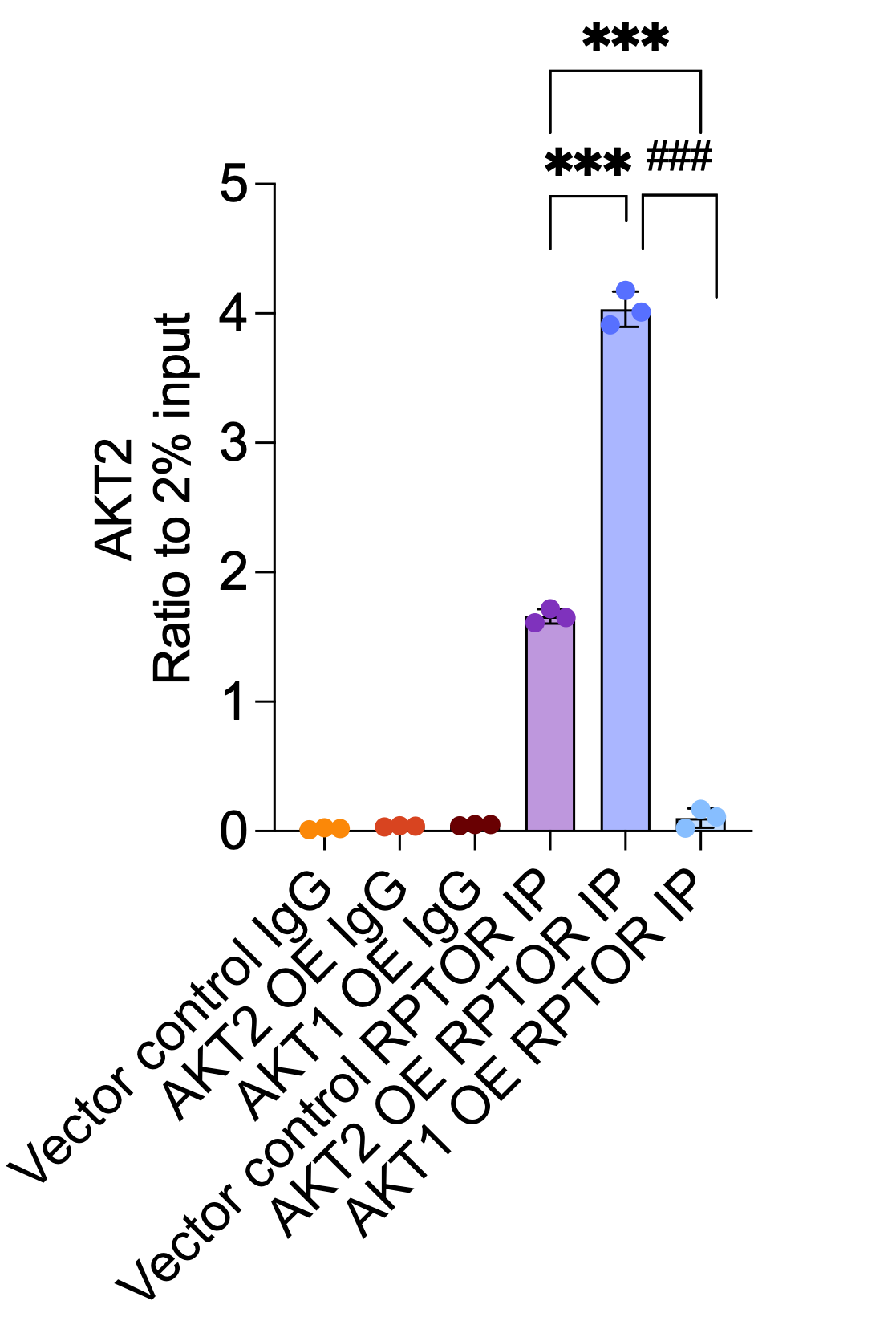

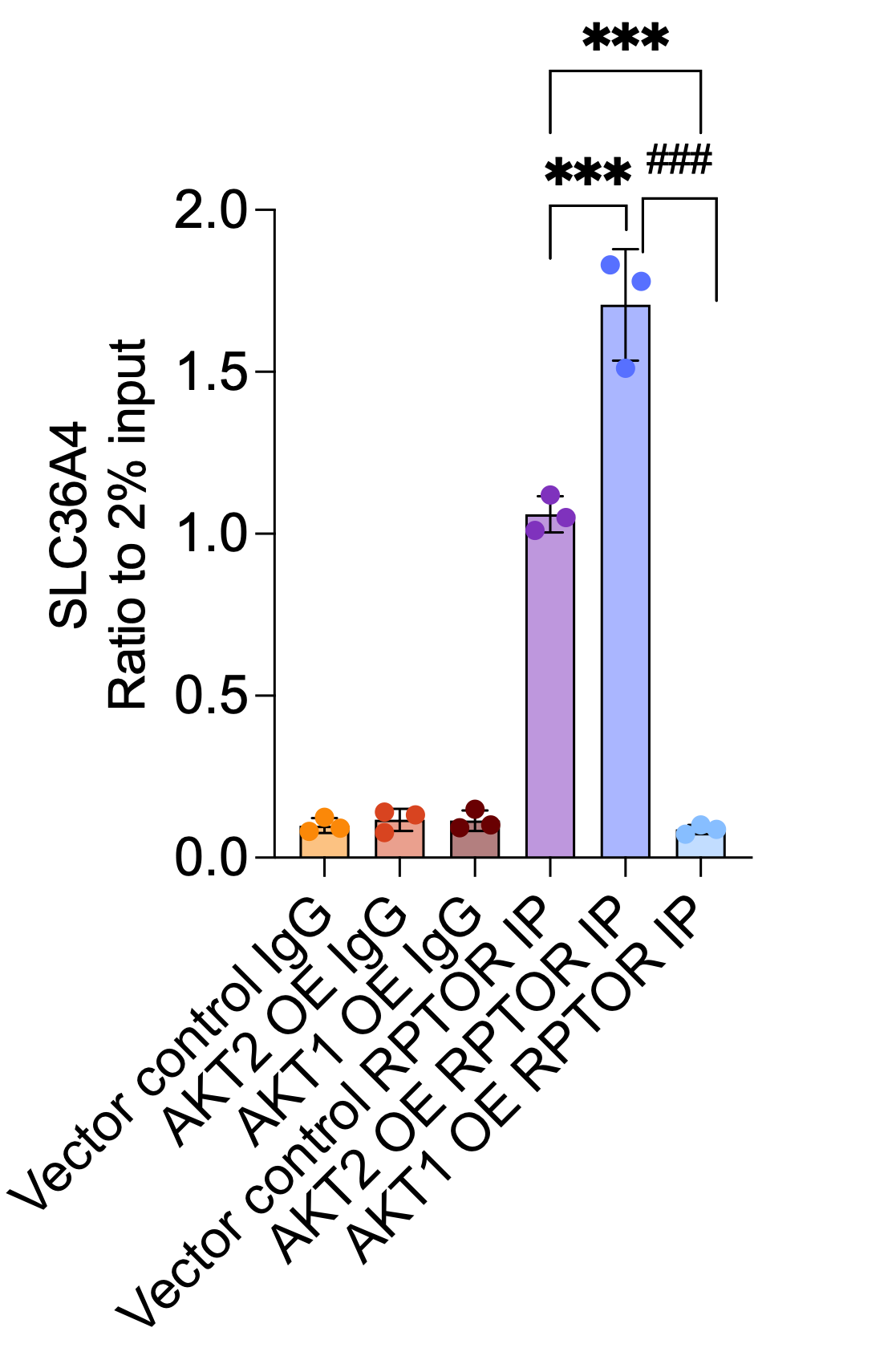

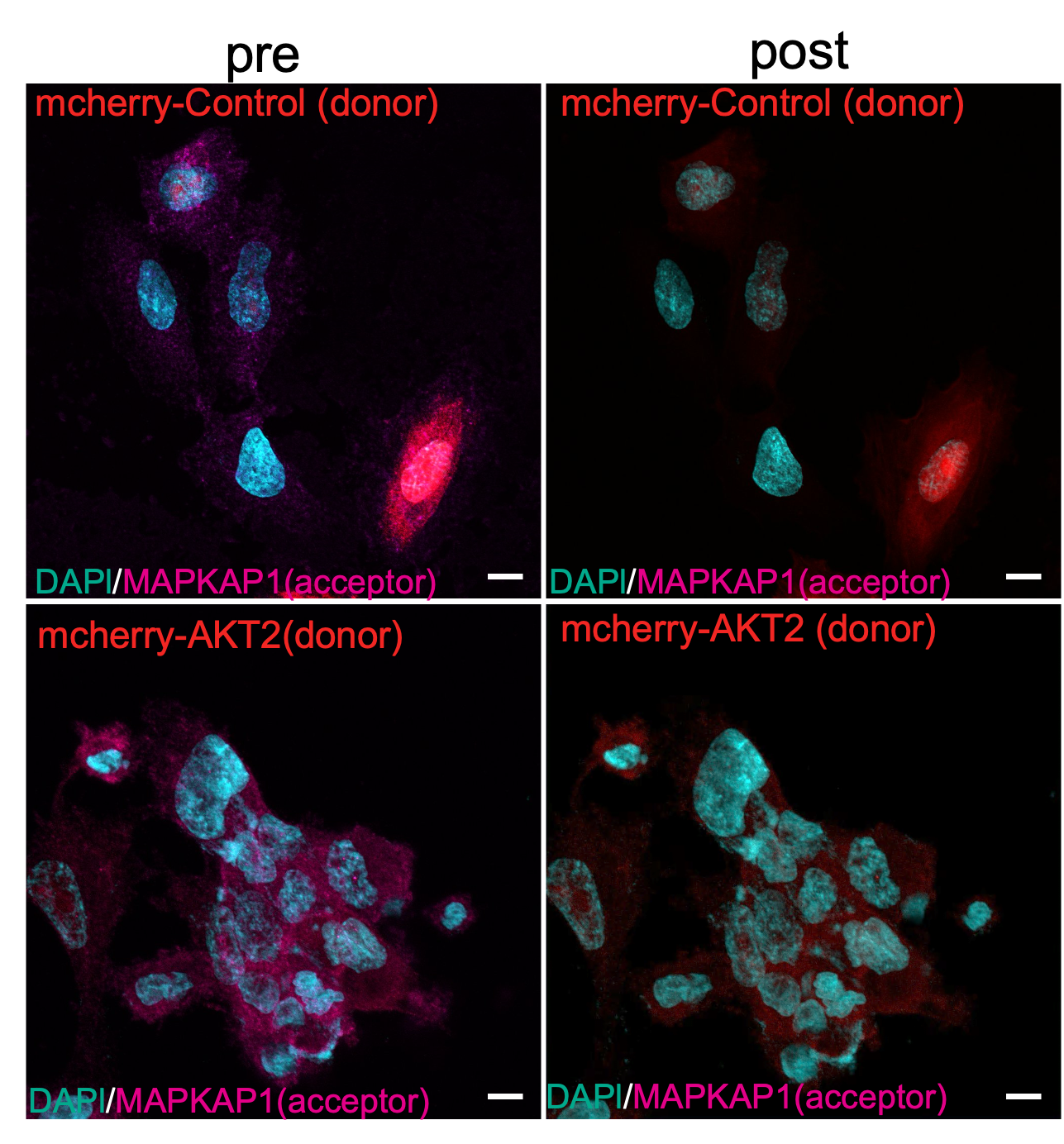

**Figure S2. Changes in mTORC1 signaling and endo-lysosomal translation programs in RPE from AMD models.** (**A**) Proteomics showing normalized expression (row Z-score) of selected proteins involved in mTORC1 signaling and translation (STX7, MKI4, BIN1, EIF3C, AP1EC1, FLCN, EIF3B) across *Cryba1* flox (FL) and *cryba1* conditional knockout (cKO) RPE (n = 2 per group). Hierarchical clustering (Euclidean distance, complete linkage) groups samples by control and AMD models. Scale bar indicates Z‑score. (**B**) Radar plot of integrated RNA seq and proteomics analysis identified mTORC1 specific pathway in *Cryba1* FL and *Cryba1* cKO RPE. Radial axes represent normalized enrichment scores for selected GO/Reactome terms (labels shown); shaded areas indicate the distribution for each condition. Pathways shown include mTORC1/ribosomal translation, lysosomal transport, endosomal/lysosomal fusion and stress‑translation responses. (**C–E**) Dot plots of pathway enrichment for (**C**) Control RPE vs AMD RPE (**D**) upregulated pathways in DMSO vs AKT2i treatment in iPSC RPE *CFH* ^Y402H^ and (**E**) Downregulated pathways in DMSO vs AKT2i treatment in iPSC RPE *CFH* ^Y402H^. Dot position and color indicate enrichment statistic (x‑axis, enrichment score; color, –log10 p‑value), and dot size indicates the number of genes in each term. Significance thresholds and enrichment method are described in methods. (**F**) Continuation of immunoblot analysis of mTOR complex components in human iPSC‑RPE *CFH* ^Y402H^ cells following overexpression (OE) of AKT1 or AKT2 and shRNA‑mediated knockdown of RICTOR (from Fig. 2J). The mTORC1 component RPTOR and its immediate downstream targets 4EBP1 and RPS6K were increased upon AKT2 OE, whereas RICTOR knockdown did not affect their expression. AKT1 OE significantly reduced SLC36A4 expression, and this reduction was reversed by RICTOR knockdown. n = 3 biological replicates. (**G, H**) Quantification of phospho- RPS6K (G) and SLC36A4 (H) in western blot signals from (**F**). (**I**) Representative RICTOR co-immunoprecipitation (Co-IP) and immunoblots showing high AKT1/RICTOR and low AKT2/RICTOR interactions in plasma membrane fractions of iPSC RPE *CFH* ^Y402H^ cells. ITGB1 as plasma membrane fraction control, TUBB for cytoplasm fraction control. Blots are representative of n = 3 independent biological replicates. (**J**) Bar graphs showing increased AKT1 interaction with RICTOR. (**K**) Representative RPTOR co-immunoprecipitation (Co-IP) and immunoblots showing high AKT2/RPTOR and low AKT1/RPTOR interactions in lysosomal fractions of iPSC RPE *CFH* ^Y402H^ cells. LAMP1 as lysosomal fraction control, GAPDH for cytoplasm fraction control. Blots are representative of n = 3 independent biological replicates (L-M) Densitometry showing (**L**) AKT2/RPTOR and (**M**) SLC36A4/RPTOR interaction. (**N**) Representative FRET imaging of iPSC RPE *CFH* ^Y402H^ cells expressing mcherry-Control or mcherry-AKT2 (donor) and stained for magenta-MAPKAP1 (acceptor). Quantification of FRET efficiency via acceptor photobleaching of magenta-MAPKAP1 (n=10 cells). (**O**) Bar graph represents mean intensity pre and post acceptor (magenta-MAPKAP1) bleaching demonstrates no direct physical interaction between AKT2 and the mTORC2 subunit MAPKAP1. Scale bars: 10μm Bars represent mean ± SD (n = 3); *p< 0.05, **p < 0.01, ***p < 0.001, ^###^p < 0.001. Statistical significance was determined by one-way ANOVA with post hoc test.

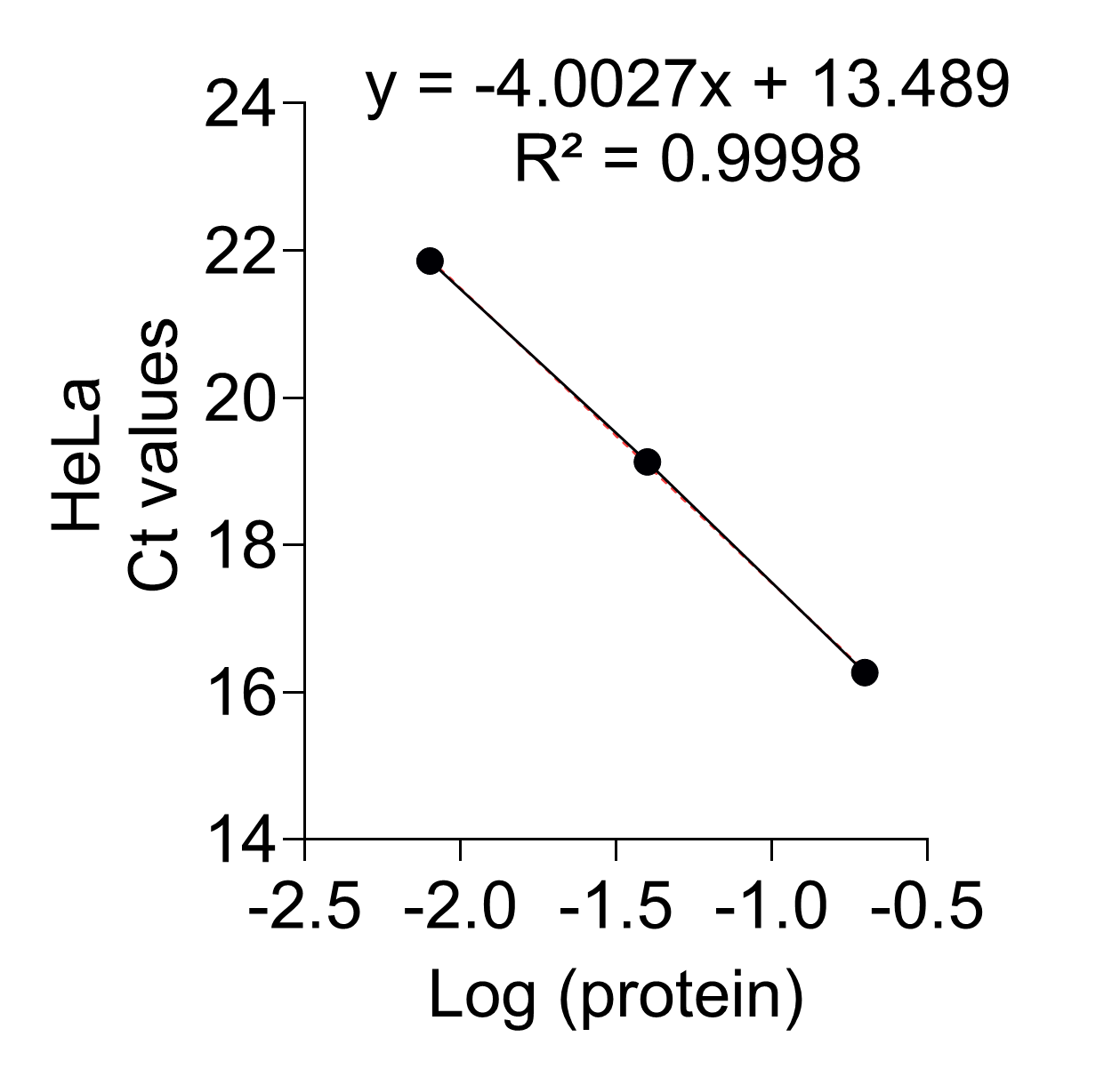

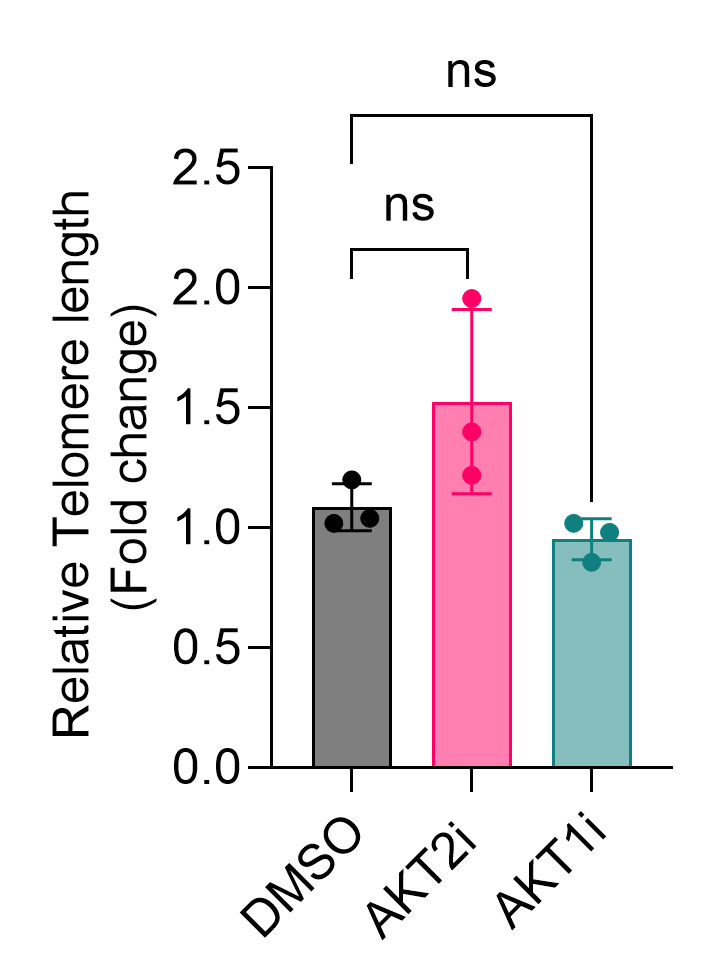

A

B

C

D

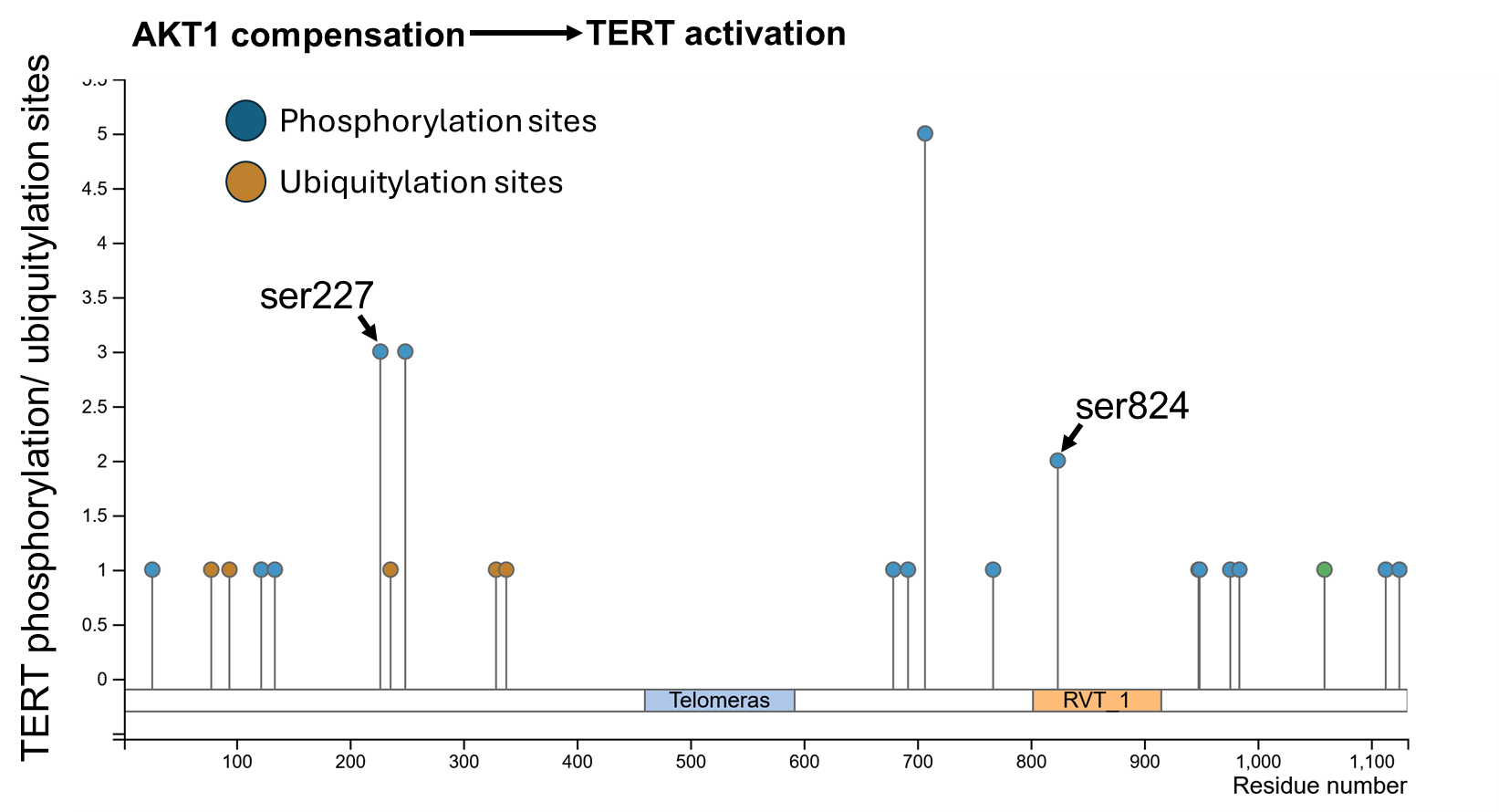

E

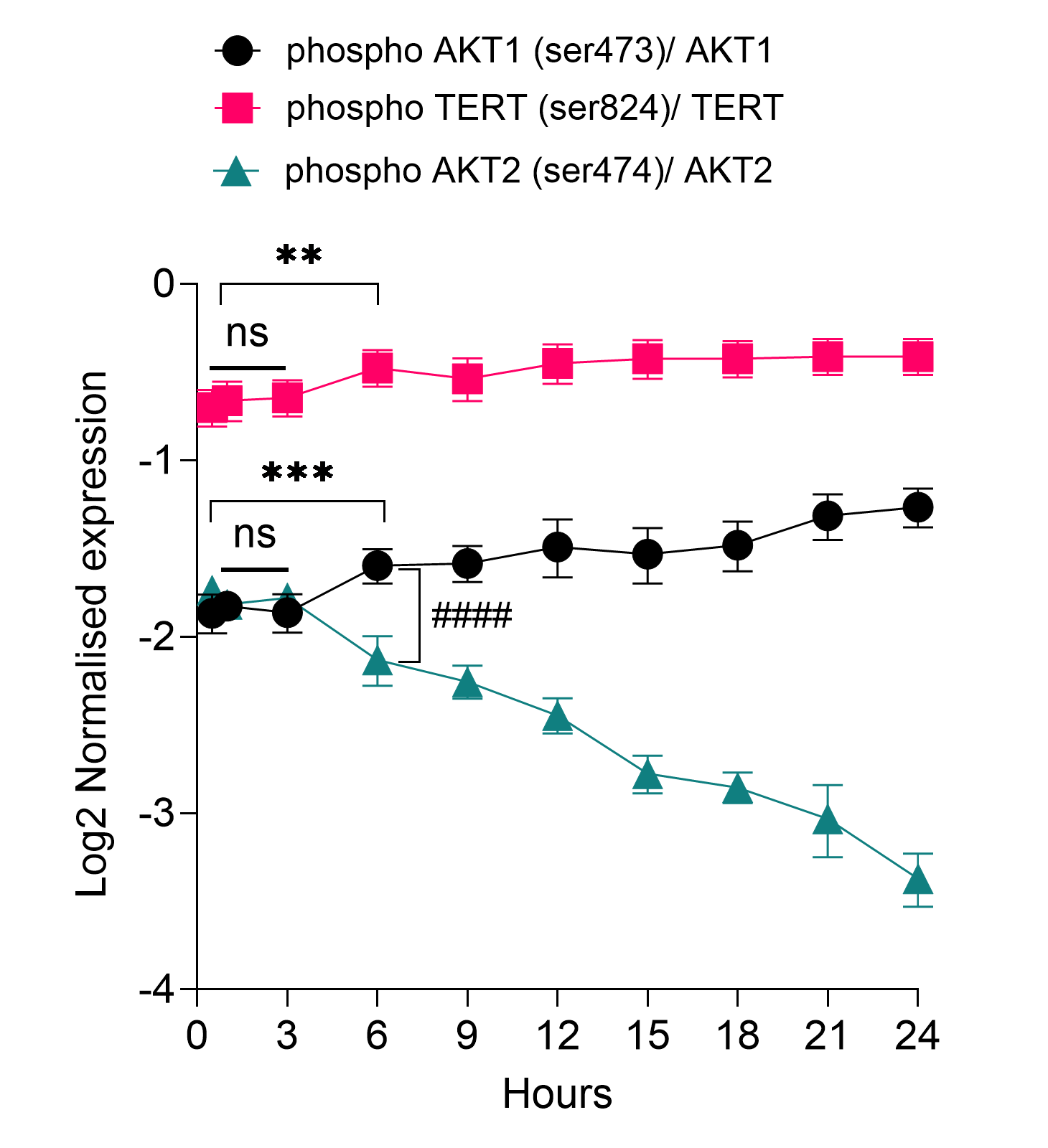

I

J

K

F

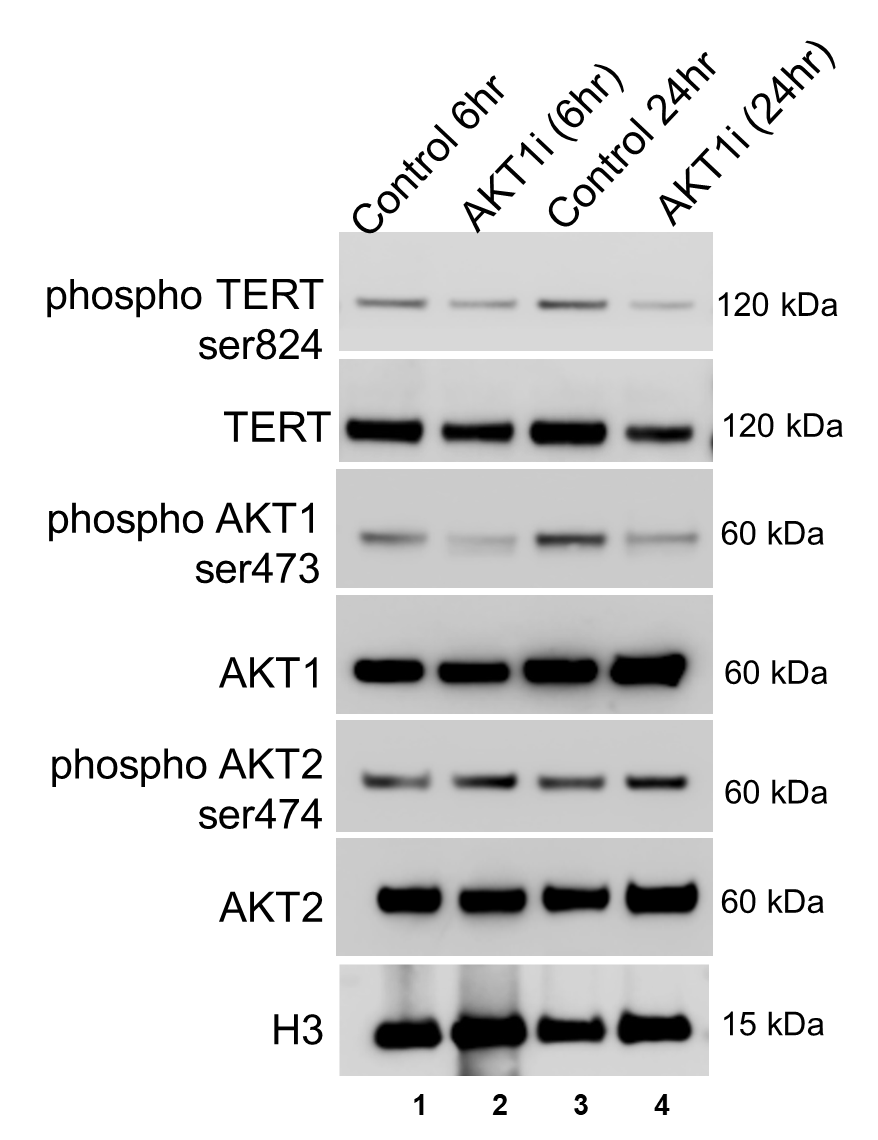

G

H

L

M

N

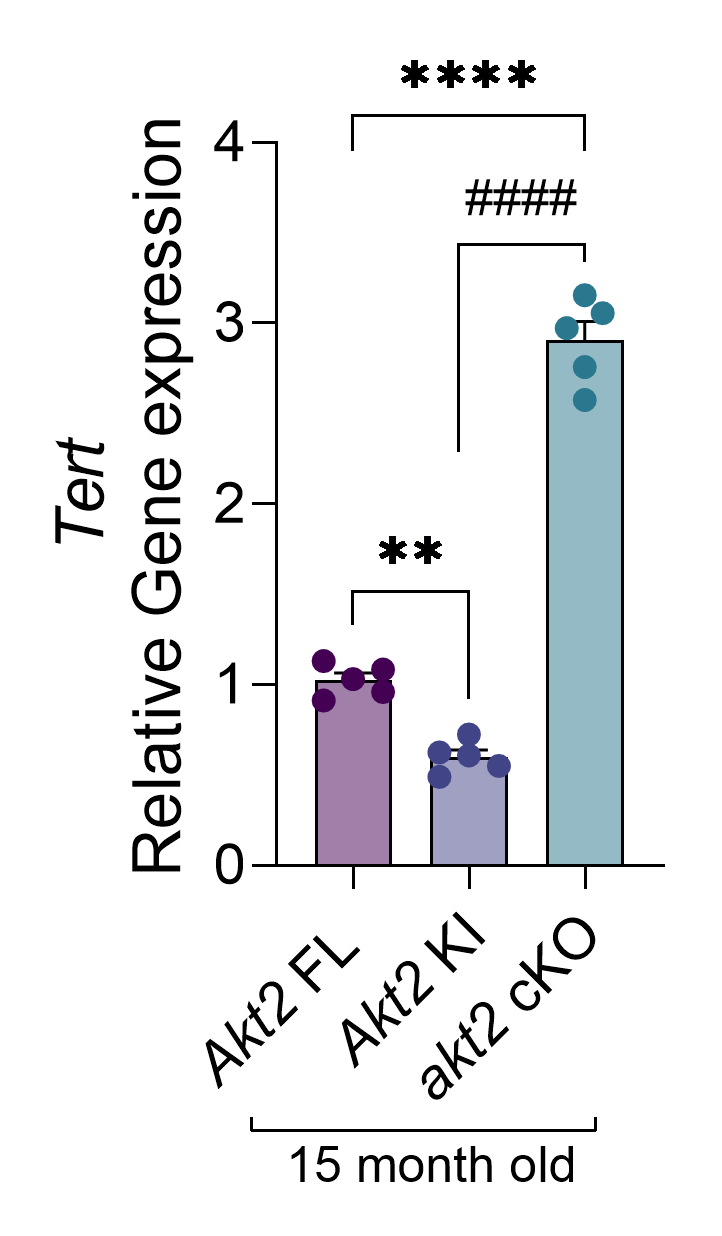

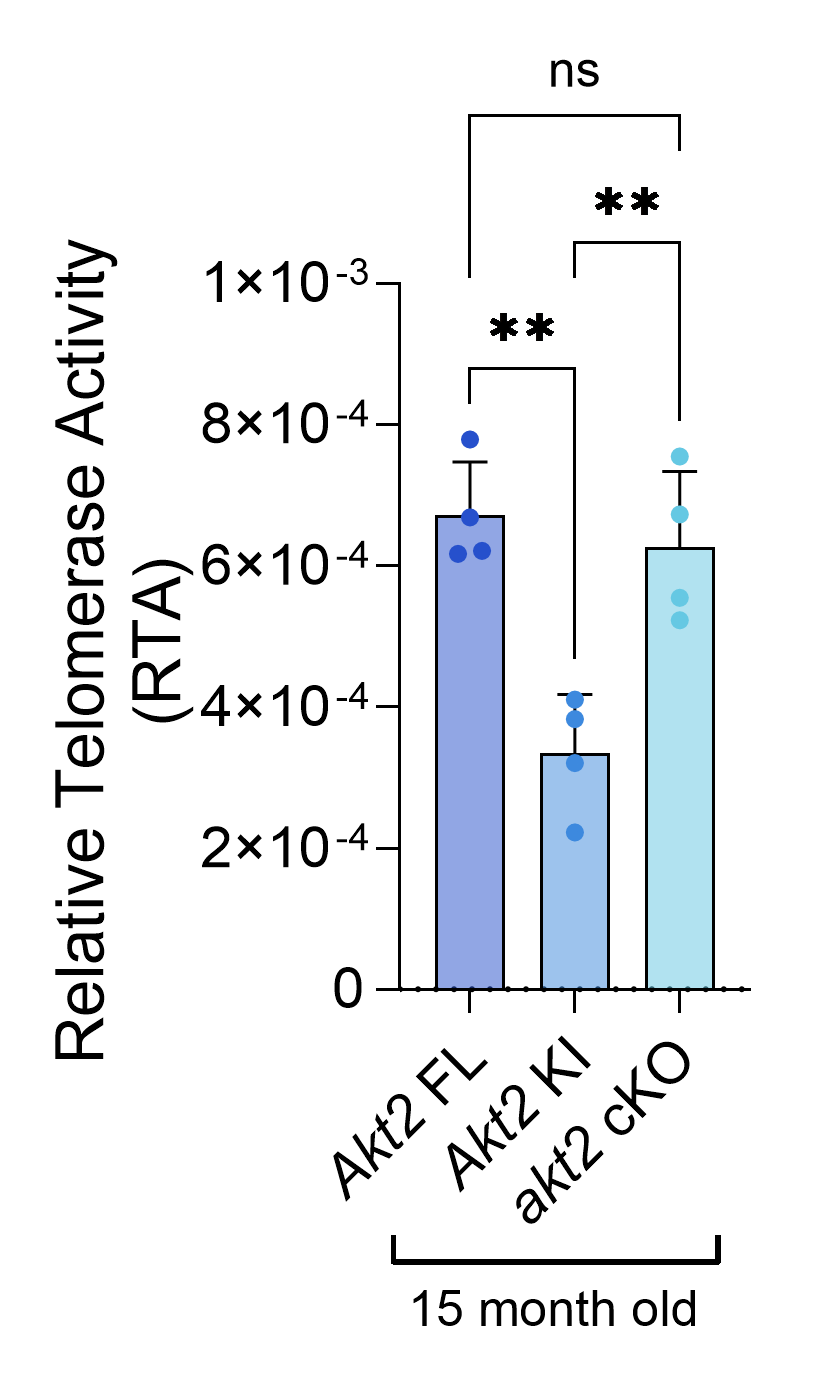

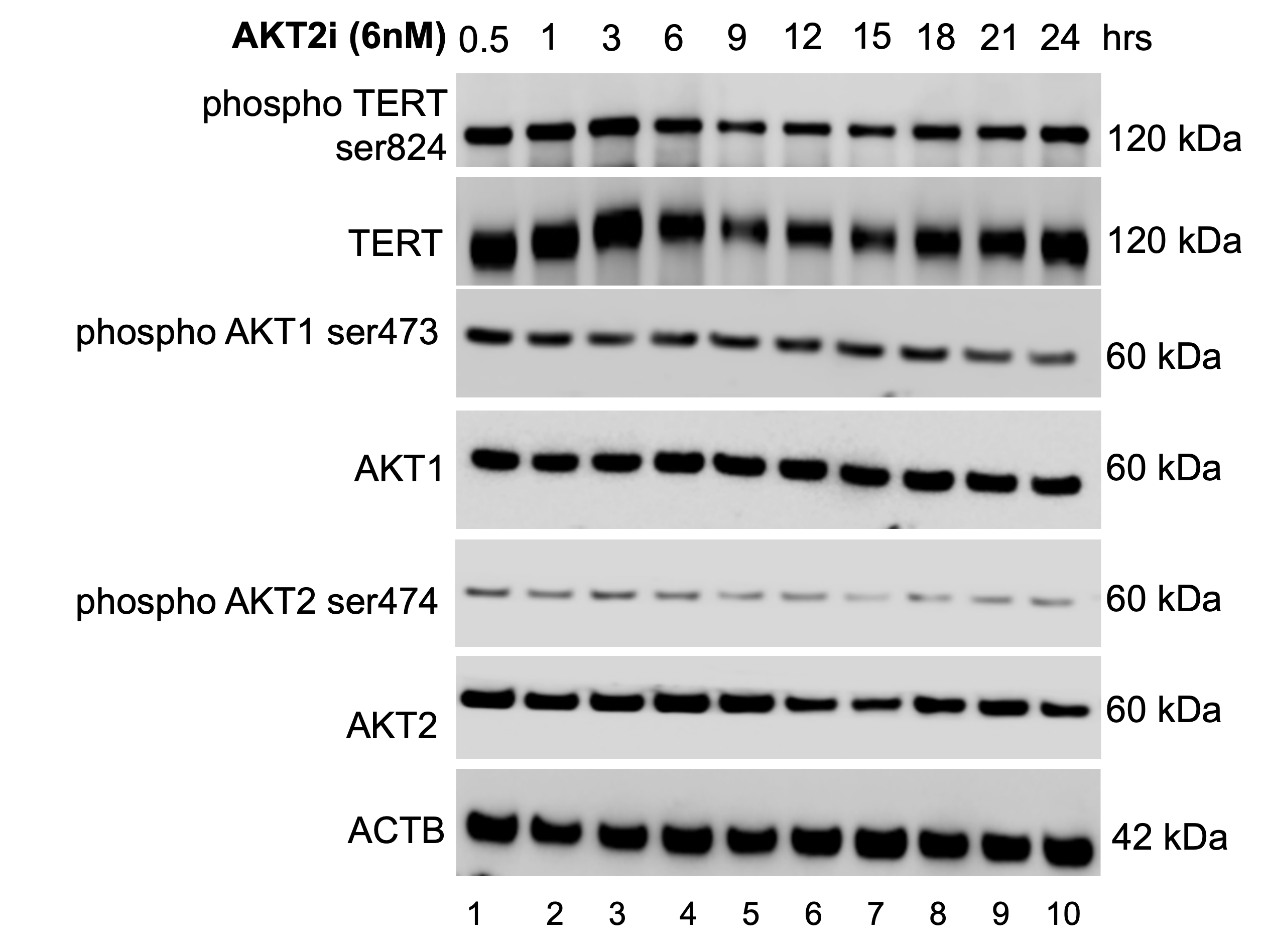

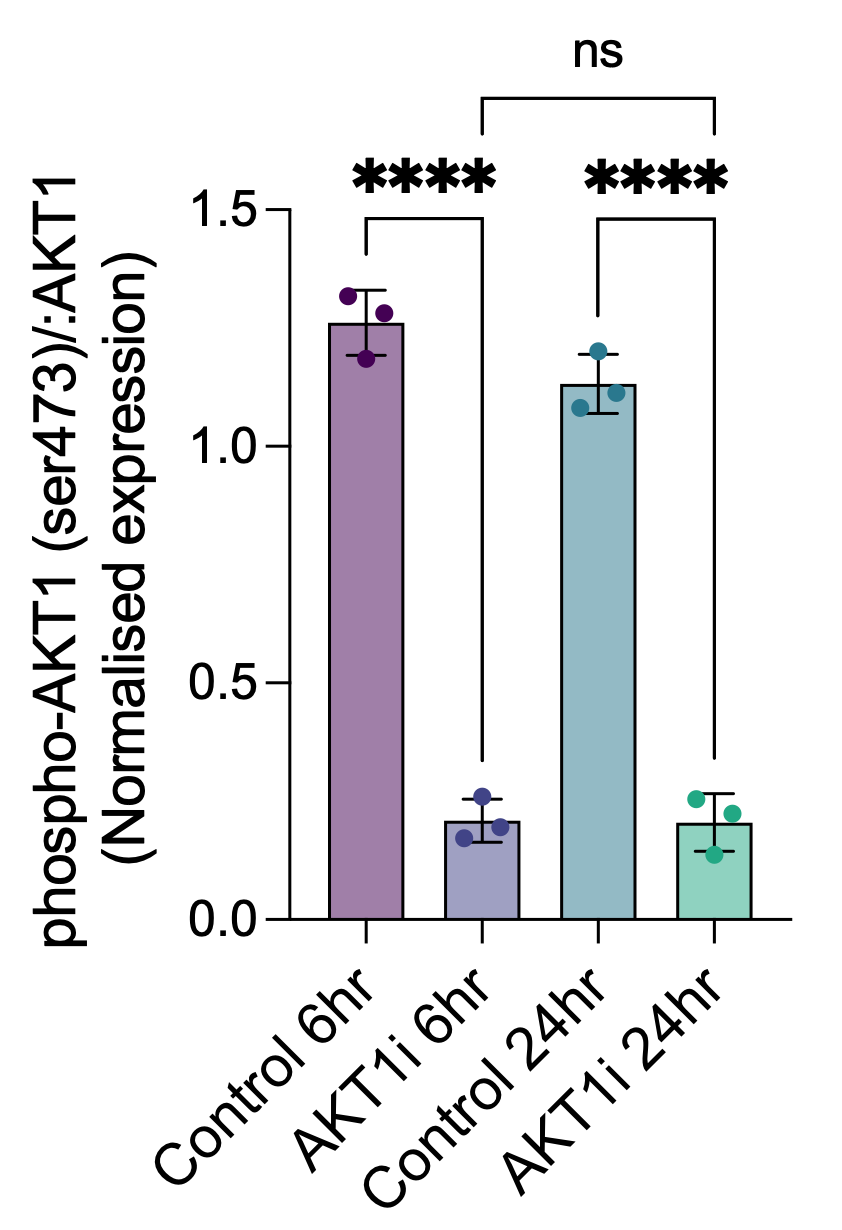

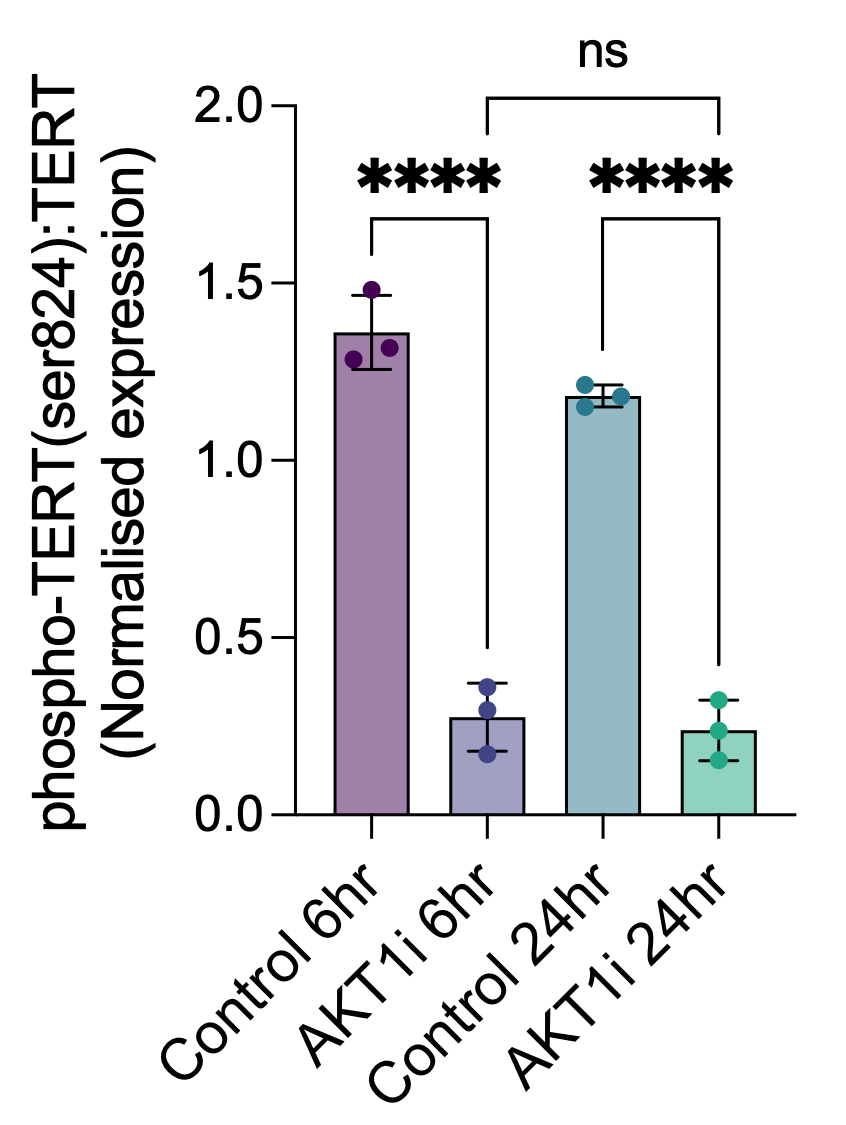

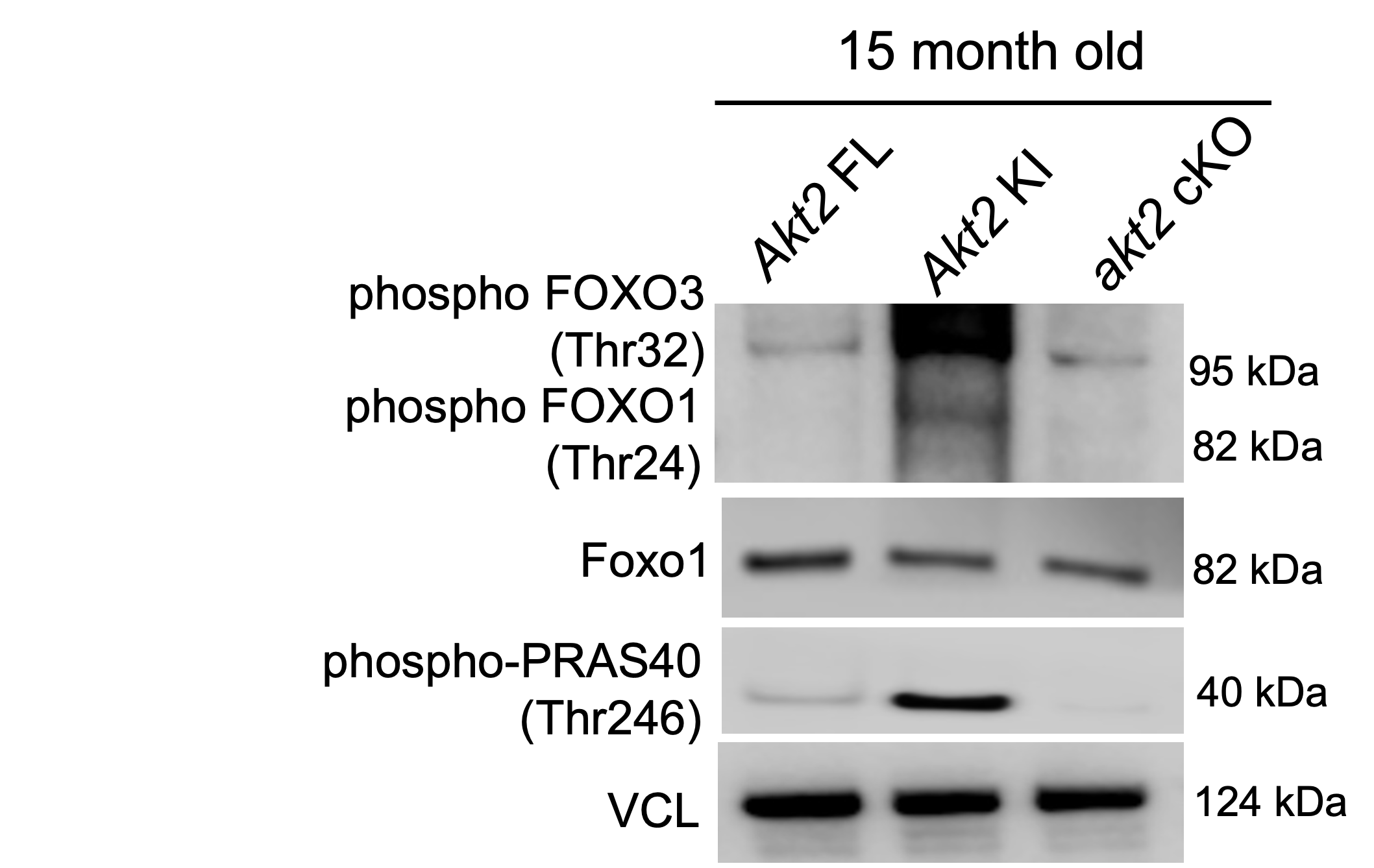

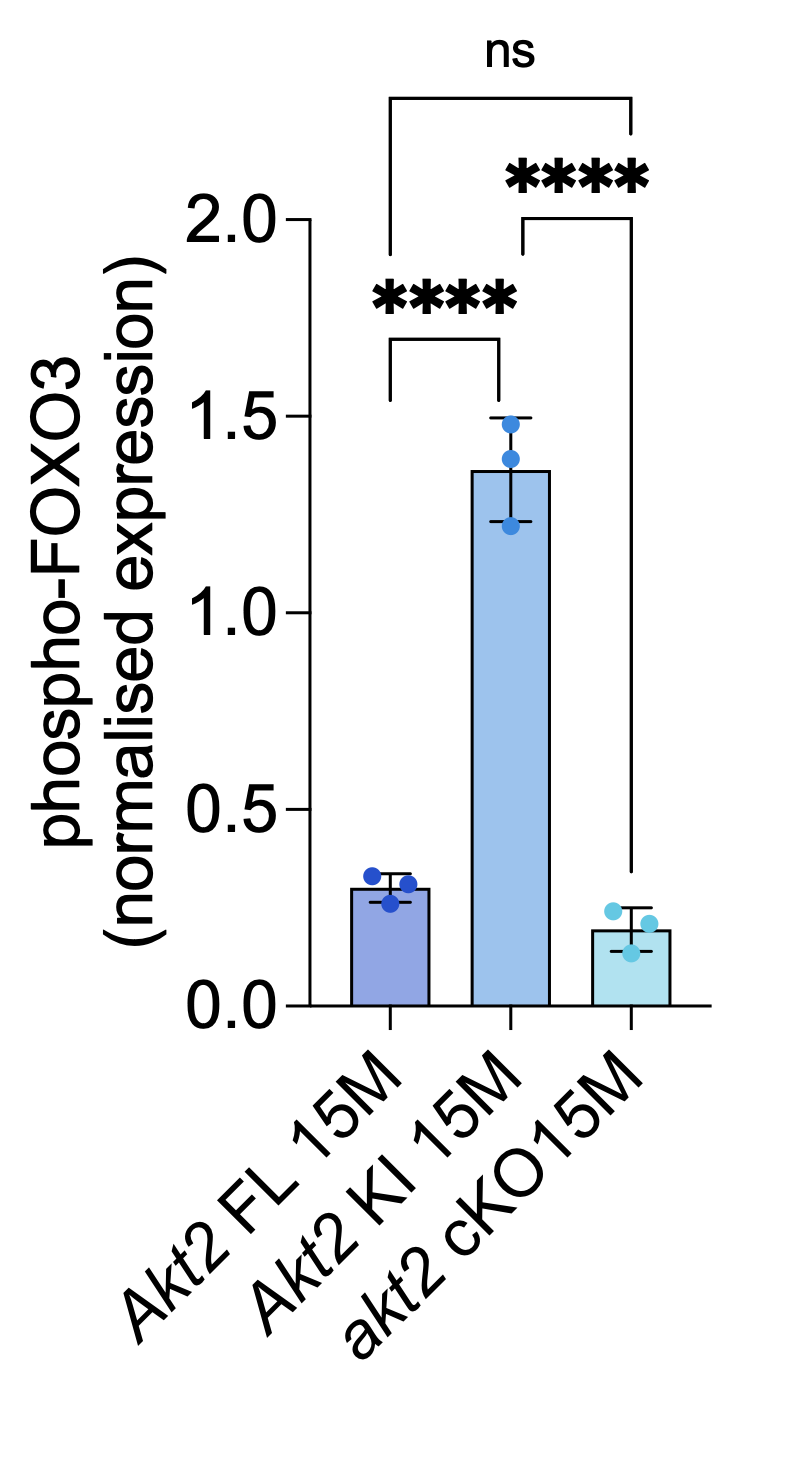

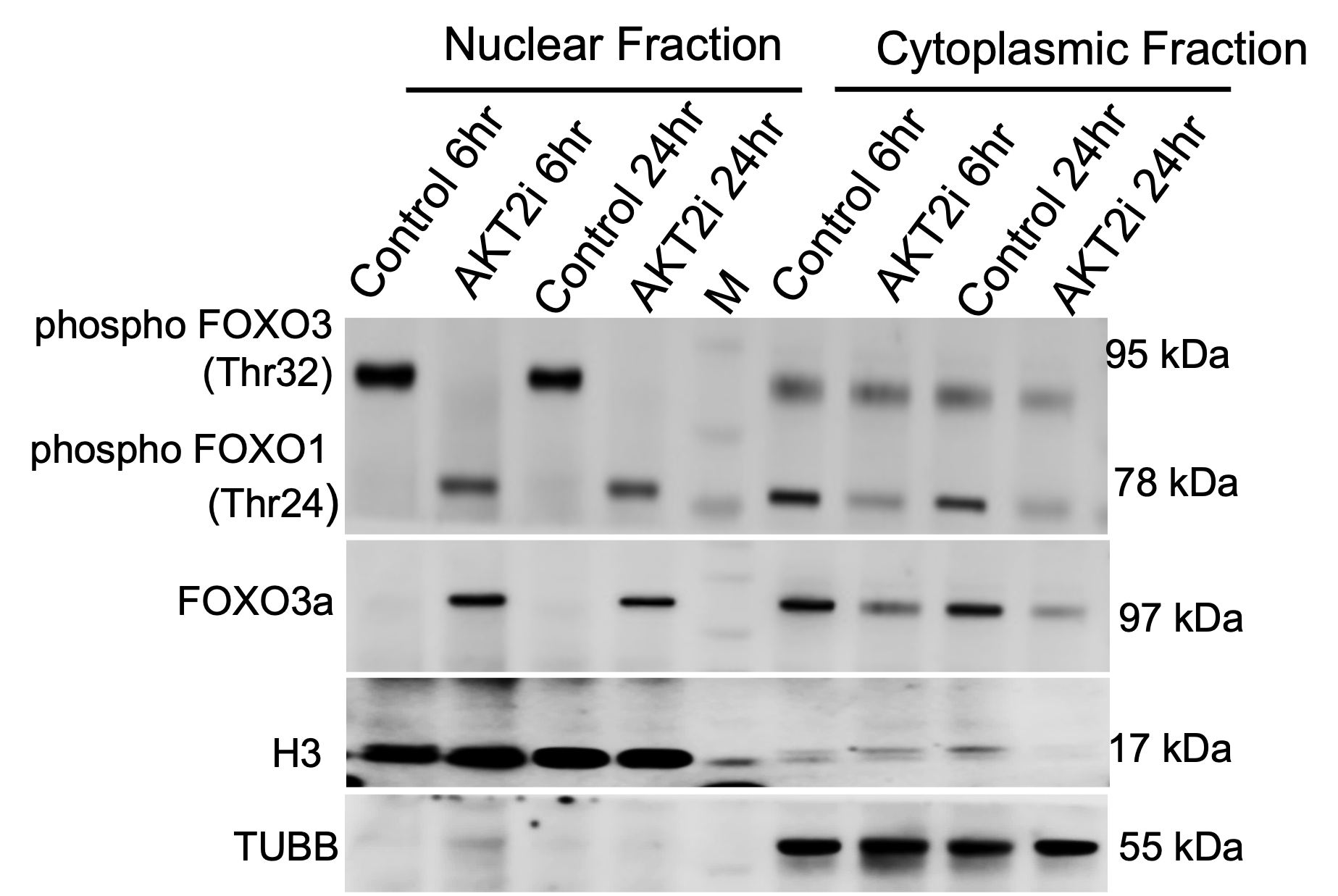

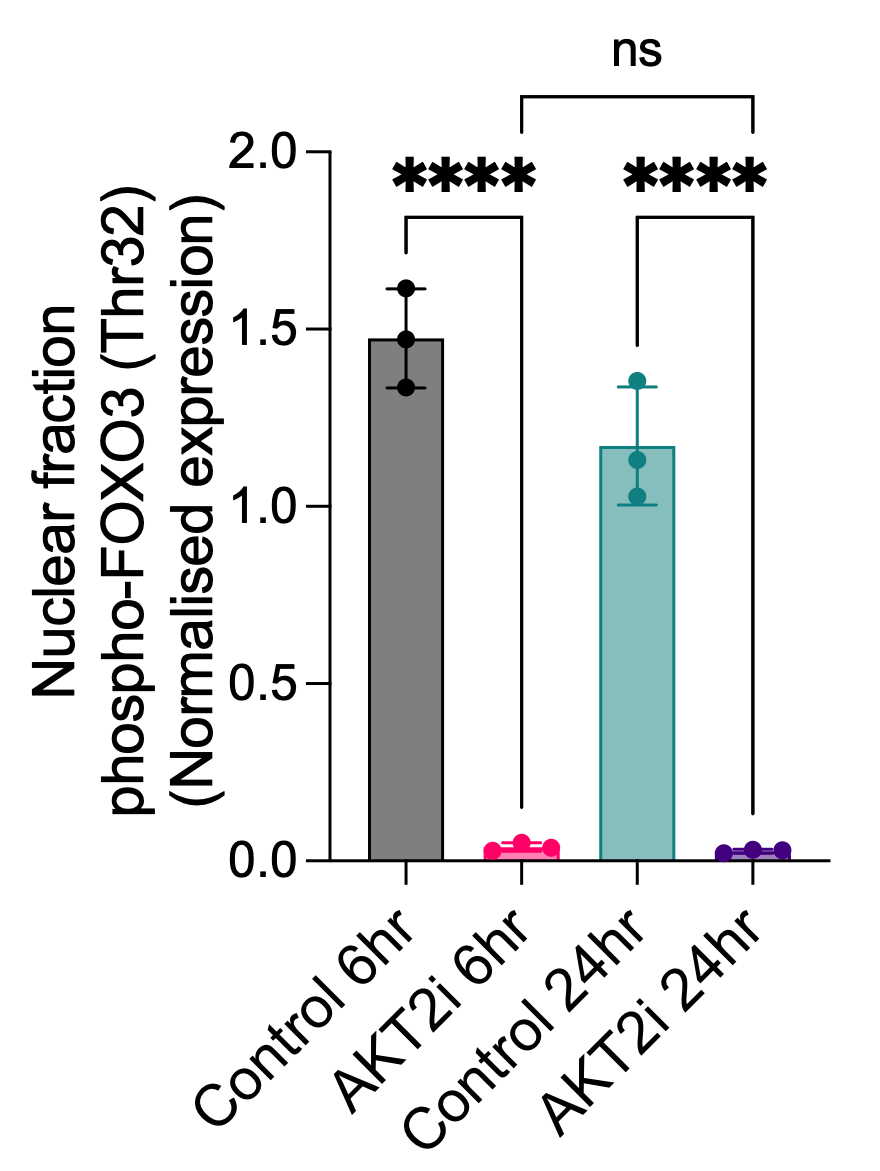

**Figure S3. Compensatory AKT1 activation facilitates TERT and FOXO3a nuclear localization in RPE.** (**A**) RT-PCR analysis shows significant upregulation of *Tert* gene expression in the 15-month-old RPE of *Akt2* cKO mice, whereas a significant downregulation of Tert expression is observed in the age matched RPE of *Akt2* KI and *Akt2* FL mice. n=3**.** (**B**) A standard curve and linear relationship for Q-TRAP were established, with the Ct values (± standard deviation) of HeLa lysates plotted against log[protein] to derive the linear equation. The slope and Y-intercept values from this equation are subsequently used to quantify the relative telomerase activity (RTA) of the RPE samples. (**C**) Telomerase activity measured by Q-TRAP reveals elevated TERT activity in the 15-month-old RPE of *Akt2* cKO mice compared to *Akt2* KI, while low TERT activity is observed in the age matched RPE of *Akt2* KI mice. For Q-TRAP quantification, the RTA of the RPE samples was calculated using the following standard curve and equation obtained from the same Q-TRAP assay: y = −4.0027x + 13.489; RTA = 10^ [(Ct sample−Yint)/slope]. (**D**) A schematic diagram illustrates the phosphorylation sites in TERT, highlighting Ser227 and Ser824 as phosphorylation sites on the AKT1 schematic map. (**E**) Time course analysis of AKT2 inhibition and AKT1 activation from 30 minutes to 24 hours upon treatment with the AKT2 inhibitor CCT128930 at 6 nM in human RPE cells. AKT1 activation at the 6 h time point demonstrates strong TERT phosphorylation at Ser824, which remains stable up to 24 h. 20μg of protein was loaded per condition, and ACTB was used as loading control**.** (**F**) A line plot displays the expression levels of phospho-AKT1 (Ser474), phospho-TERT (Ser824), and phospho-AKT2 (Ser474) following AKT2 inhibition with a 6 nM dose of the inhibitor CCT128930 for 24 h. Treatment at 6 nM resulted in a time-dependent reduction of phospho-AKT2 over 24 h, while compensation and differential expression of phospho-AKT1 and phospho-TERT were observed. Data points represent time intervals from 30 min to 24 h following AKT2 inhibition. (**G**) Western blot showing changes in the expression of phospho-AKT1 (Ser473), phospho-TERT (Ser824), and compensatory AKT2 activation in telomerase-immortalized RPE cells (hTERT RPE) upon selective AKT1 inhibition using A674563 HCl at 11 nM, indicating that AKT1 inhibition can significantly impact TERT expression and AKT2 signaling. 20μg of protein was loaded per condition, and H3 was used as loading control. (**H, I**) Densitometry analysis showed the expression of (**H**) phospho-AKT1 (Ser473) and (**I**) phospho-TERT (Ser824) in hTERT RPE cells following treatment with the AKT1 inhibitor for 6 and 24 h. n=3 (**J**) Relative telomere length was measured using the RHTLQ method in human fetal RPE cells treated with AKT2 and AKT1 inhibitors for 48 h. Neither AKT2 inhibition nor AKT1 inhibition resulted in significant changes in telomere length in human iPSC RPE cells. (**K**) Immunoblot analysis showing the expression levels of phospho-FOXO3a, phospho-Foxo3a and phoshpho-PRAS40 in the RPE of 15-month-old *Akt2* FL, *Akt2* KI, and *Akt2* cKO mice (n=3). Elevated phosphorylation of Foxo3a, which indicates its inactive state in cytoplasm, was observed in the RPE of *Akt2* KI mice compared to *Akt2* cKO and wild-type controls. A total of 10 µg of protein was loaded in each well, and Vinculin was used as the loading control. n=3. (**L**) Densitometry analysis shows the normalized expression of phospho-FOXO3a in the RPE of 15-month-old *Akt2* FL, *Akt2* KI and *Akt2* cKO mice (n=3). (**M**) Western blot analysis shows the subcellular localization of phospho-FOXO3a and total FOXO3a in iPSC RPE *CFH* Y402H cells. In DMSO-treated controls, phospho-FOXO3a is primarily nuclear, indicating its degradation, while dephosphorylated FOXO3a suggests an active form. Total FOXO3a levels increase in the nucleus and reduce in the cytoplasm after AKT2 inhibition. A total of 25 µg of protein was loaded in each well. TUBB was utilized as a control for the cytoplasmic fraction, while H3 served as a nuclear loading control. (**N**) Densitometry analysis of phospho-FOXO3a/FOXO3a in iPSC RPE *CFH* Y402H cells following AKT2 inhibitor treatment for 6 and 24 h. n=3. Bars represent mean ± SD (n = 3); Statistical significance was determined by one-way ANOVA with post hoc test. ***p* < 0.01, ****p* < 0.001, *****p* < 0.0001, ####*p* < 0.0001. ns= not significant.

A

B

C

D

E

F

P

I

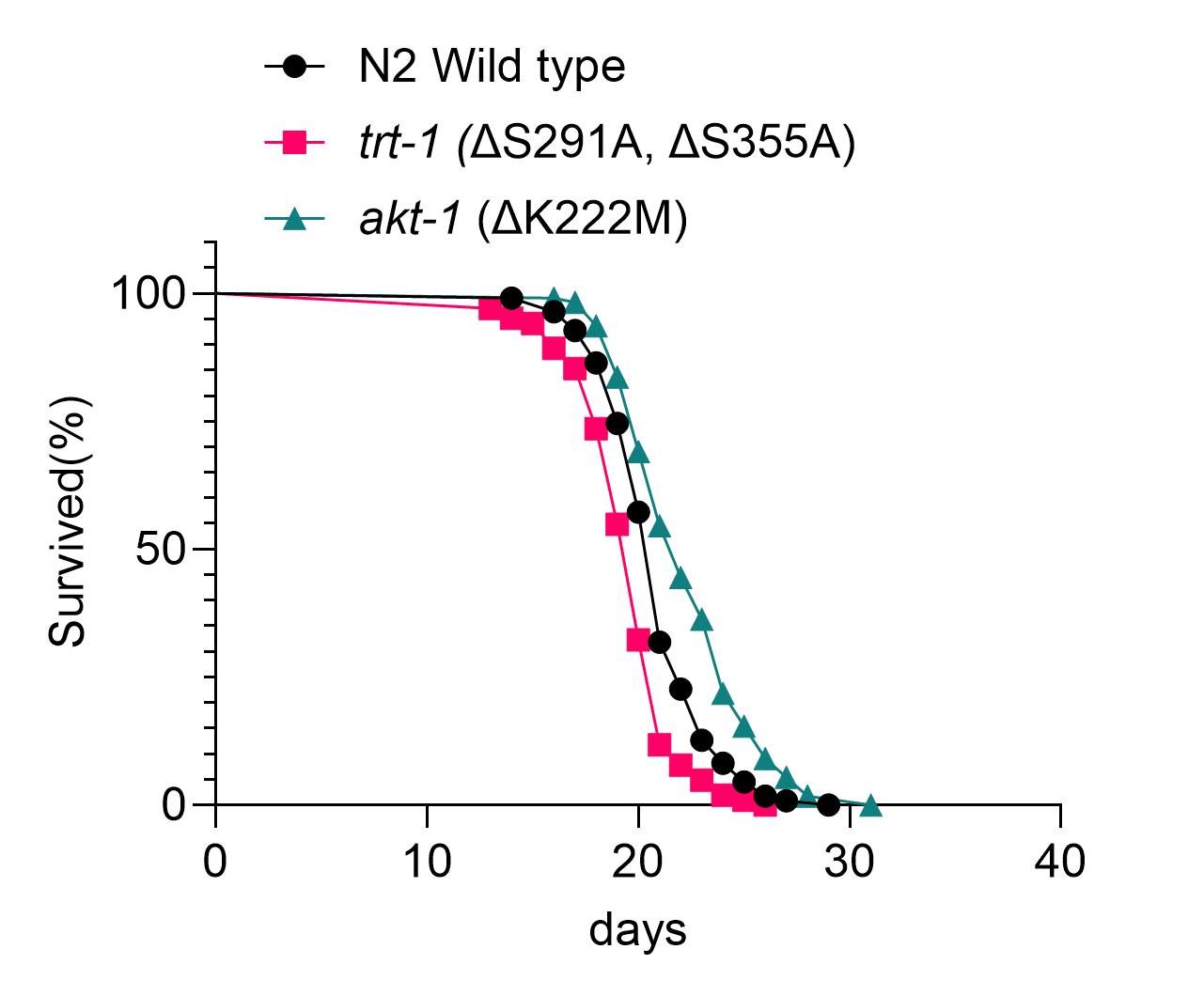

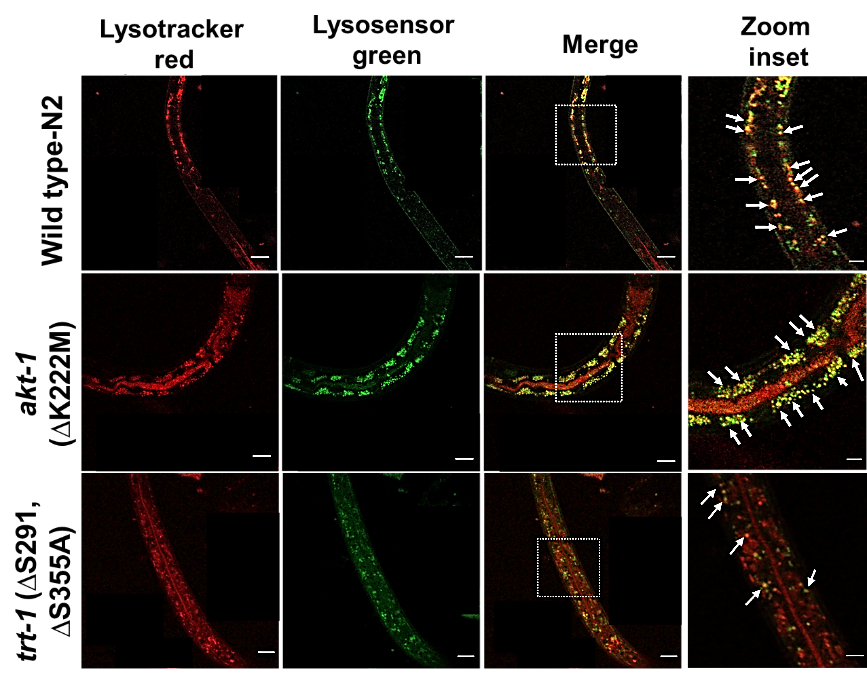

J

K

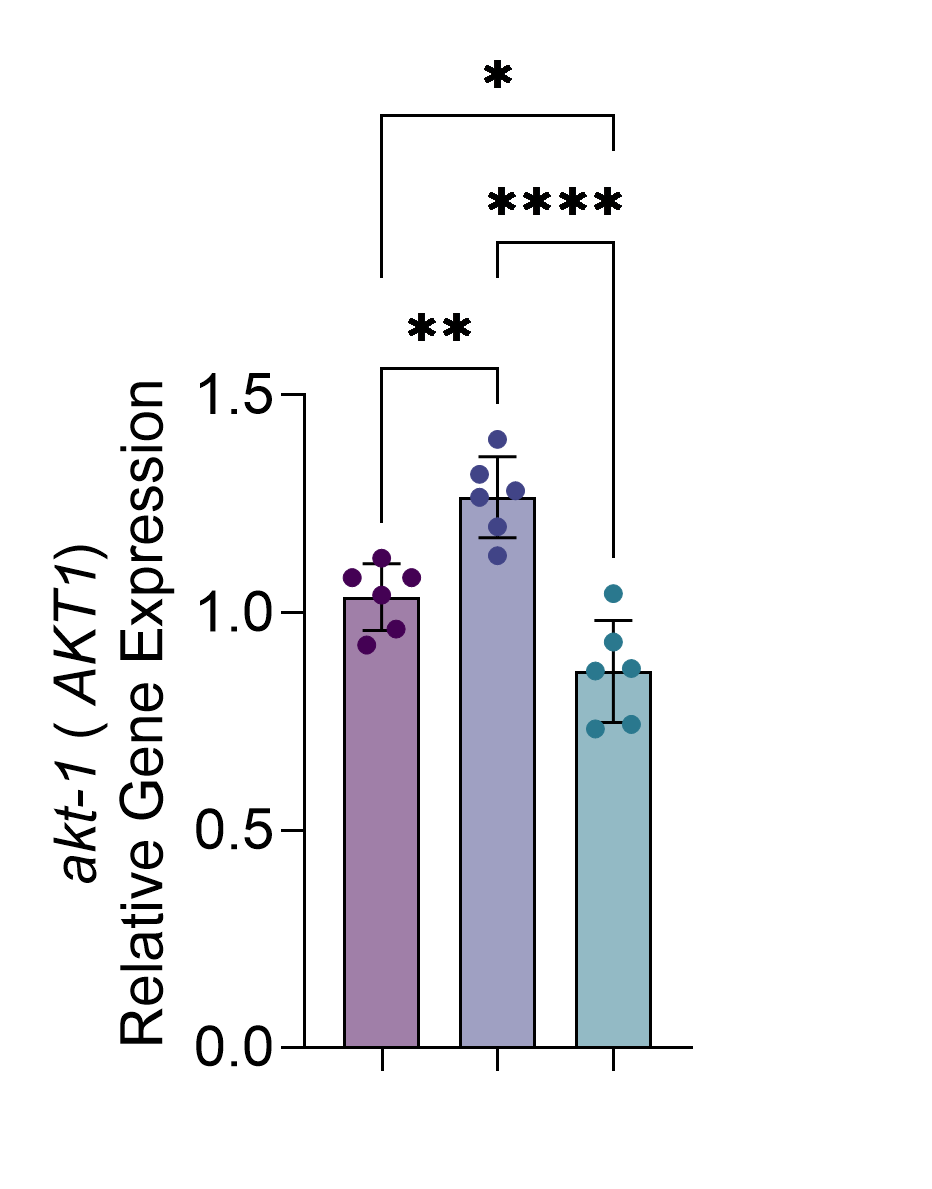

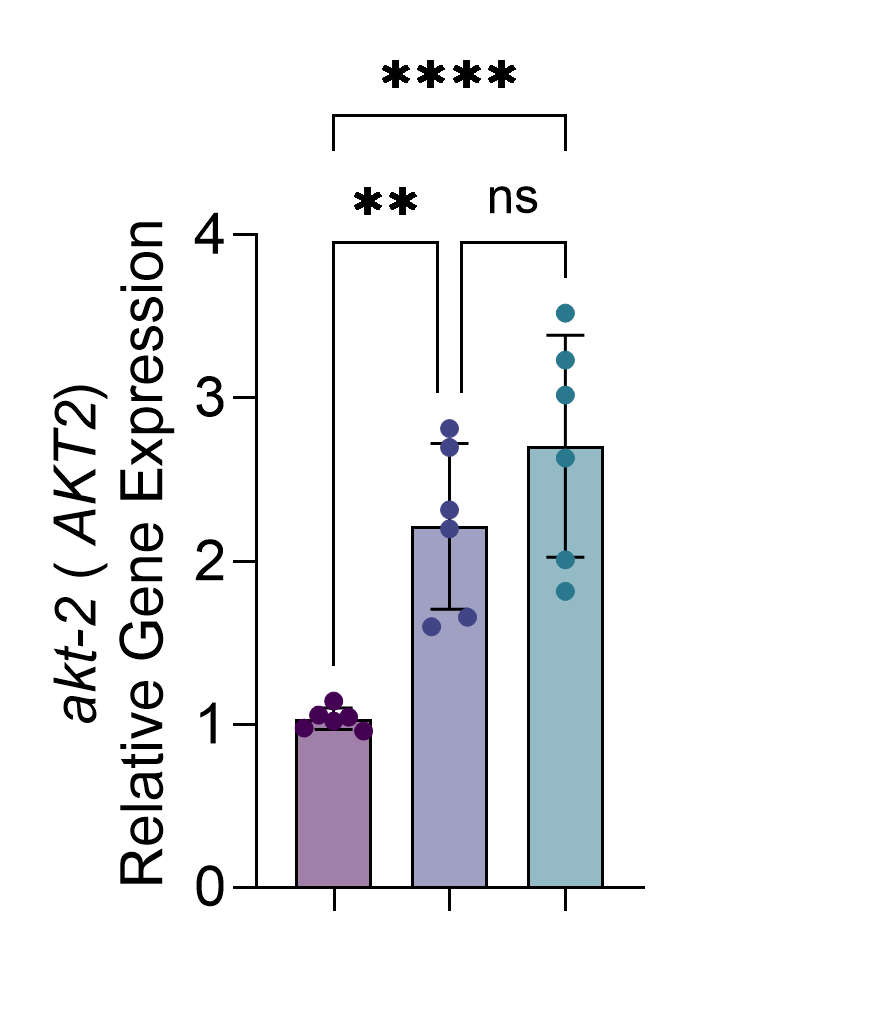

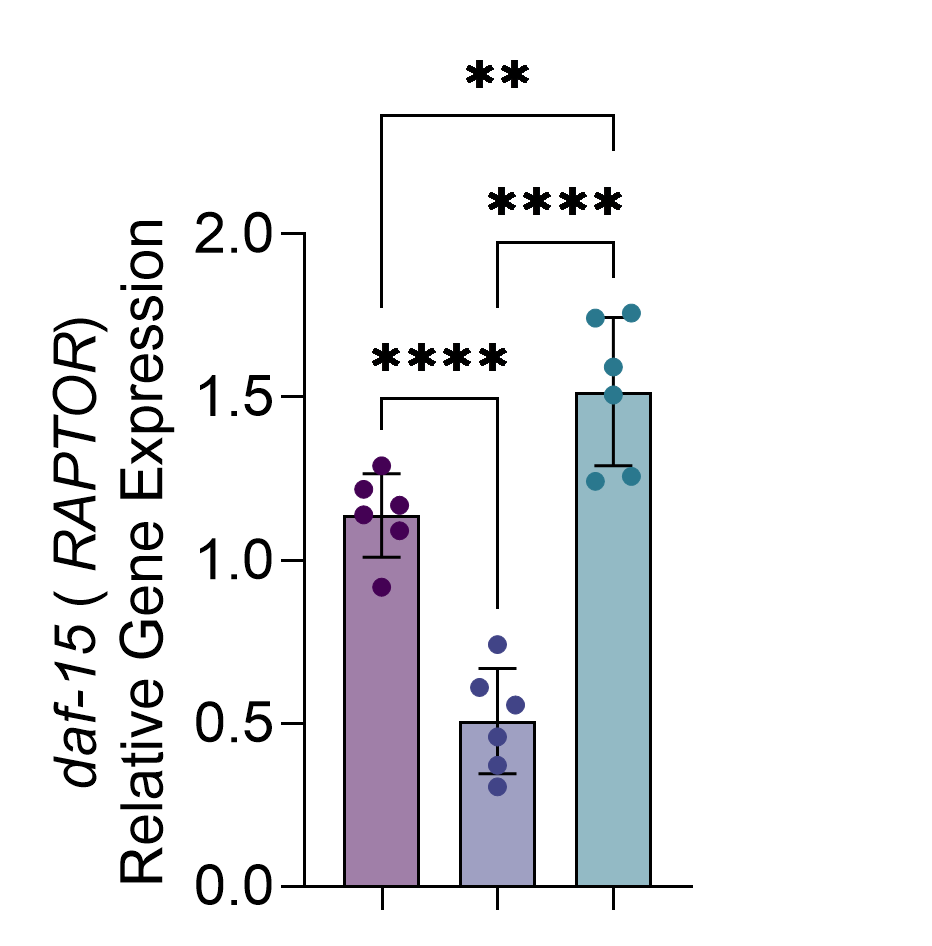

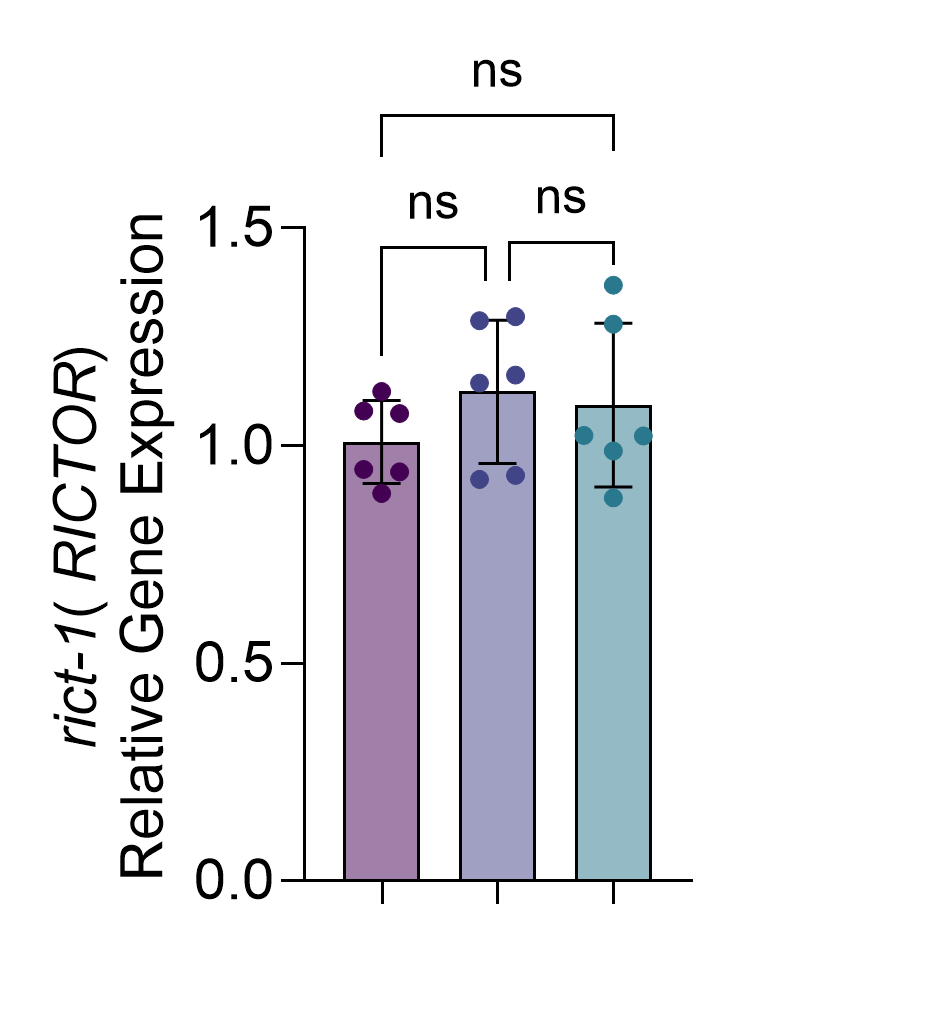

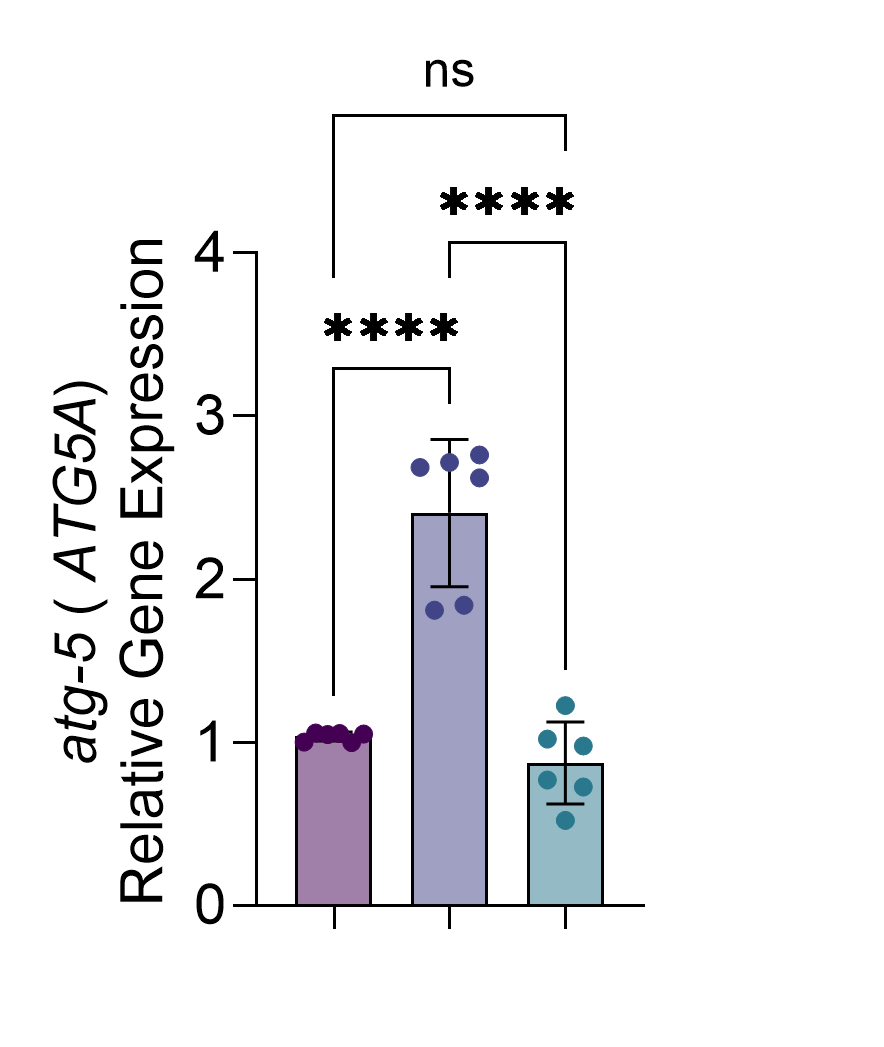

G

H

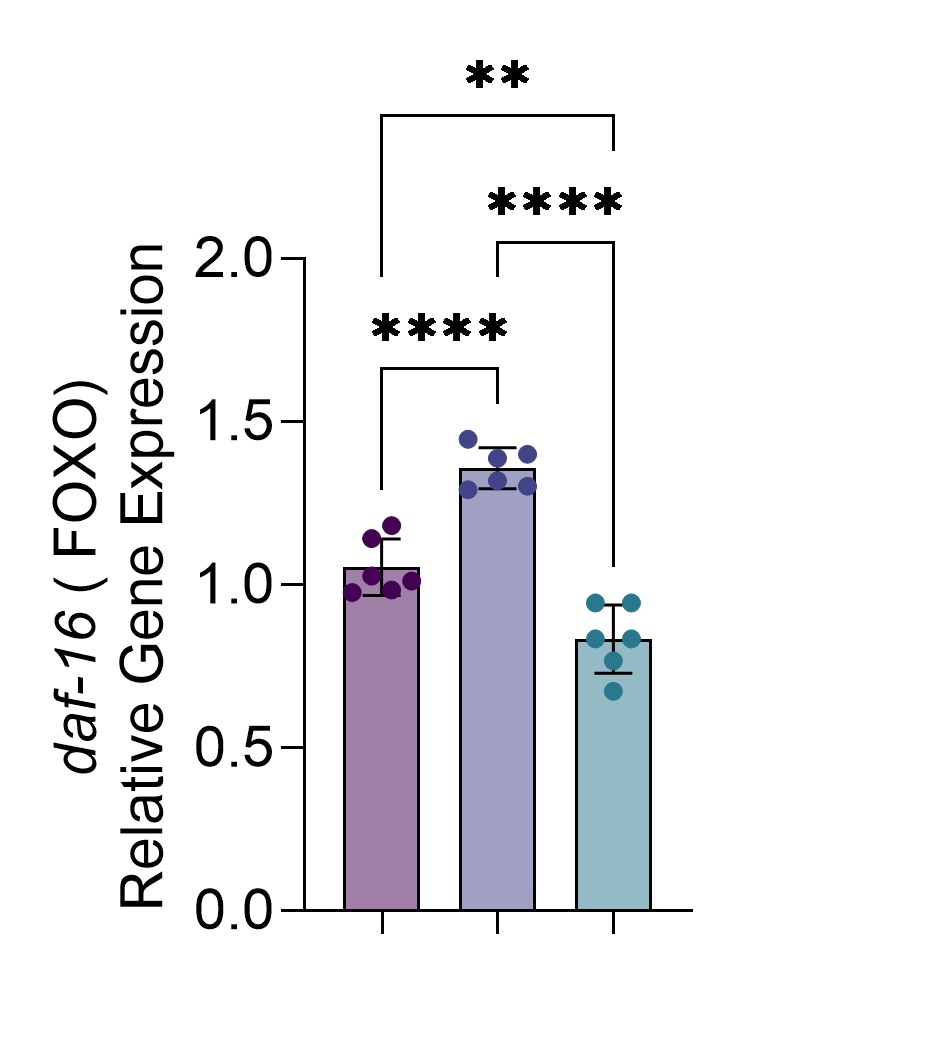

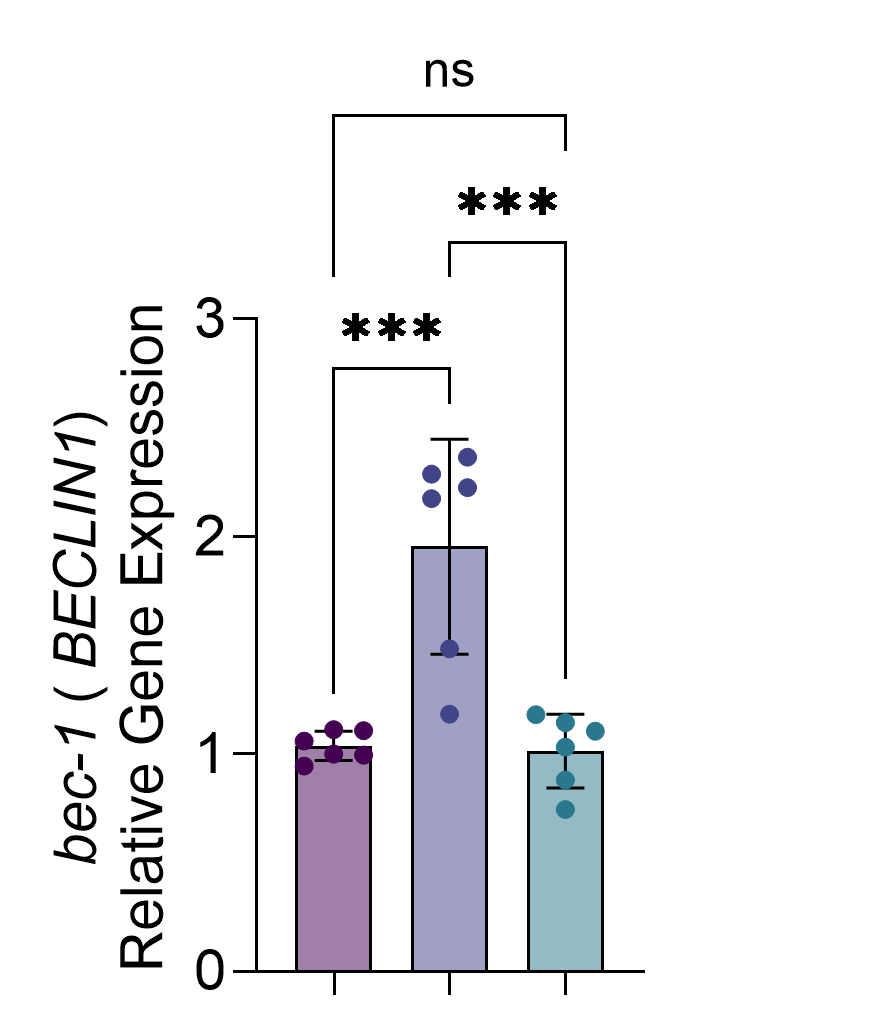

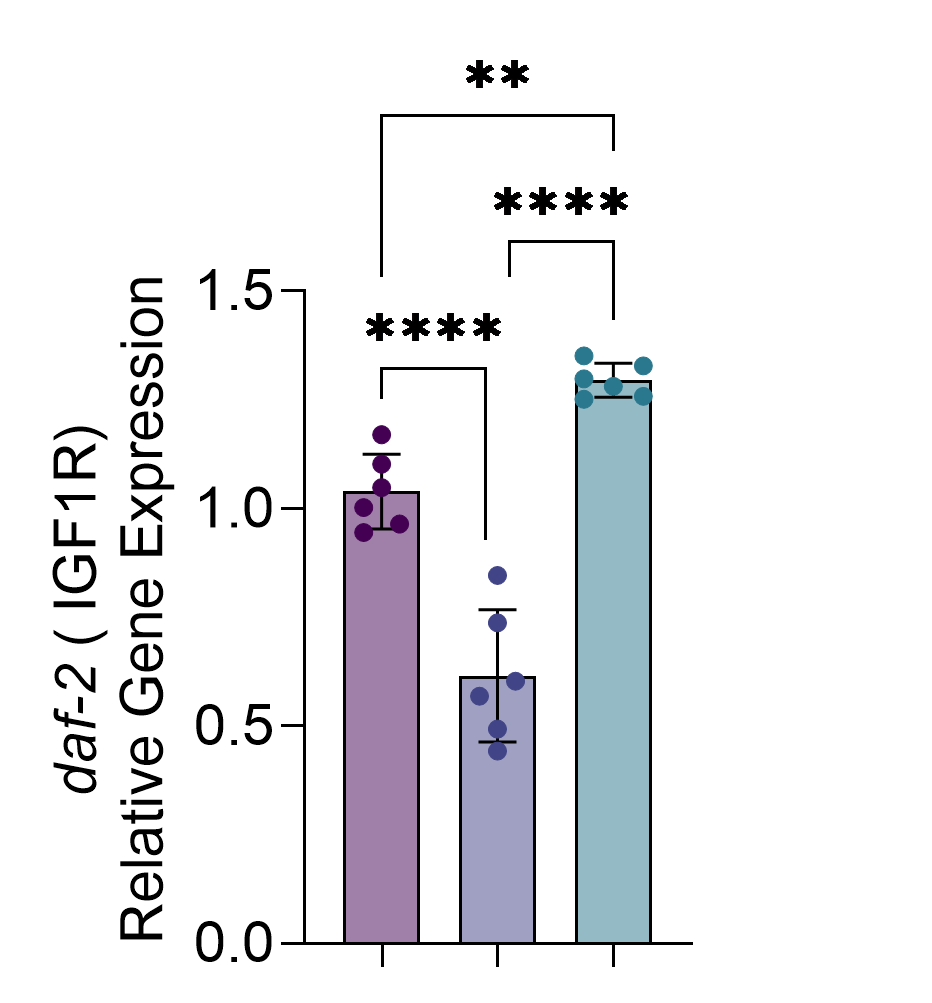

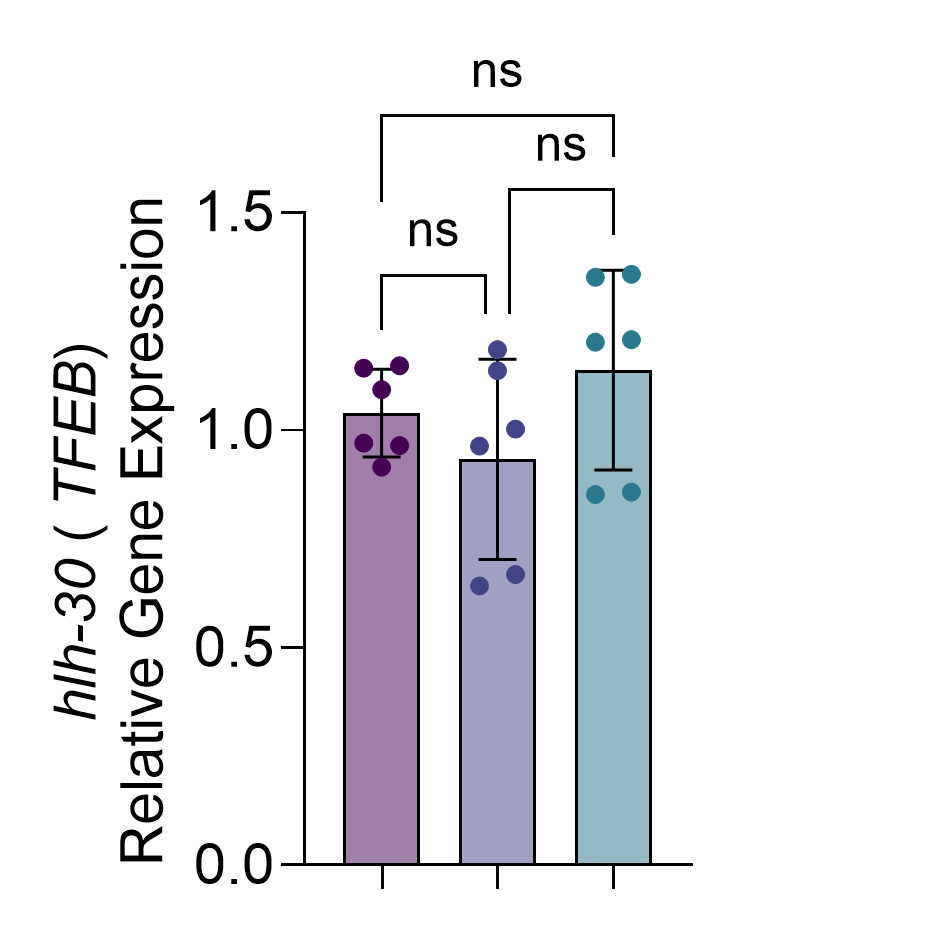

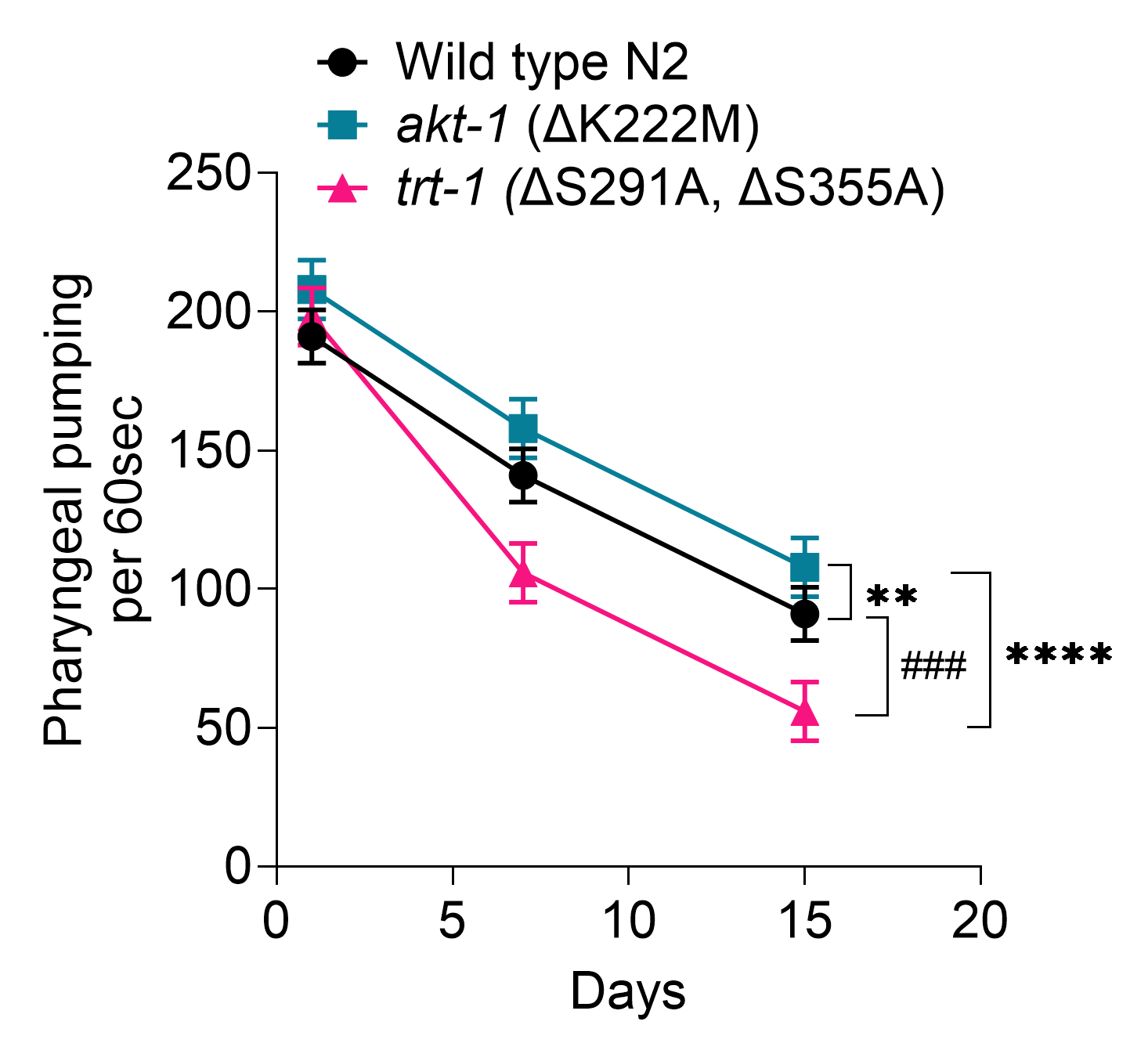

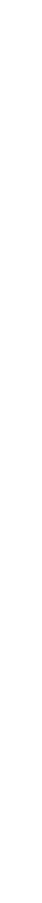

L

M

N

O

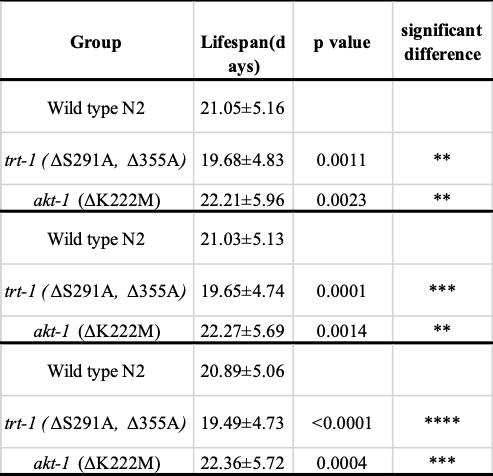

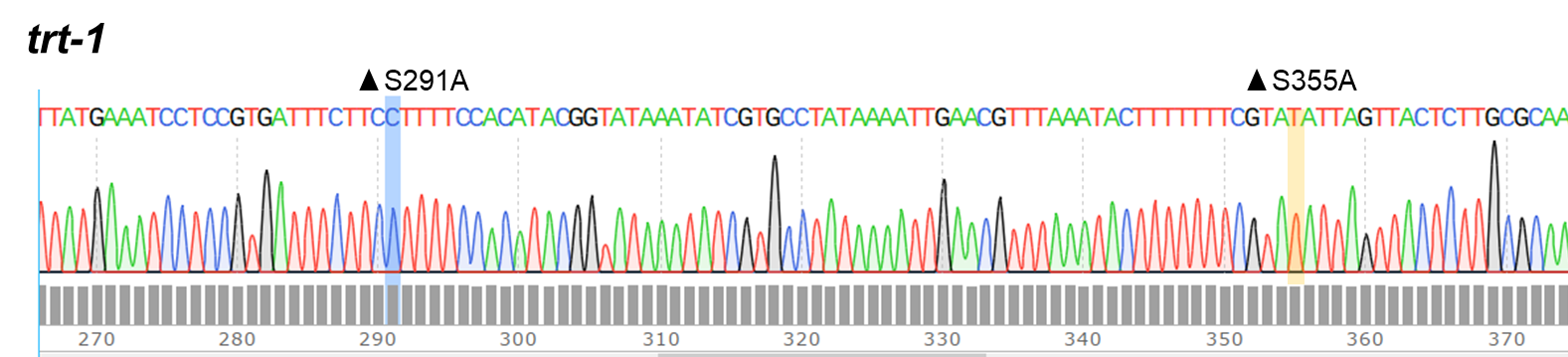

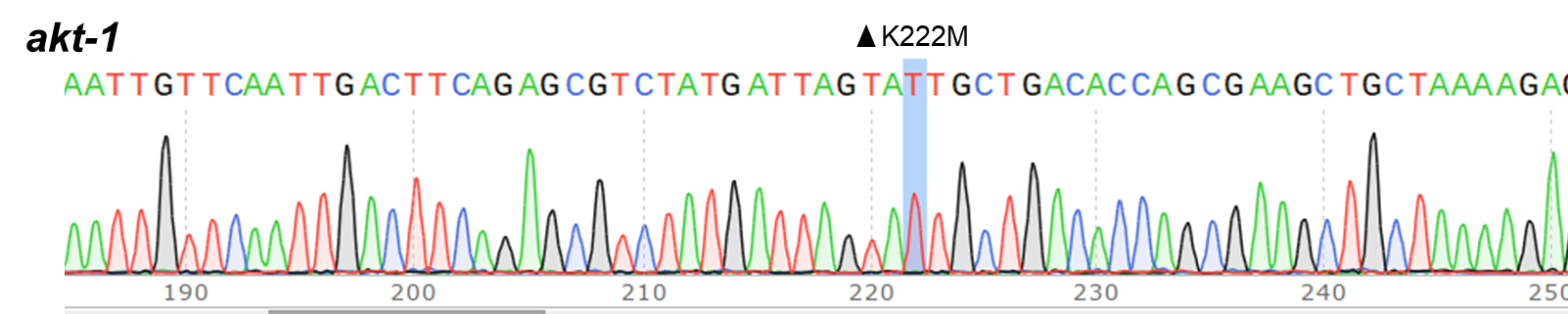

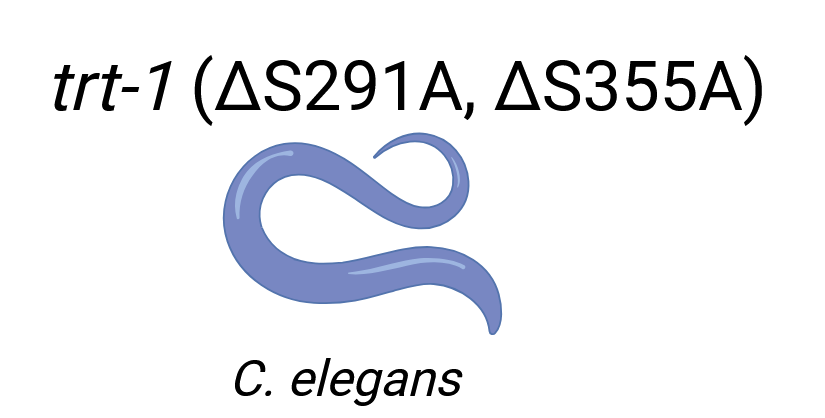

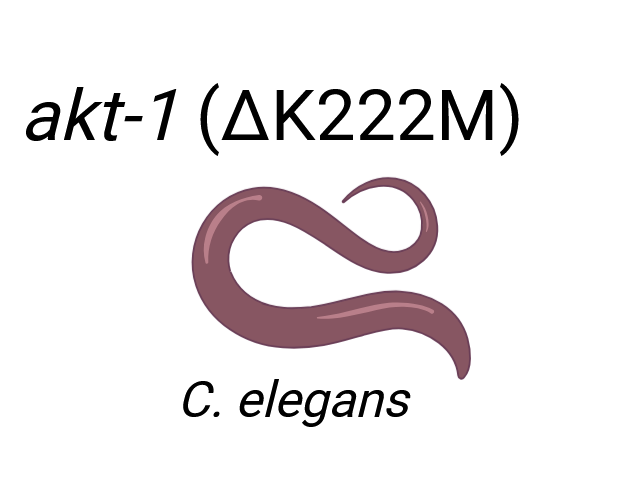

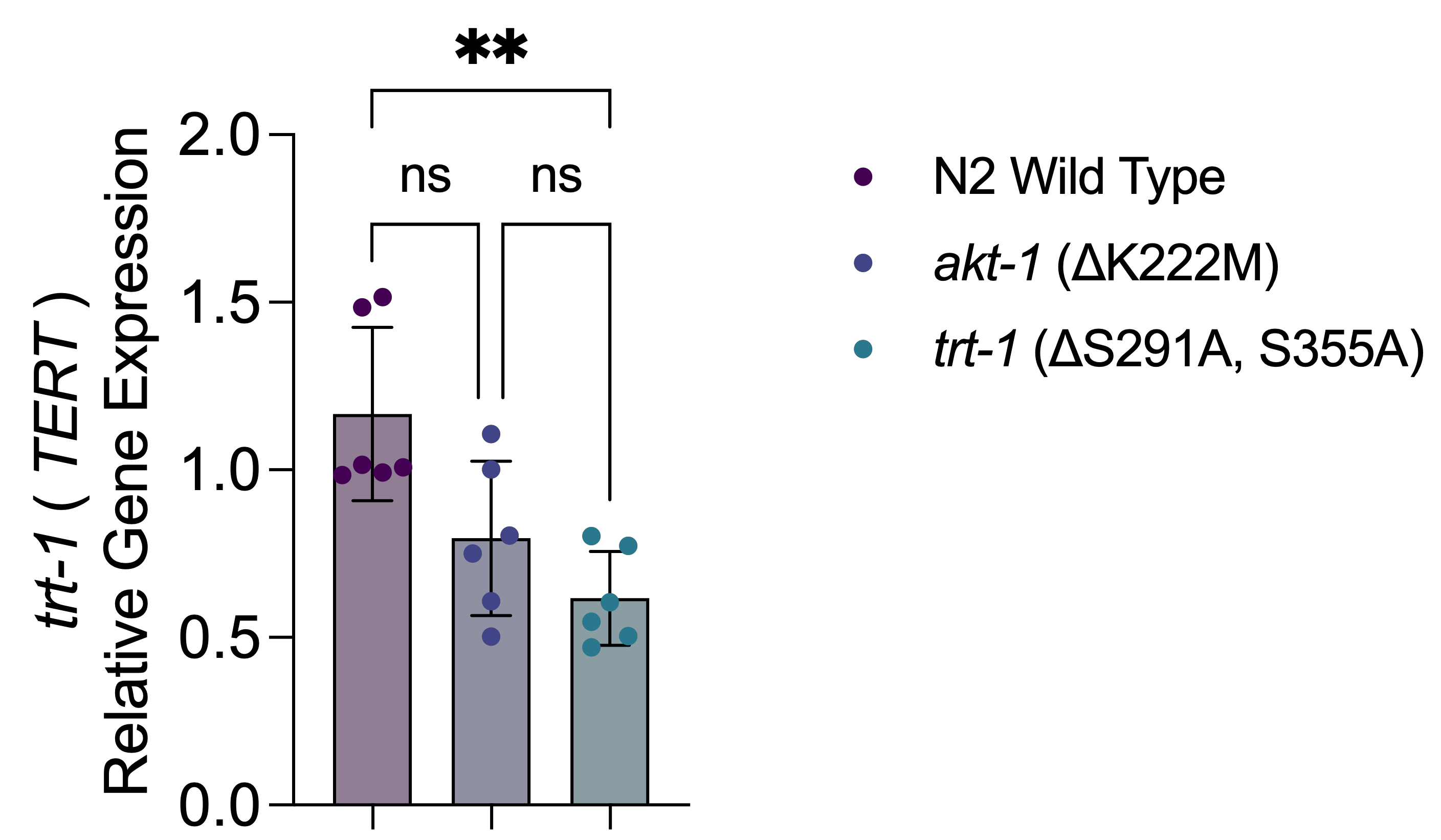

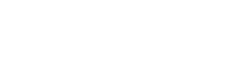

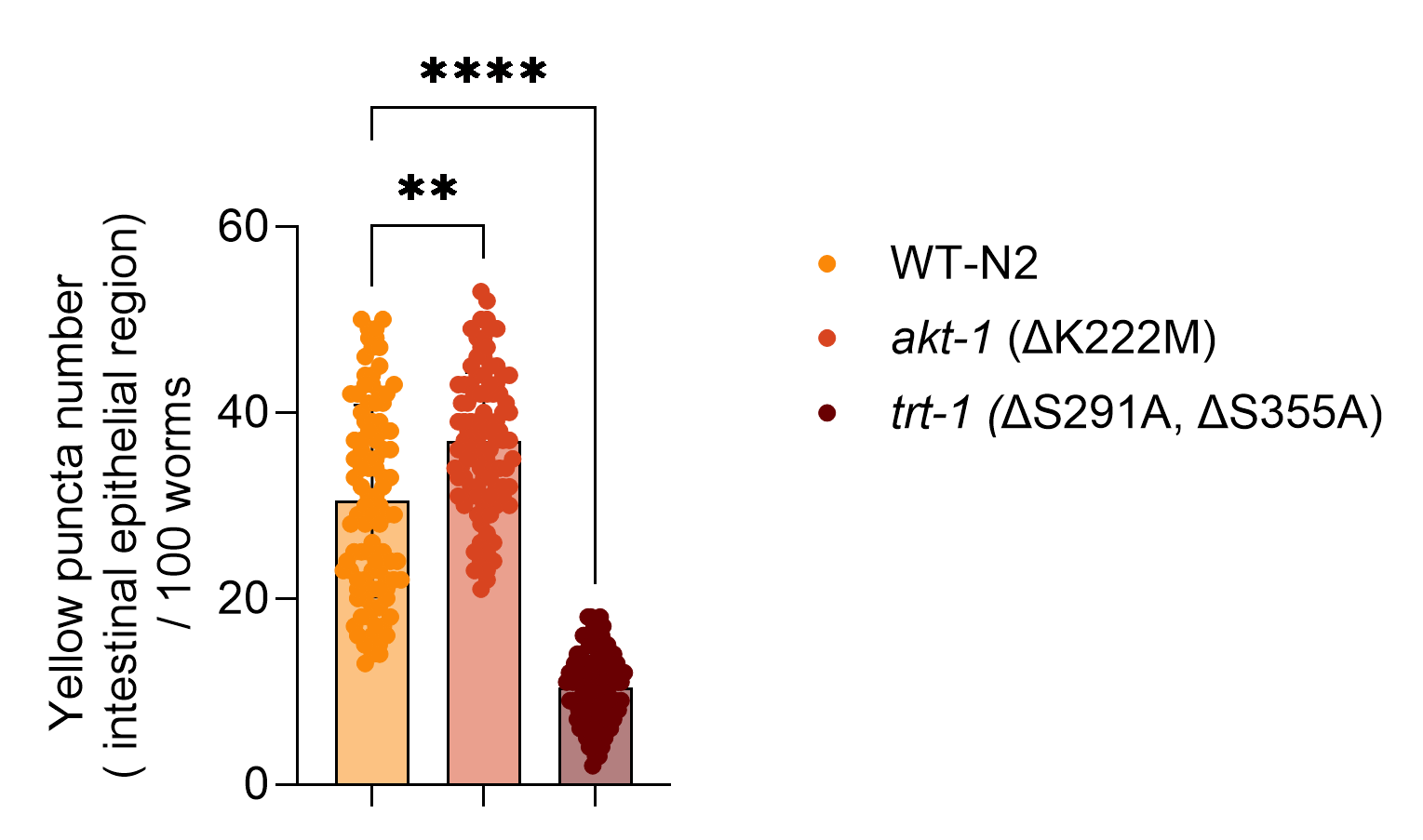

**Figure S4. Loss of AKT1 phosphorylation in *trt-1* mutants impairs lysosomes and decreases survival in *C. elegans.***

(**A**) Sanger sequencing shows the mutational profile of *C. elegans* involving a) the *trt-1* variant with ΔS291A and ΔS355A mutations, and b) the *akt-1* variant with the ΔK222M mutation. (**B–K**) Quantitative PCR analysis of (**B**) *trt-1*, (**C**) *akt-1*, (**D**) *akt-2*, (**E**) *daf-16*, (**F**) *daf-2*, (**G**) *daf-15*, (**H**) *rict-1*, (**I**) *atg-5*, (**J**) *bec-1*, and (**K**) *hlh-30* in the indicated genotypes. Expression levels are normalized to *act-1* (Actin) as internal control. Data represent mean ± SEM from n = 3 biological replicates. (**L**) Confocal live imaging demonstrates the distribution of functional lysosomes in intestinal epithelial region in the *akt-1* kinase-dead and *trt-1* mutants, visualized using LysoTracker red and LysoSensor green dye. LysoTracker red highlights active lysosomes, while LysoSensor green dye indicates lysosomal pH changes, providing insights into lysosomal function and integrity. Images reveal altered lysosomal distribution and functionality in mutant strains compared to control groups, emphasizing the impact of AKT-1 kinase activity on lysosomal dynamics. *trt-1* variants show low distribution of functional lysosomes compared to *akt-1* variants. Scale bar =25μm. (**M**) Quantification of red and yellow puncta reveals significant differences in lysosomal distribution and functionality between mutant strains and wild type groups, highlighting the role of AKT1 kinase activity in regulating lysosomal dynamics. Each dot represents an individual animal. Horizontal lines indicate mean ± SD. n=100 per group. Statistical significance was determined by one-way ANOVA with Tukey’s post-hoc test. (**N**) Survival line plots show health span of *C. elegans* measured by pharyngeal pumping for 60 s in synchronized L4-stage worms at Days 1, 7 and 15. The *trt-1* mutants shows reduced pumping rate and vitality compared to significantly increased pumping rate displayed by *akt-1* kinase dead mutant. (**O**) Survival line plot shows percentage lifespan of *C. elegans* variants measured for 30 d. The *akt-1* kinase dead mutant shows increased lifespan compared to reduced lifespan in *trt-1* mutants (**P**) Table shows mean life span of each strain in three independent experiments. Statistical significance was determined by one-way ANOVA and Tukey’s post-hoc test; **p* < 0.05, ***p* < 0.01, *** *p* <0.001, **** *p* <0.0001, ^###^*p* <0.001, ns=not significant.

A

B

C

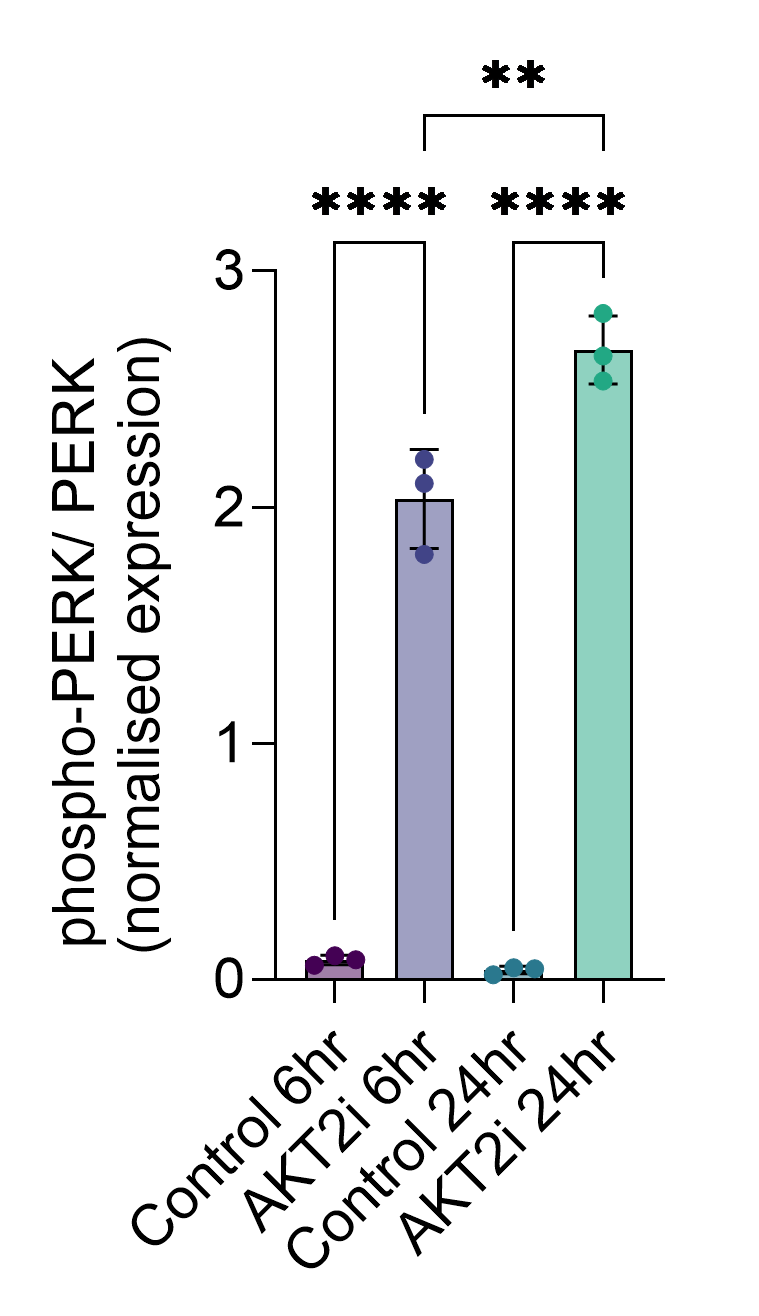

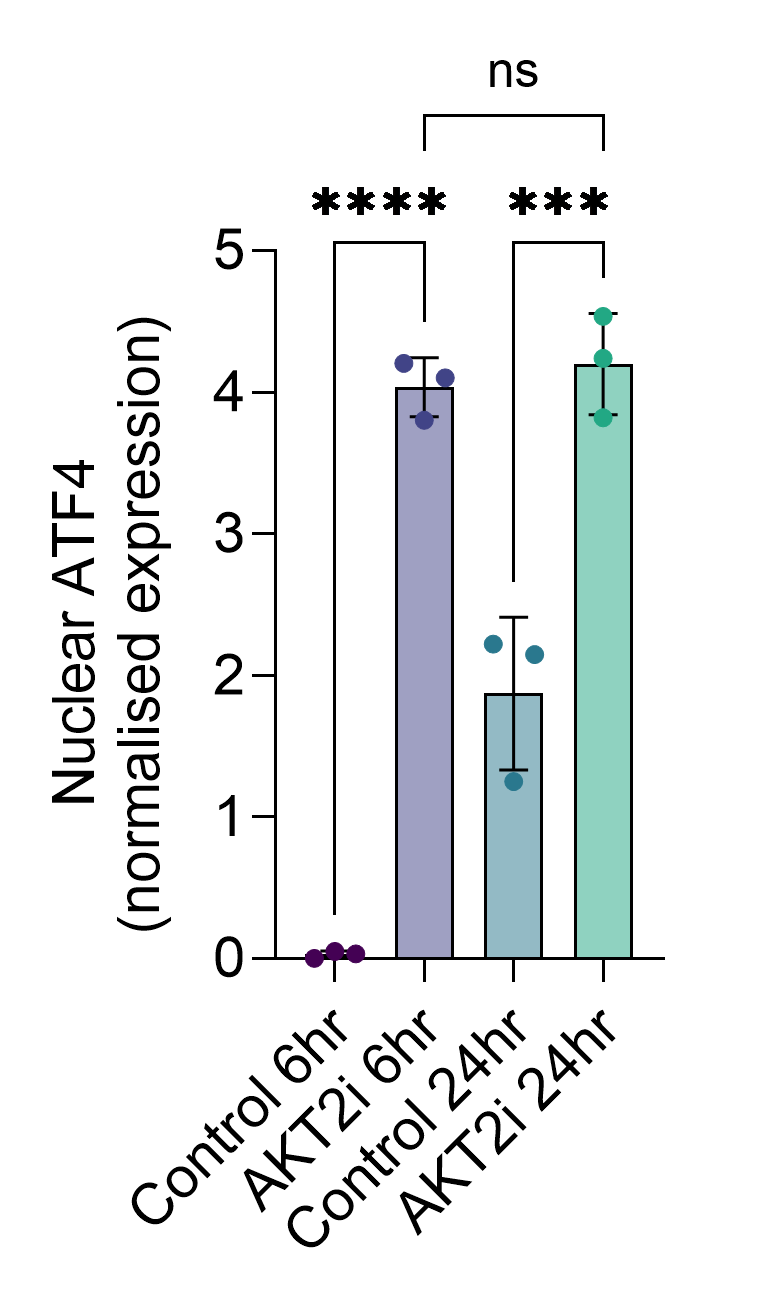

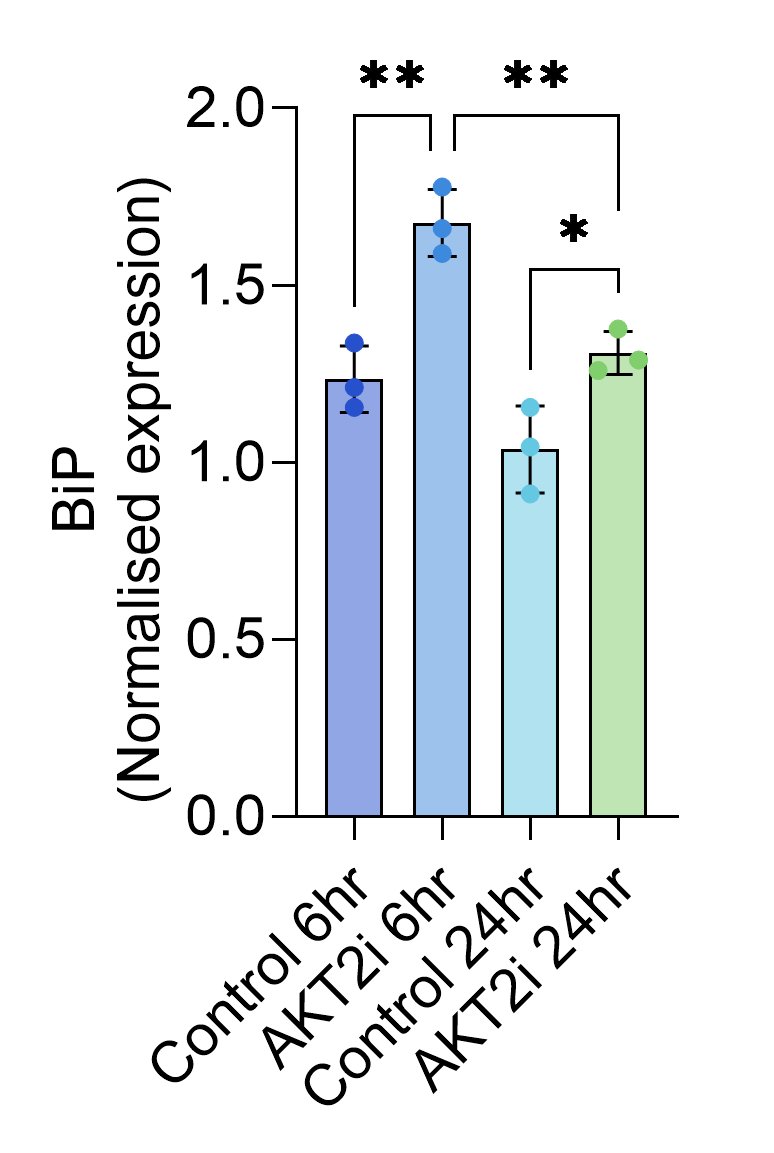

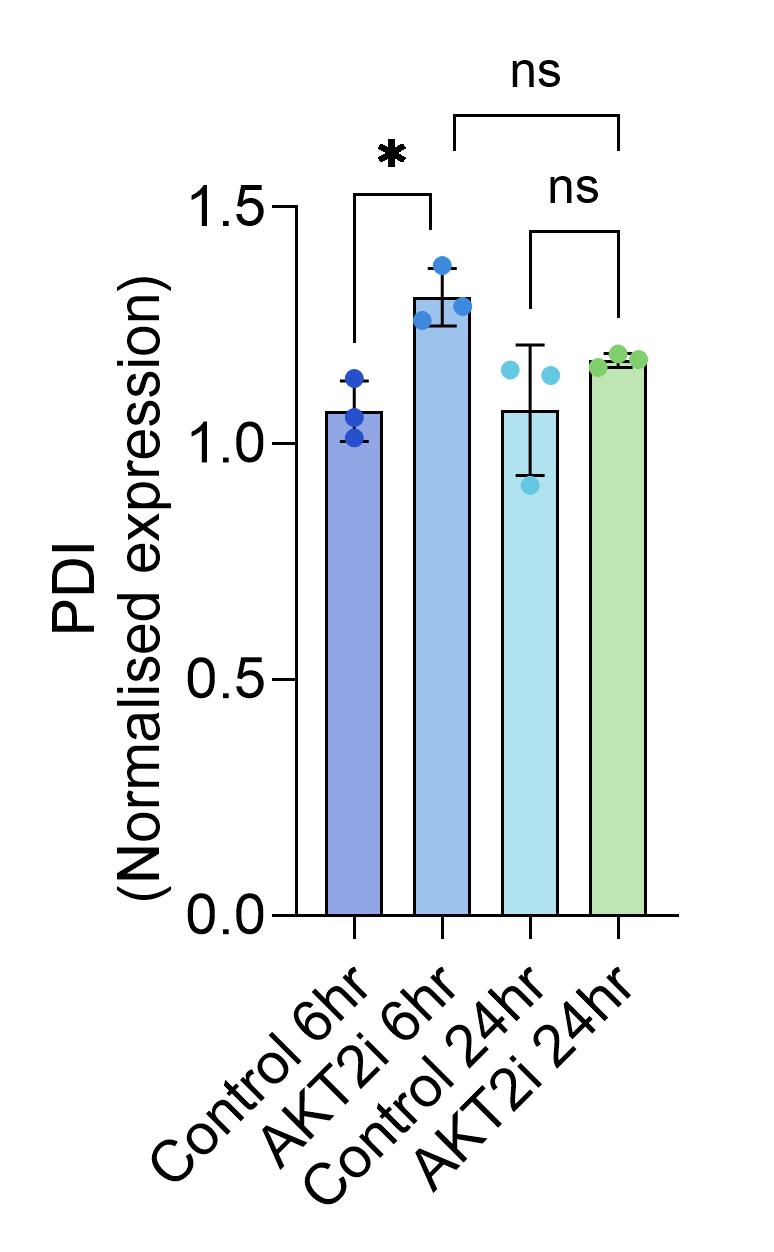

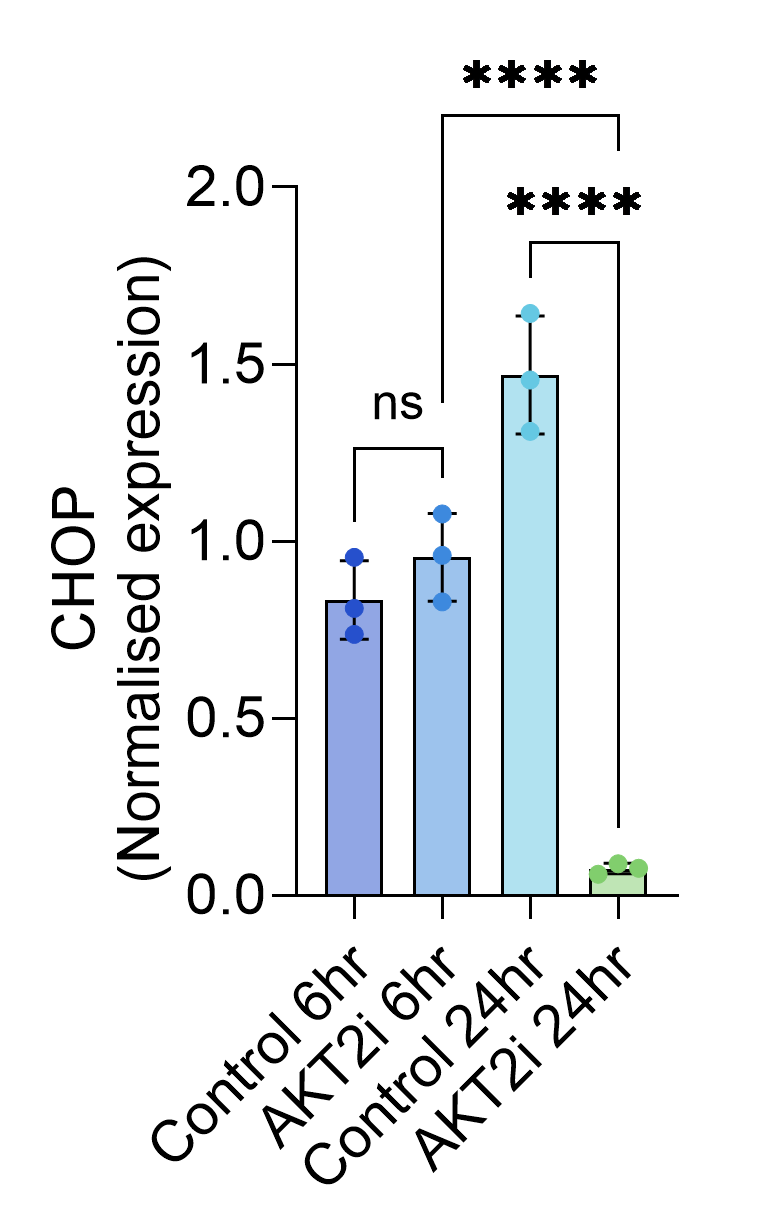

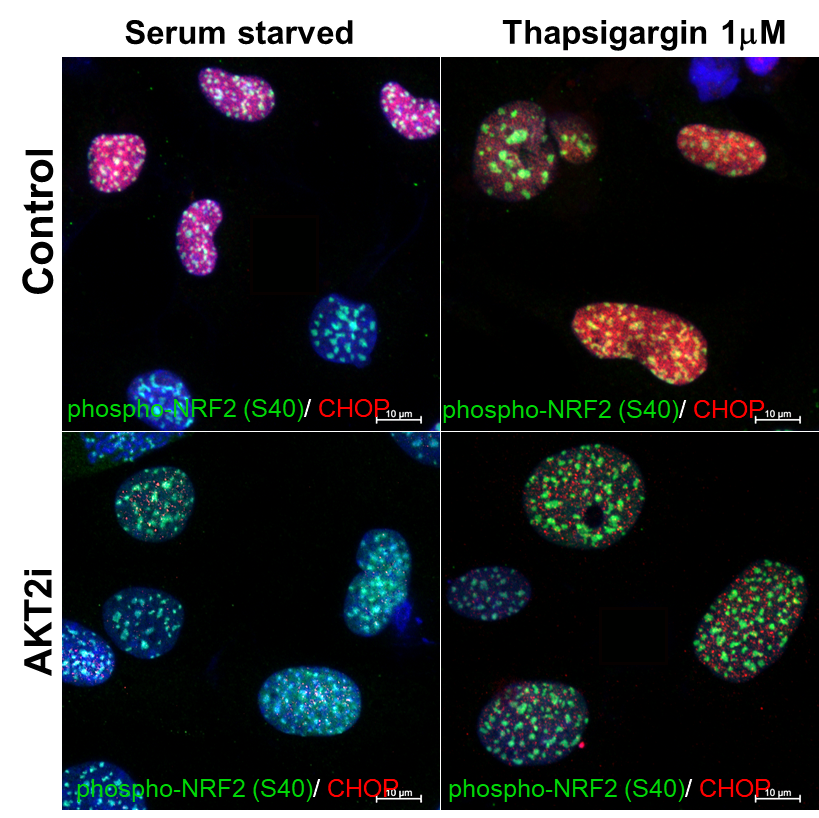

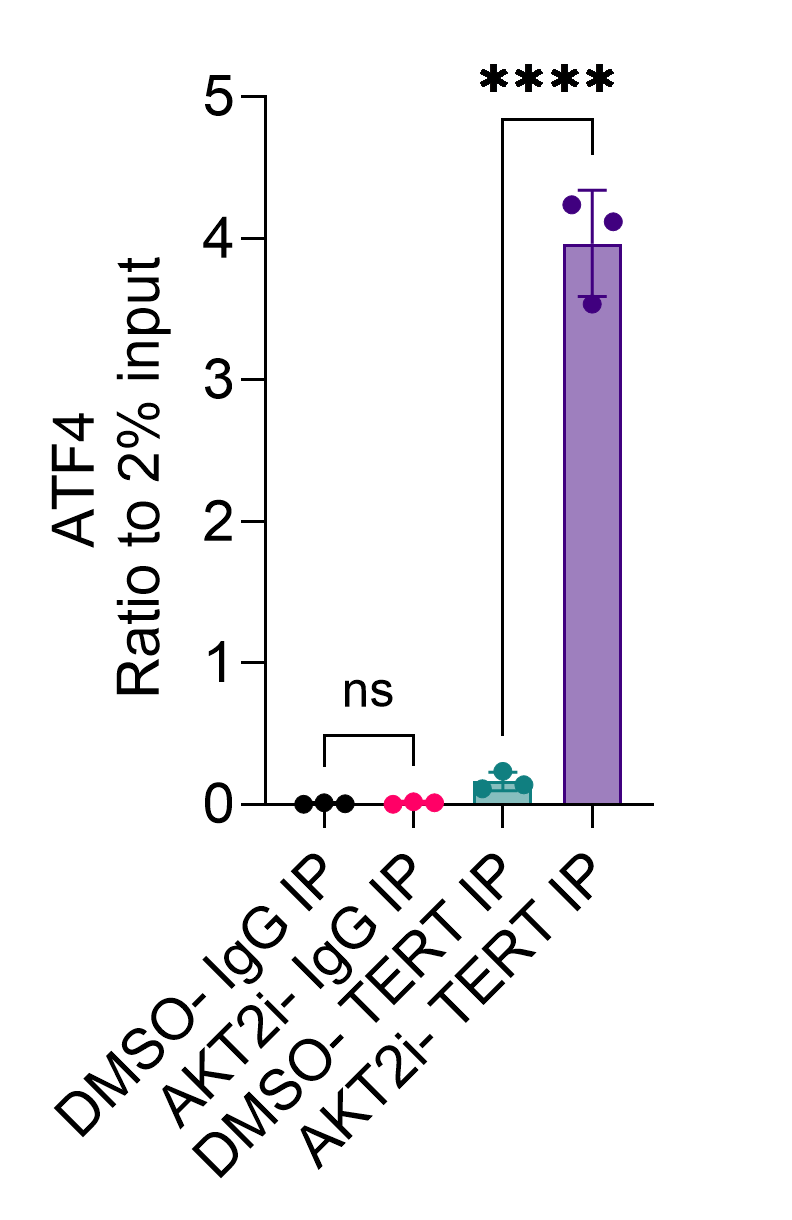

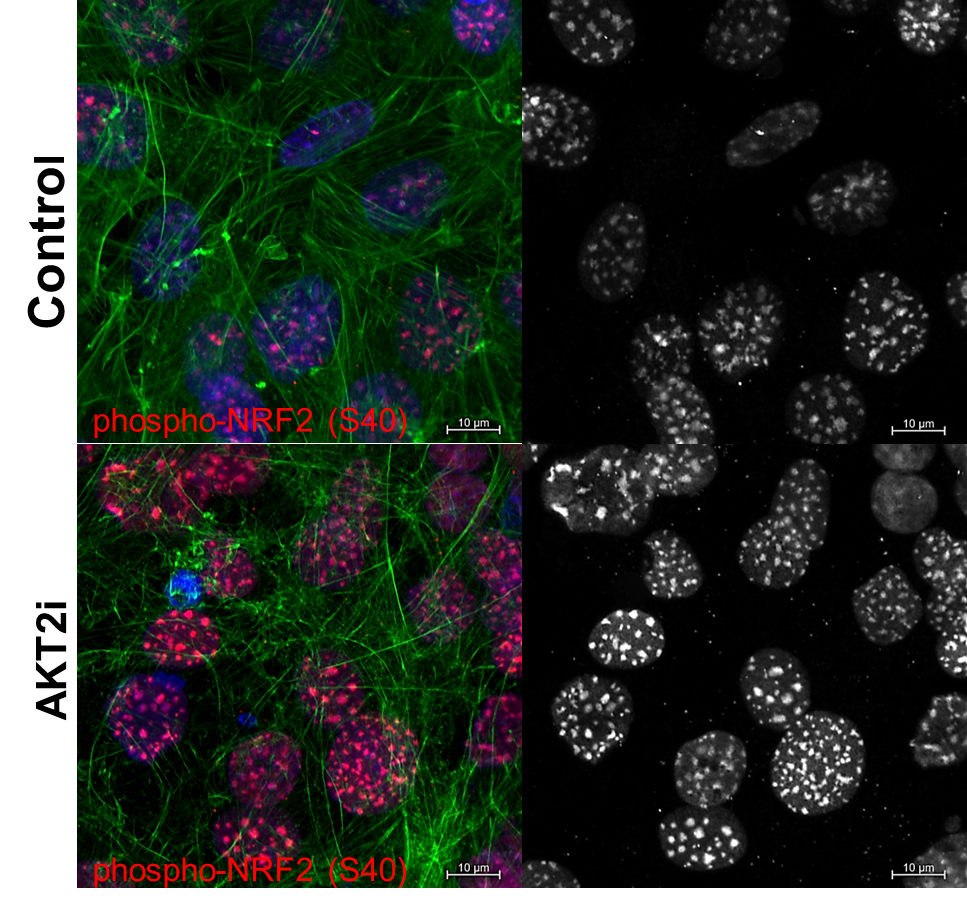

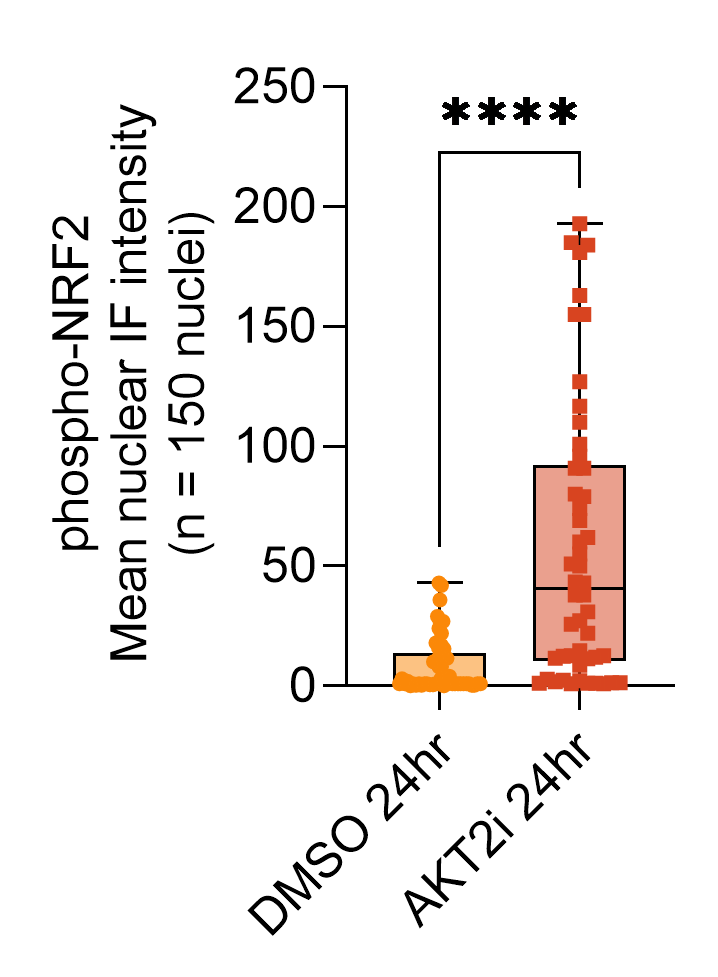

D

E

F

G

H

I

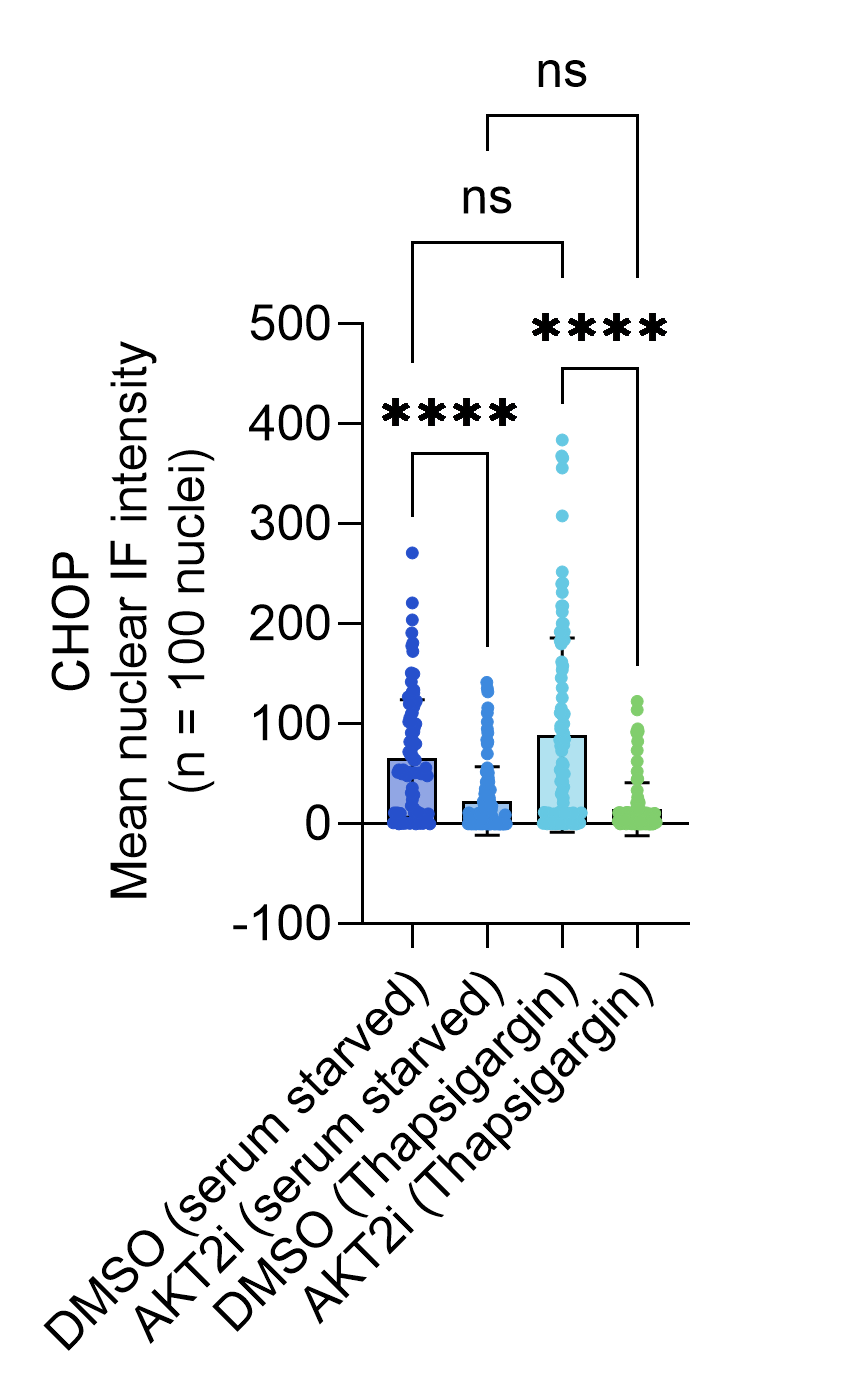

J

K

L

M

N

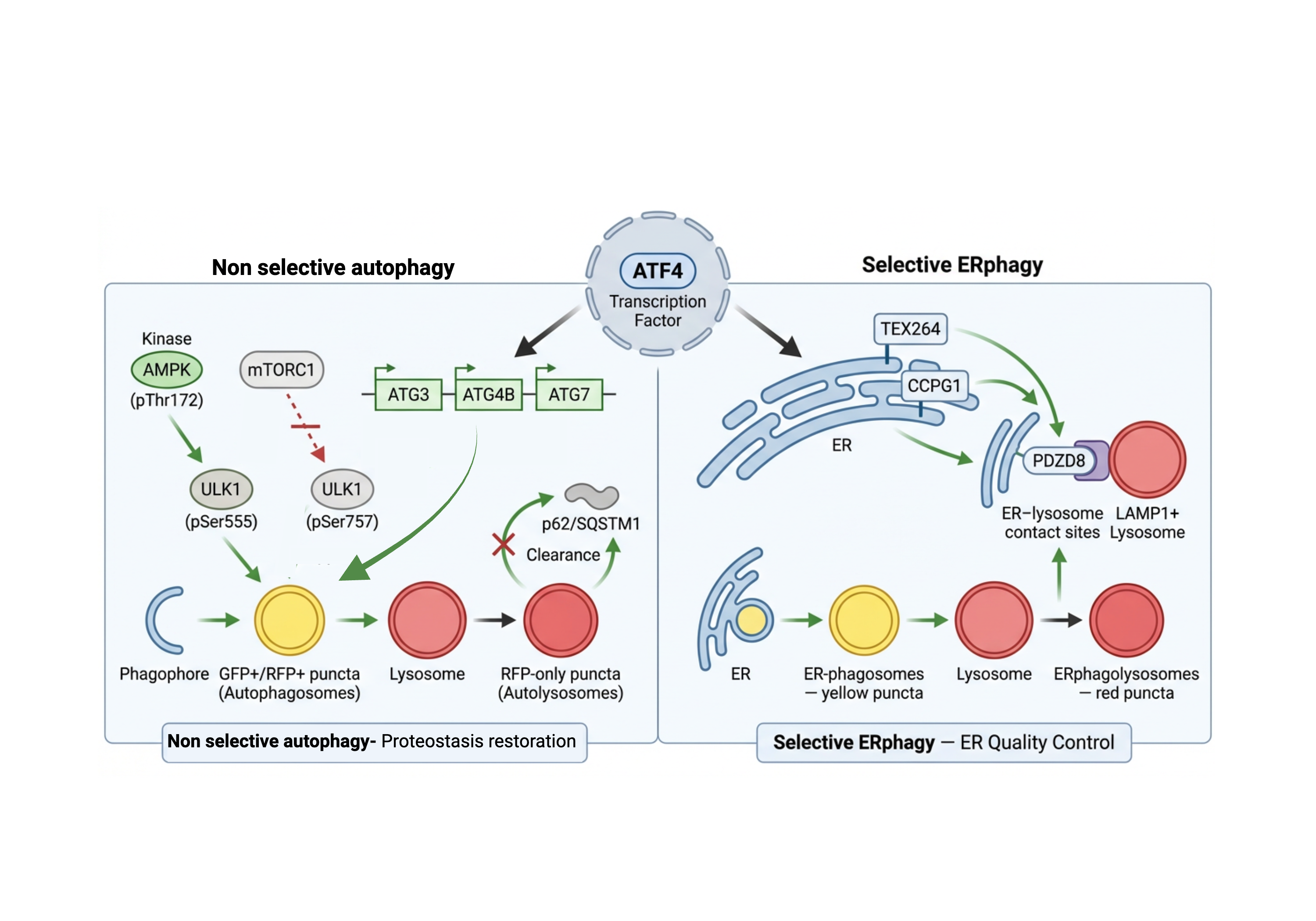

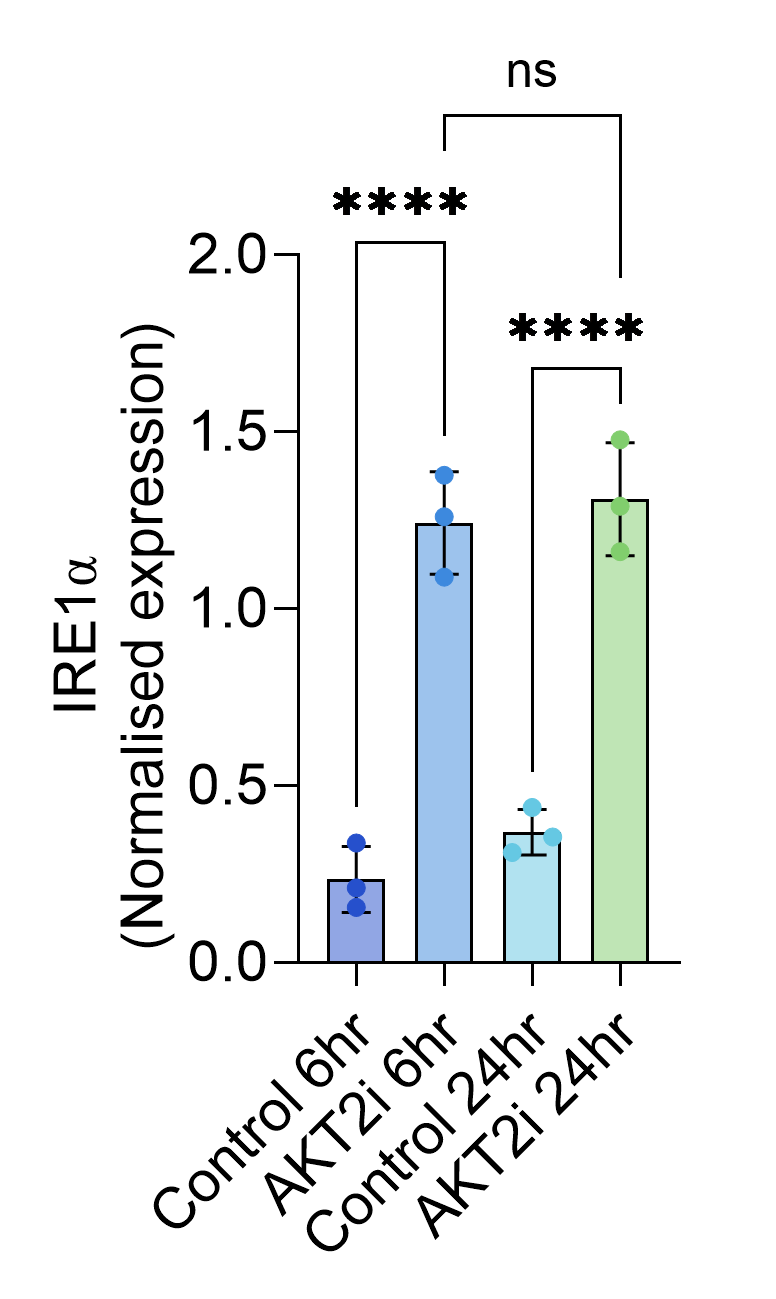

ERN1

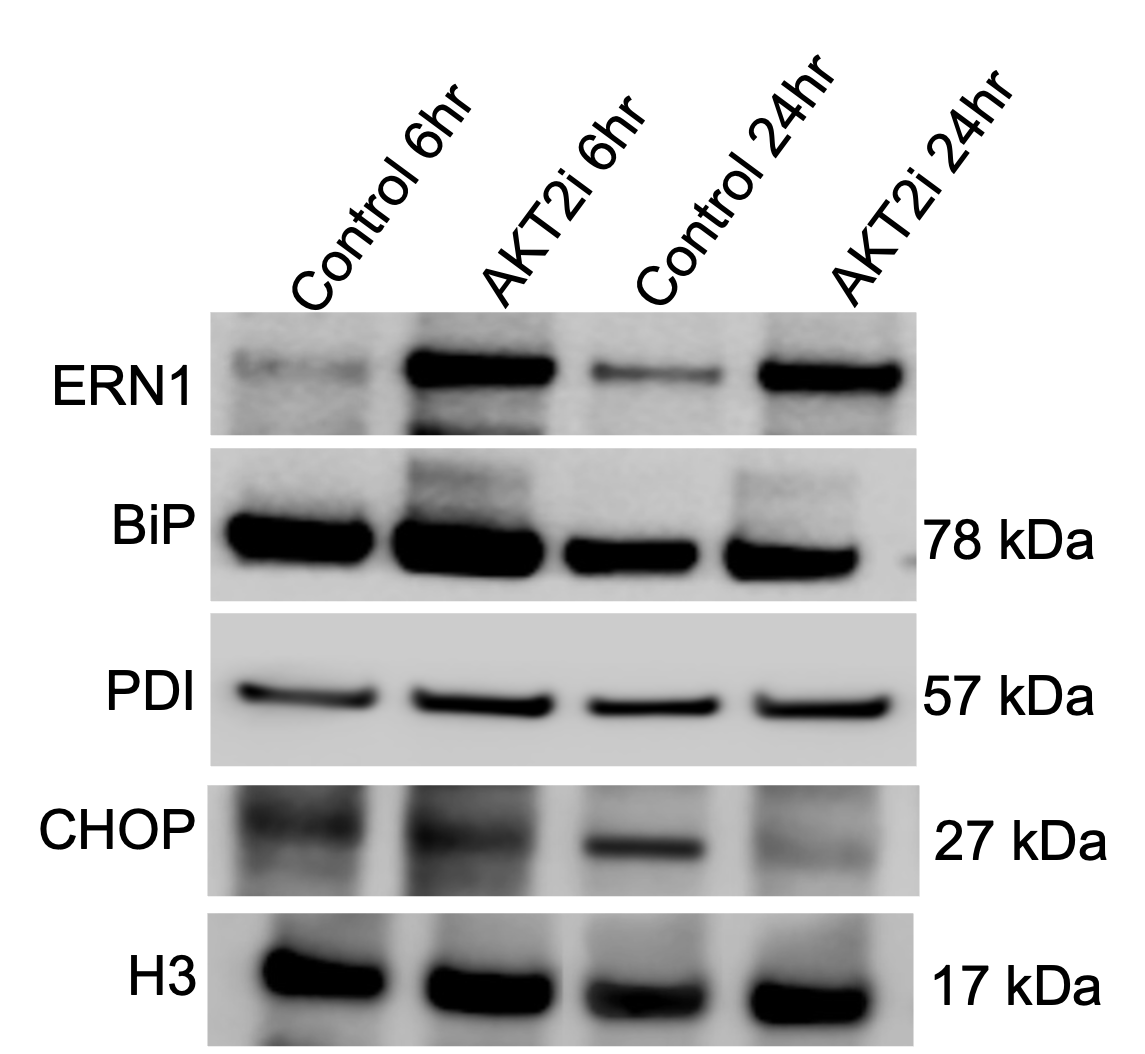

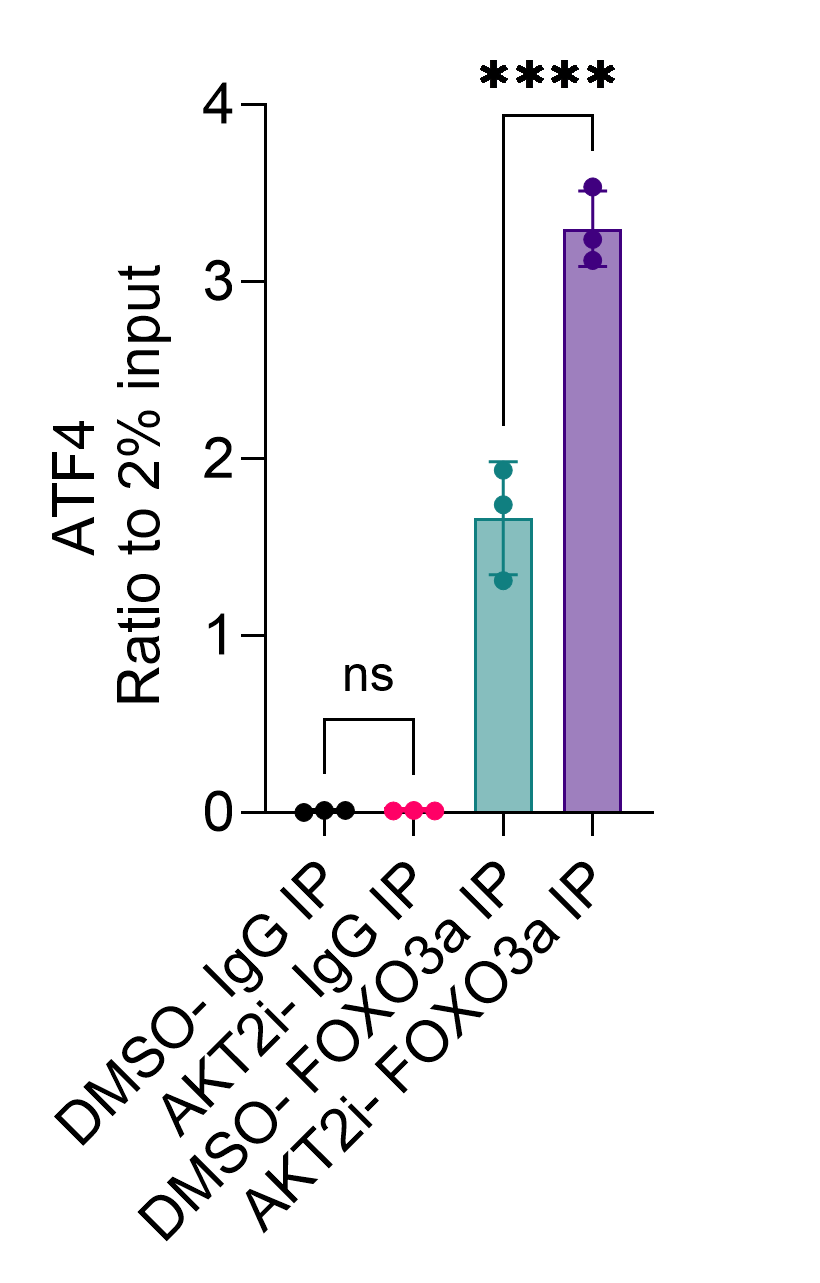

DMSO-IgG IP

AKT2i-IgG IP

DMSO-FOXO3 IP

AKT2i-FOXO3 IP

**Figure S5. AKT1-mediated phosphorylation of TERT regulates NRF2 nuclear localization and the UPR.** (**A**) Densitometry analysis showing normalized expression of phospho-PERK/ PERK in iPSC RPE *CFH* ^Y402H^ cells treated with DMSO and AKT2 inhibitor for 6 hours and 24hours. (**B**) Densitometry analysis showing nuclear ATF expression in iPSC RPE *CFH* ^Y402H^ cells treated with DMSO and AKT2 inhibitor for 6 and 24 h. (**C**) Representative immunoblots of whole-cell lysates probed for IRE-1α, BiP, PDI and CHOP. ACTB serves as a loading control. Blots are representative of n = 3 independent experiments. (**D-G**) Densitometric quantification of protein levels from (**C**). Signal intensities were normalized to ACTB. Horizontal lines represent mean ± SD. (**H-I**) Densitometry showing ratio of ATF4 levels to 2% input immunoprecipitated with (**H**) IgG and TERT, (**I**) IgG and FOXO3a IP from DMSO and AKT2i-treated RPE cells. (**J**) Fluorescence microscopy of human RPE showing localization of phosphor NRF2 (green) and CHOP (red). Cells were co-stained with DAPI (blue) to visualize nuclei. Scale bars=10μm. (**K**) Box whisker plots show the distribution of CHOP nuclear intensities from n =100 nuclei pooled from three independent experiments. (**L**) Automated single-cell quantification of nuclear phospho‑NRF2 intensity. Alexa Fluor 488‑phalloidin stains the plasma membrane, and cells were co‑stained with DAPI (blue) to visualize nuclei. (**M**) Box-and-whisker plots show the distribution of phospho‑NRF2 nuclear intensities from n = 100 nuclei pooled from three independent experiments. Statistical significance was determined using an unpaired Student’s t test. **** *p* < 0.0001. (**N**) Schematic showing dual function of ATF4 in activating nonselective macroautophagy and selective ERphagy for damaged ER fragments. Data represent mean ± SD. Statistical significance was determined by one-way ANOVA with Tukey’s post-hoc test. **p* < 0.05, ***p* < 0.01, *** *p* < 0.001, *****p* < 0.0001

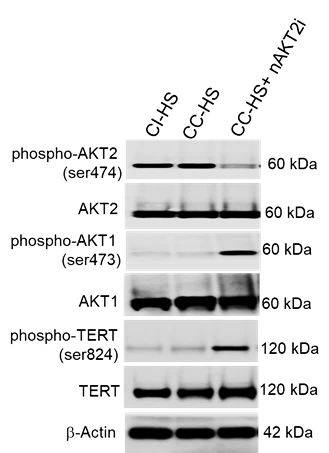

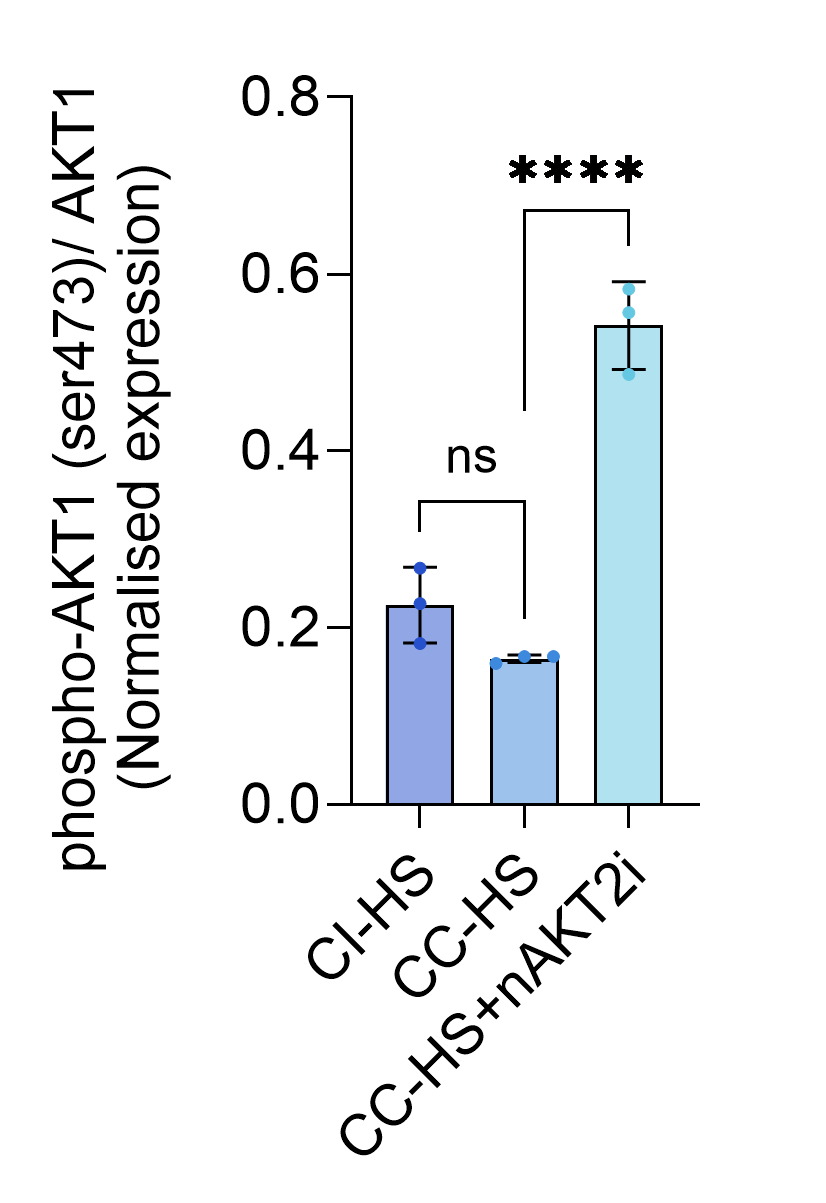

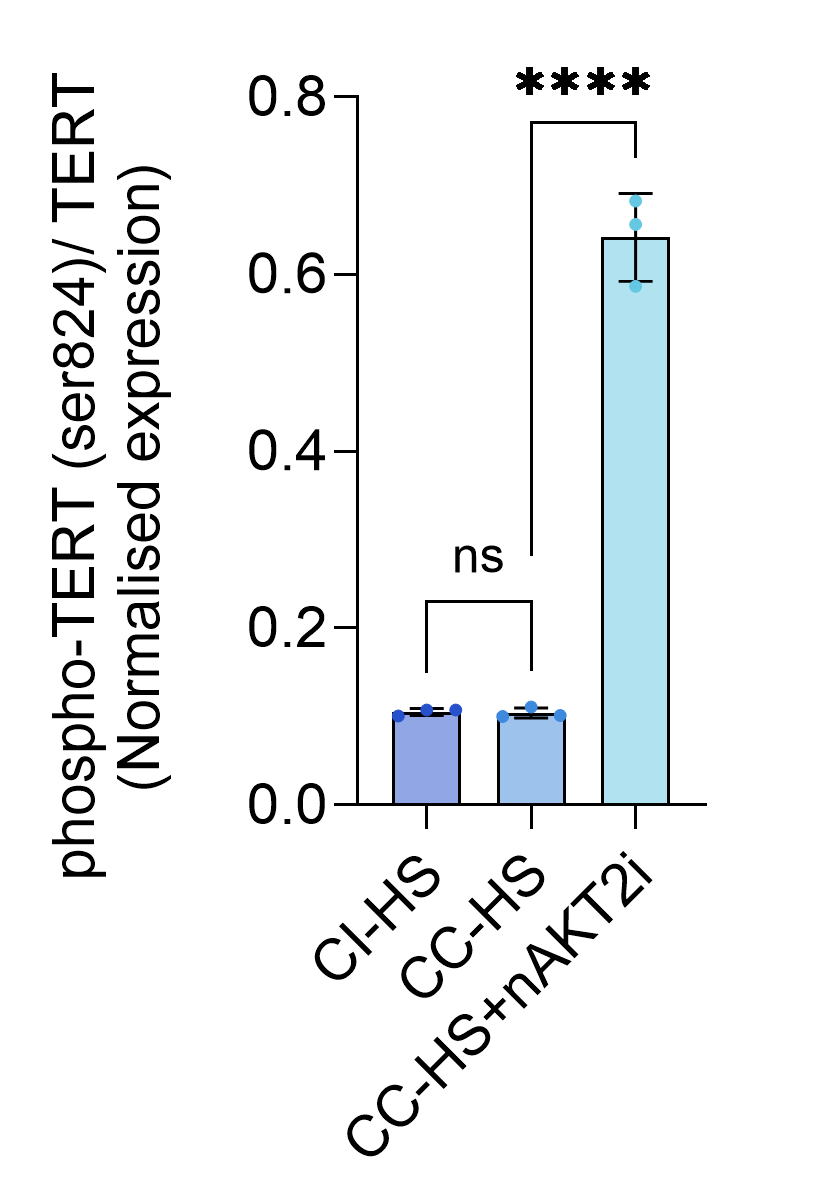

A

B

C

D

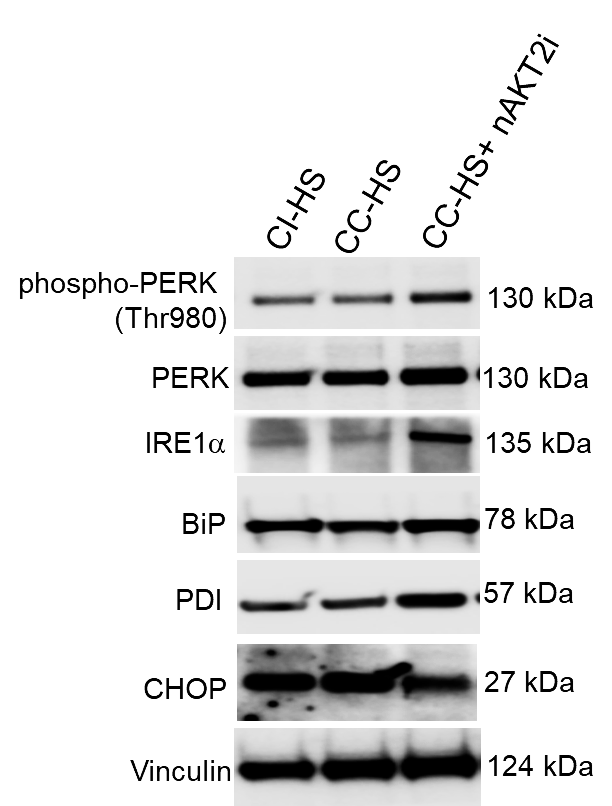

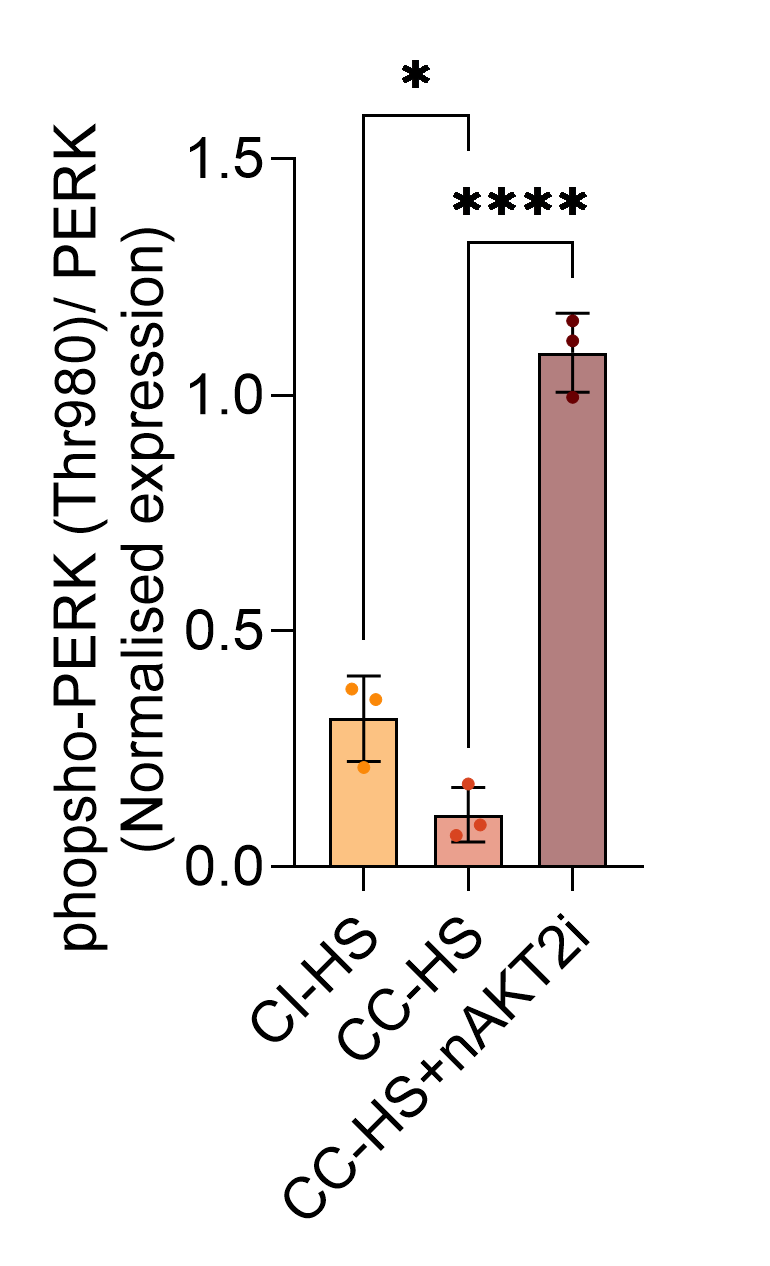

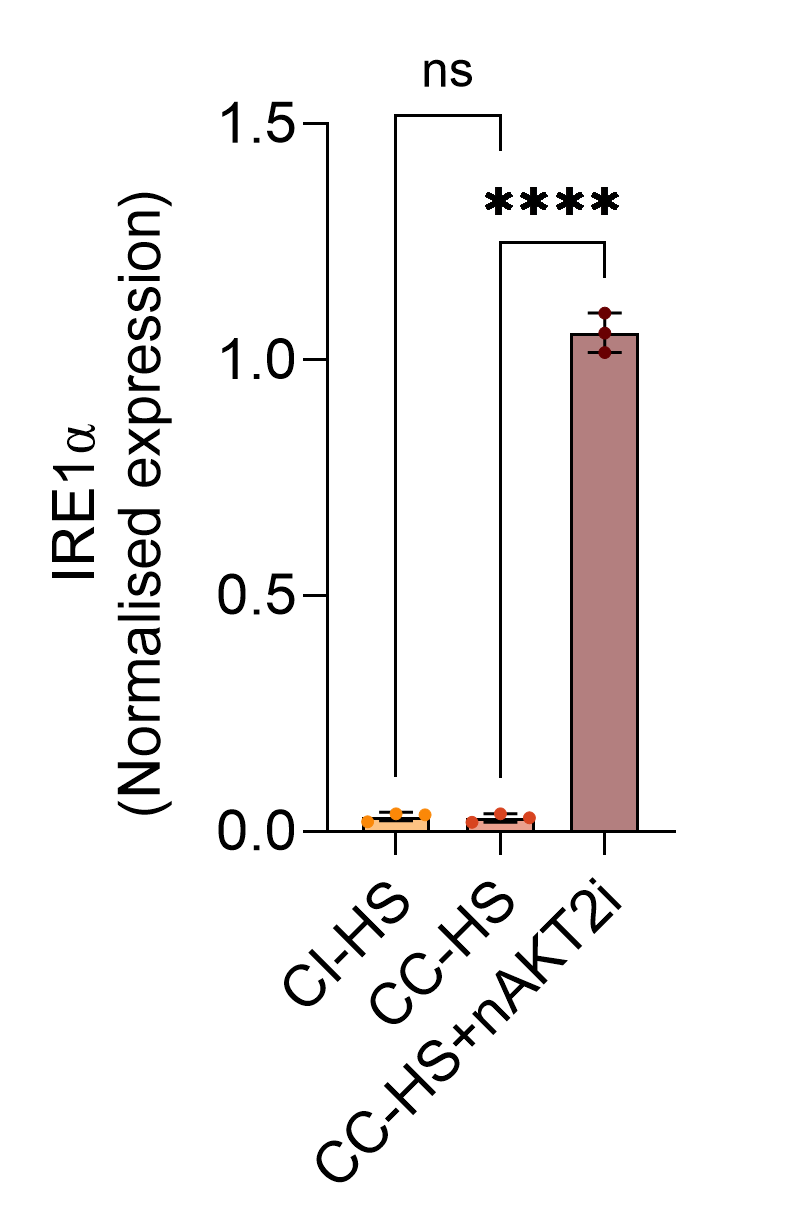

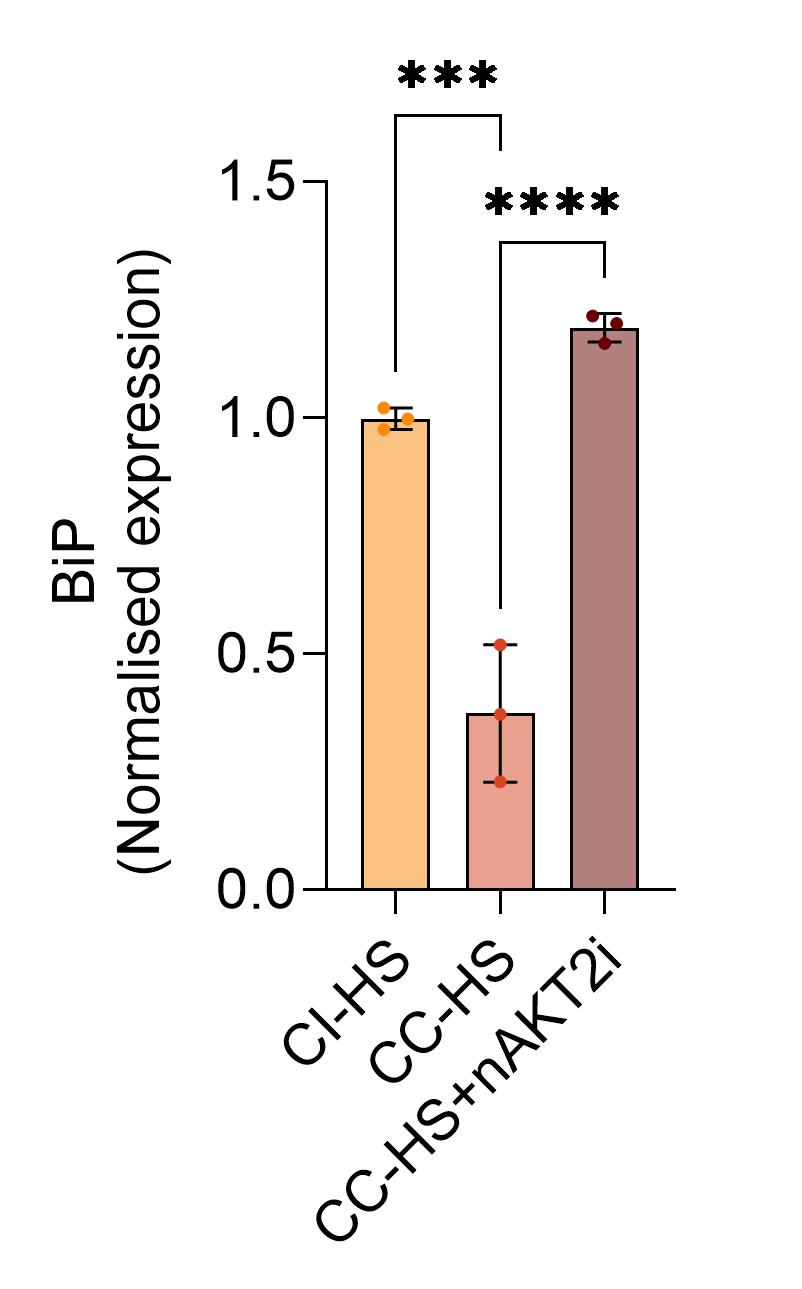

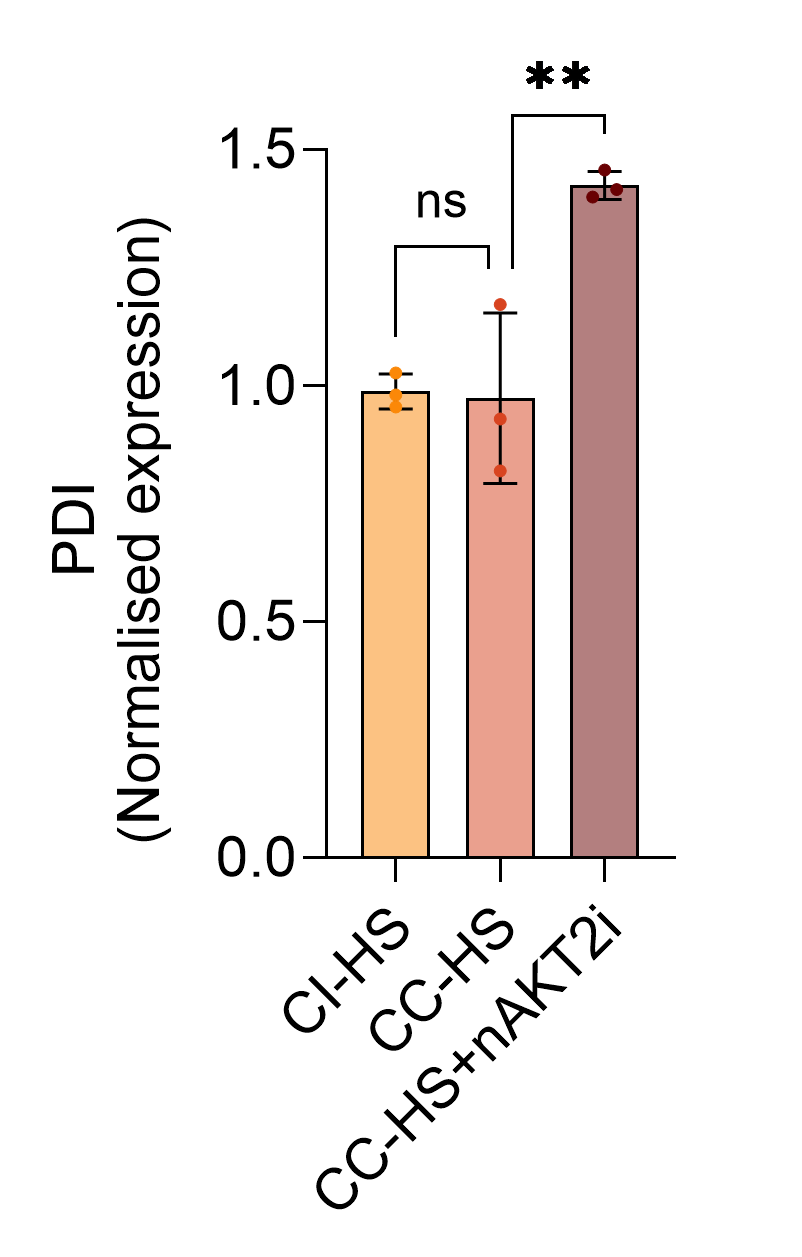

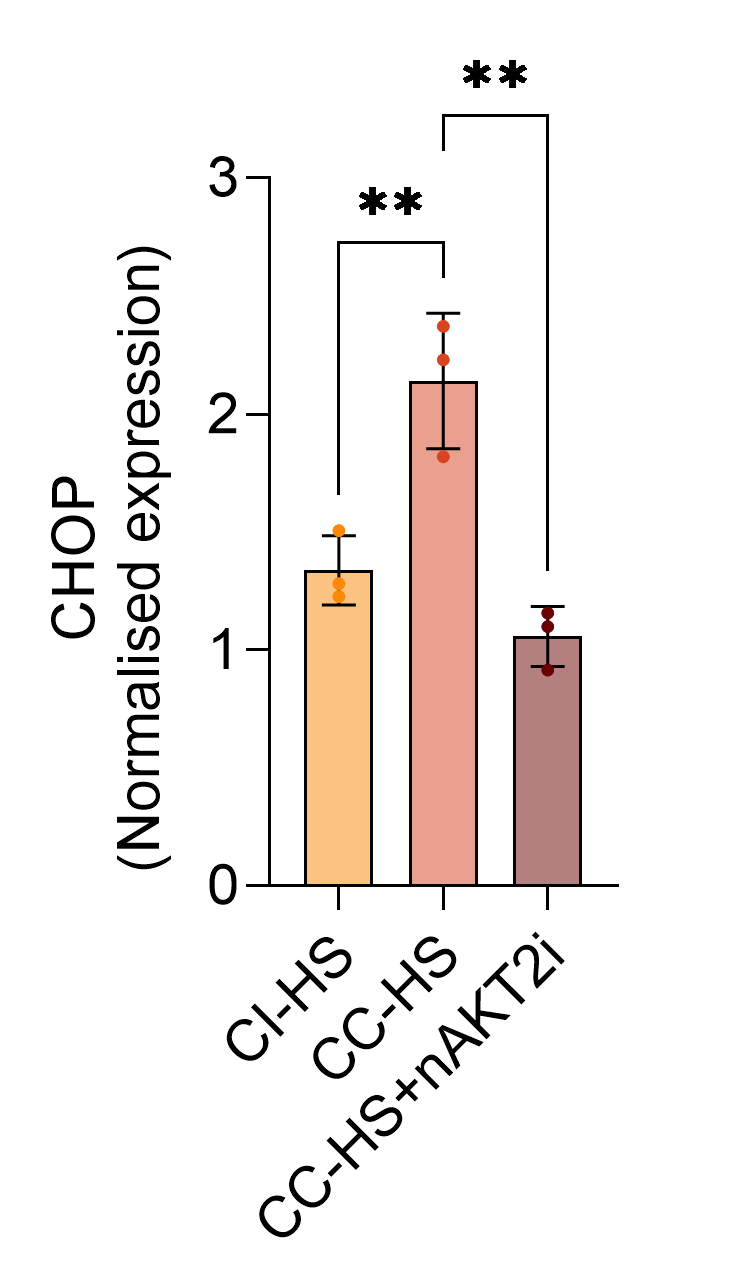

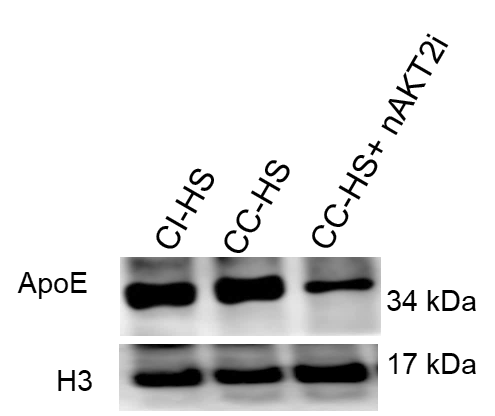

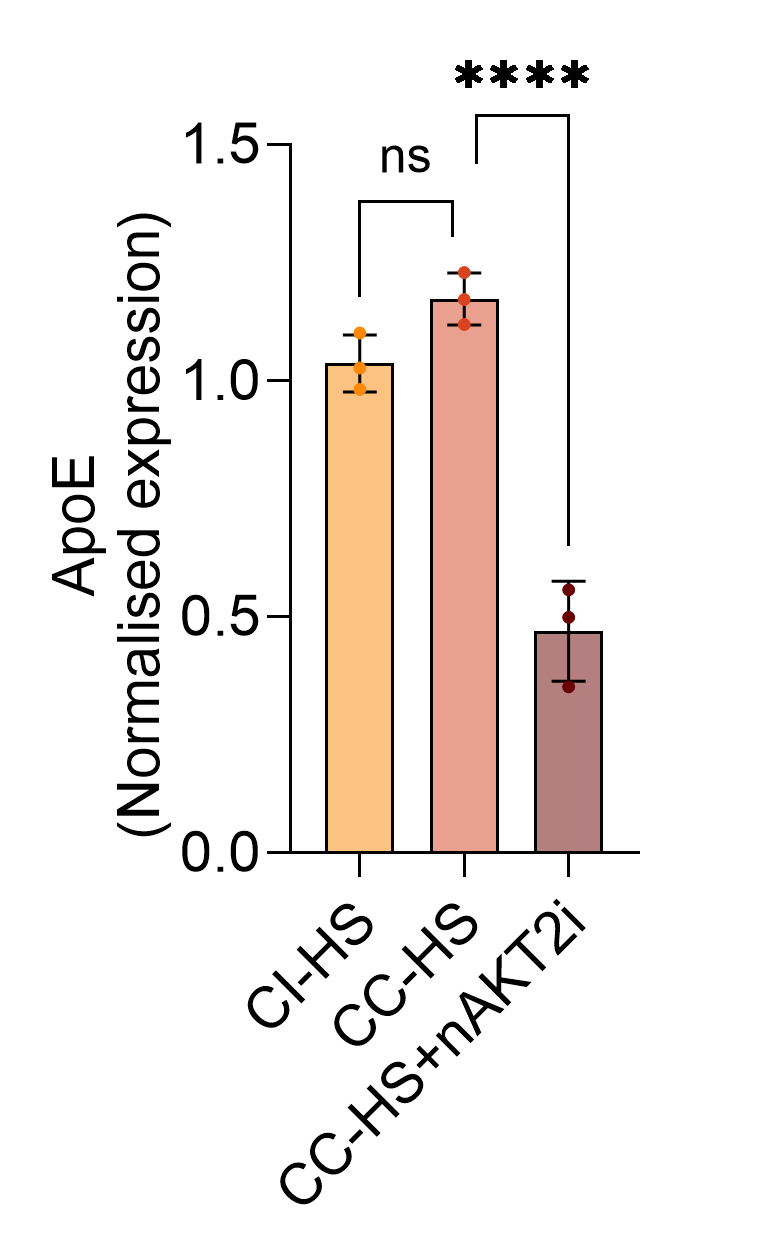

E

F

G

H

I

J

K

L

M

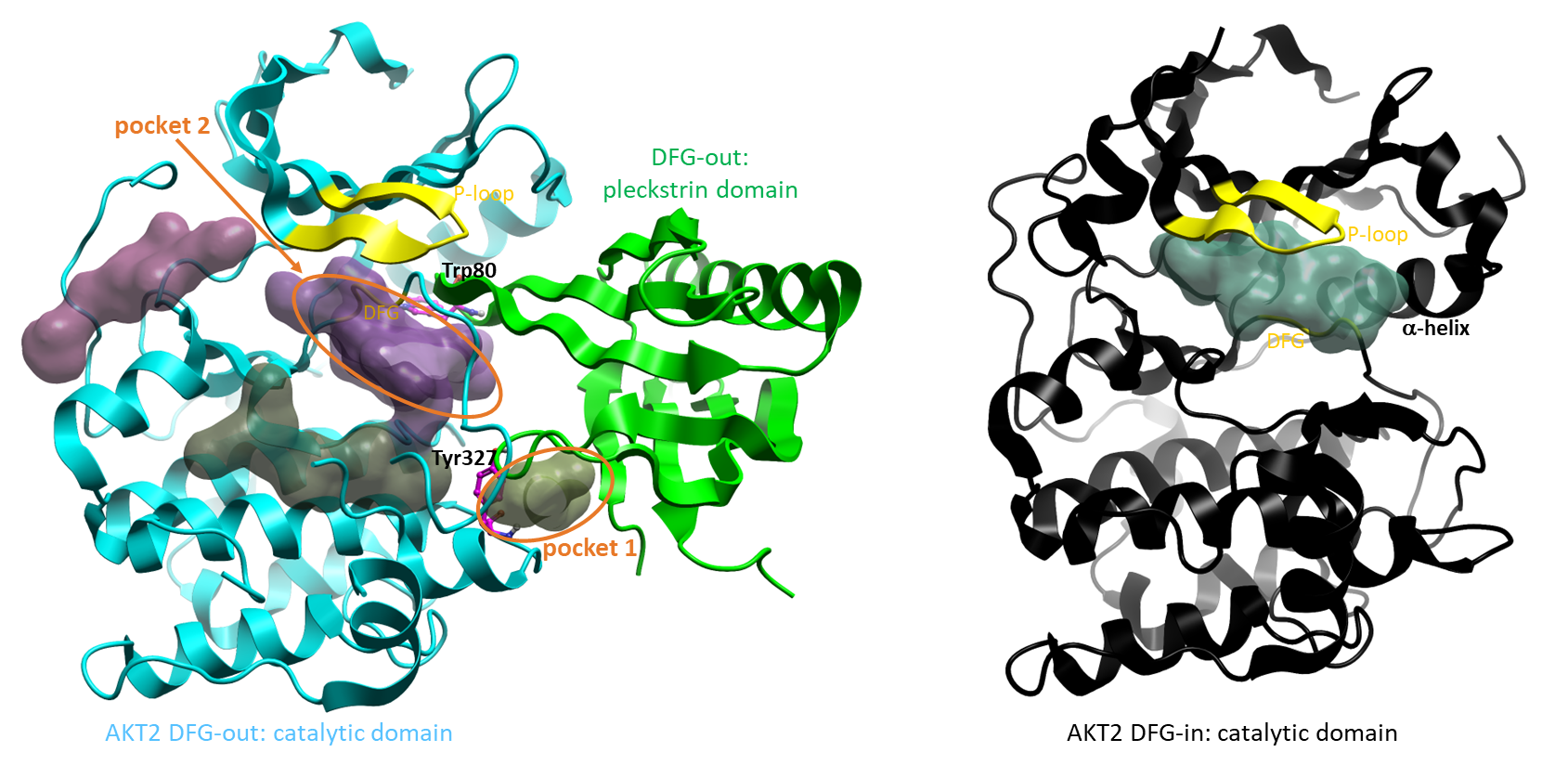

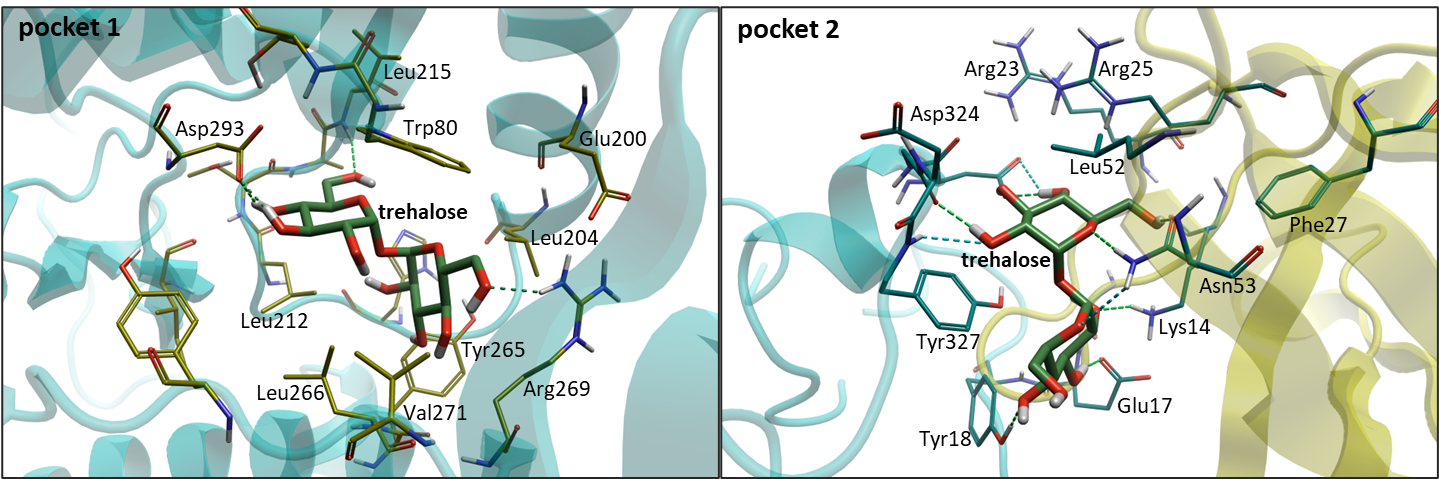

A

B

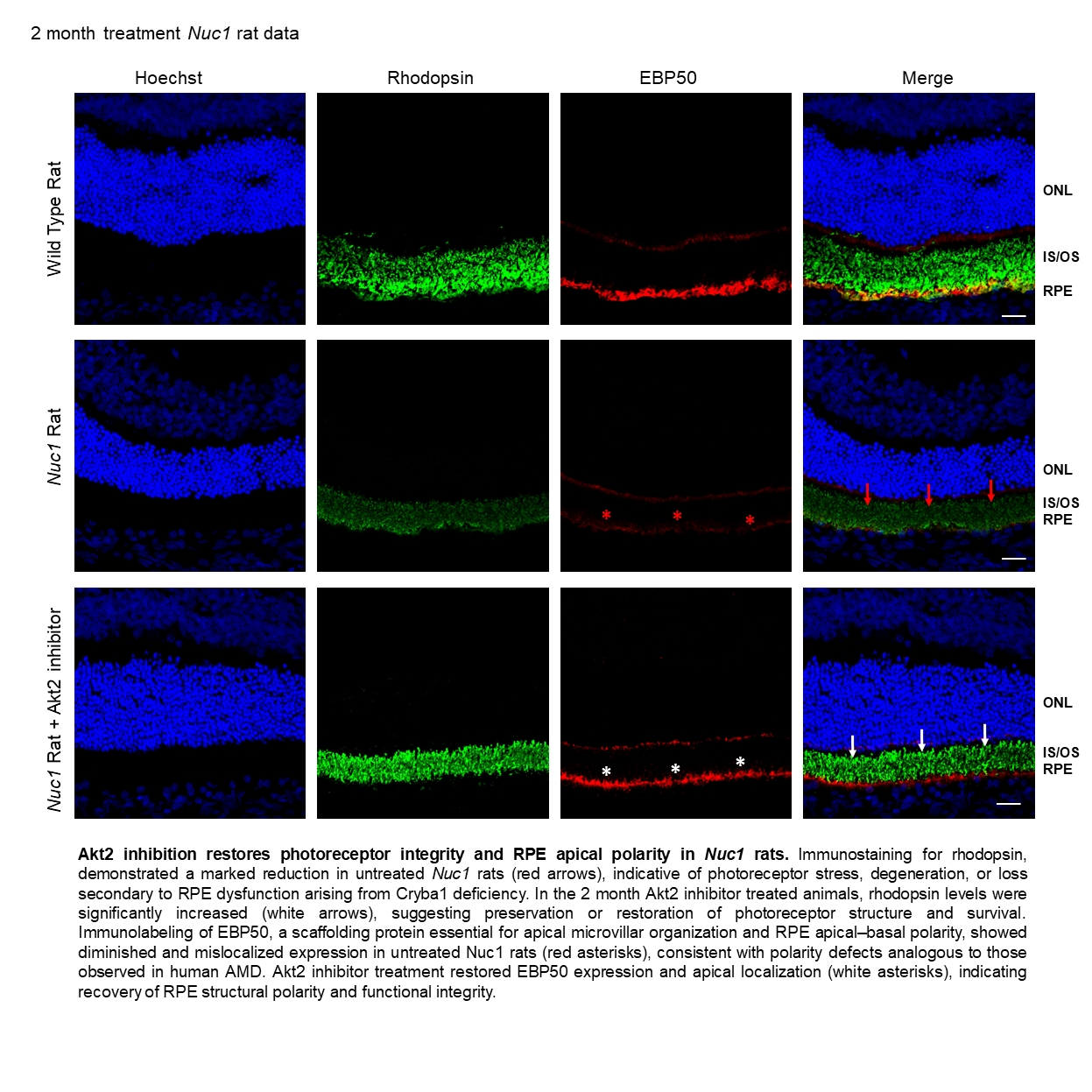

P

O

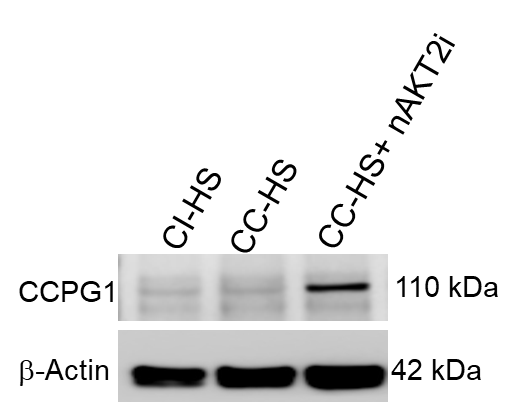

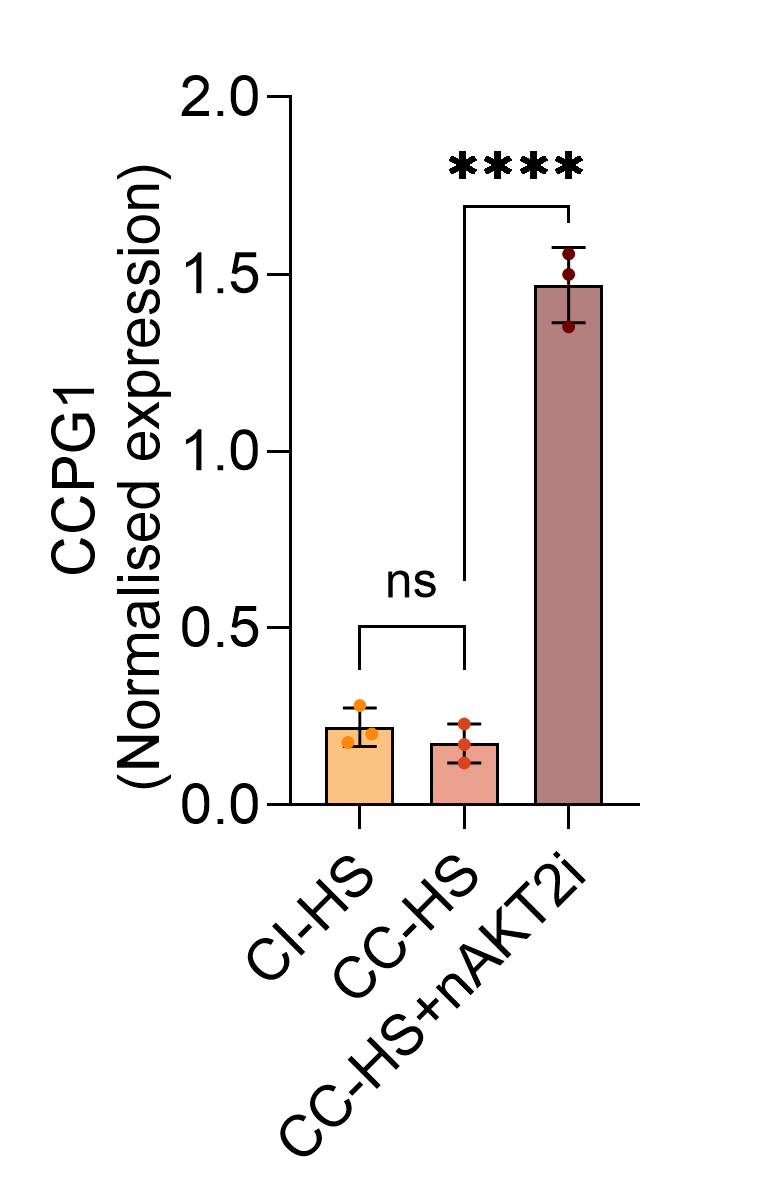

N

Q

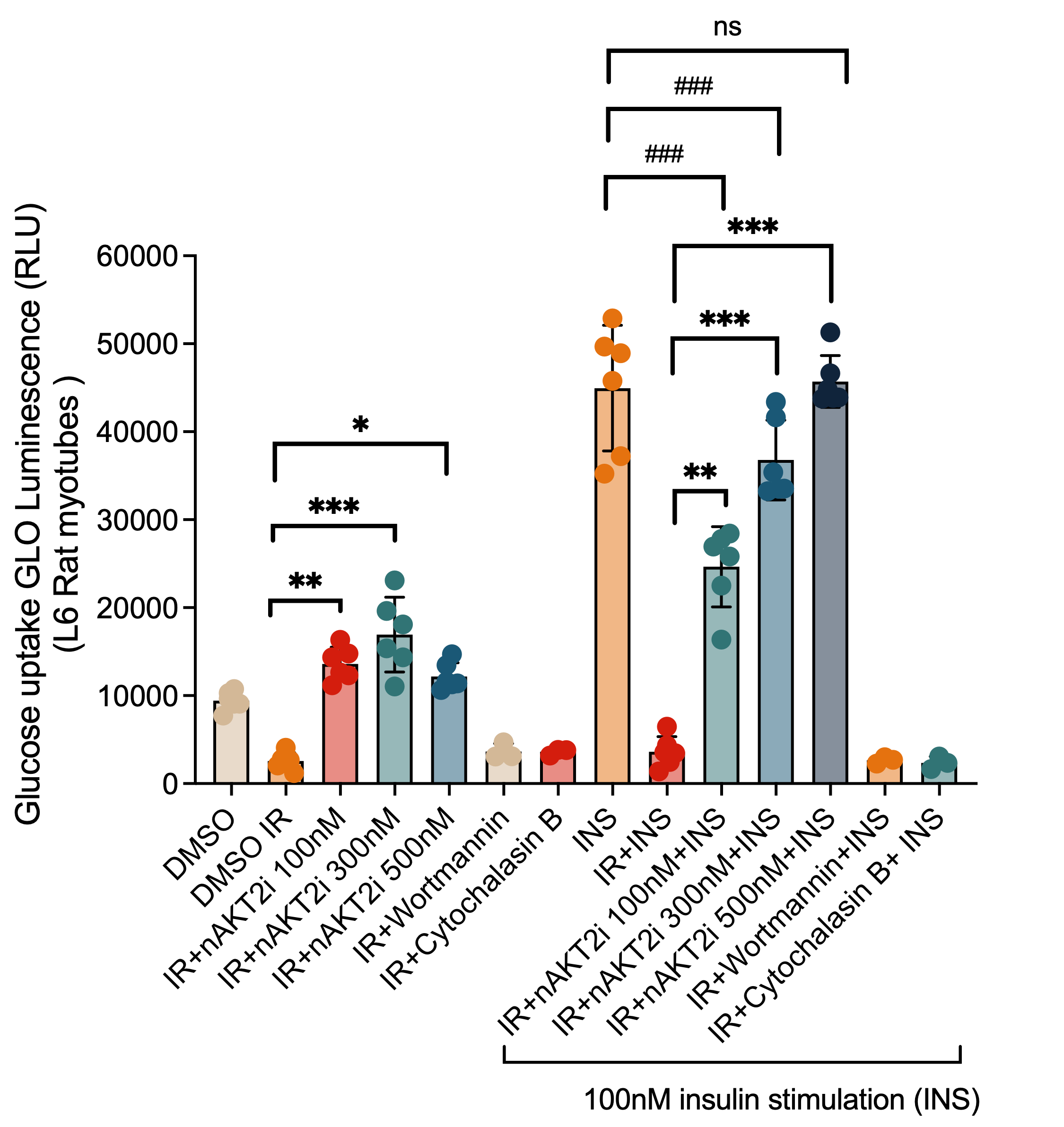

ACTB

VCL

ACTB

ERN1

VCL

ACTB

**Figure S6. Dual-pocket inhibition of AKT2 relieves ER stress and activates ERphagy for cellular adaptation.** (**A**) Structural analysis showing the identification of potential allosteric binding pockets for trehalose in the DFG-out conformation of AKT2. Comparative structural models of the activated (PDB: 1O6L) and inactivated (PDB: 8Q61) states demonstrate that only the DFG-out conformation allows binding to the pleckstrin domain, with two candidate binding pockets identified. (**B**) Molecular docking analysis of trehalose binding in pockets 1 and 2, demonstrating favorable binding modes supported by thermodynamically stable water molecules and low energy glycosidic bond torsion values. (**C**) Immunoblot analysis of phospho-AKT1, phospho-AKT2, phospho-TERT, AKT1, AKT2 and TERT expression in “disease in a dish degeneration model’ iRPE cells (H/H) exposed to complement inactive human serum (CI-HS), complement competent human serum (CC-HS), and CI-HS exposed iRPE cells treated with nAKT2i (CC-HS+nAKT2i). β-actin serves as the loading control. (**D-E**) Densitometric quantification of (**D**) phospho-AKT1 to Total AKT1 and (**E**) phospho-TERT to Total TERT. Data are normalized to ACTB and expressed as normalized expression. n = 3 independent biological replicates; error bars represent mean ± SD. Statistical significance was determined using one-way ANOVA with Tukey’s post-hoc test. ***p* < 0.01, *****p* < 0.0001, ns= not significant. (**F**) Immunoblot analysis of UPR markers phospho-PERK, PERK, IRE-1α, Bip, PDI and CHOP expression in iRPE cells (H/H) exposed to complement inactive human serum (CI-HS), complement competent human serum (CC-HS), and CI-HS exposed iRPE cells treated with nAKT2i (CC-HS+nAKT2i). VCL serves as the loading control. (**G-K**) Densitometry analysis of UPR markers in Fig S7F. Data are normalized to VCL and expressed as normalized expression. n = 3 independent biological replicates; error bars represent mean ± SD. Statistical significance was determined using one-way ANOVA with Tukey’s post-hoc test. **p* <0.05, ***p* < 0.01, *****p* < 0.0001, ns= not significant**.** (**L**) Immunoblot analysis of ApoE expression in iRPE cells (H/H) exposed to complement inactive human serum (CI-HS), complement competent human serum (CC-HS), and CI-HS exposed iRPE cells treated with nAKT2i (CC-HS+nAKT2i). H3 serves as the loading control. (**M**) Densitometric quantification of ApoE levels in iRPE cells (H/H) exposed to CI-HS, CC-HS and CC-HS+nAKT2i. n = 3 independent biological replicates; error bars represent mean ± SD. Statistical significance was determined using one-way ANOVA with Tukey’s post-hoc test. *****p* < 0.0001, ns= not significant. (**N**) Immunoblot analysis of ERphagy receptor CCPG1 expression in iRPE cells (H/H) exposed to complement inactive human serum (CI-HS), complement competent human serum (CC-HS), and CI-HS exposed iRPE cells treated with nAKT2i (CC-HS+nAKT2i). β-actin serves as the loading control. (**O**) Densitometric quantification of CCPG1 levels in iRPE cells (H/H) exposed to CI-HS, CC-HS and CC-HS+nAKT2i. n = 3 independent biological replicates; error bars represent mean ± SD. Statistical significance was determined using one-way ANOVA with Tukey’s post-hoc test. *****p* < 0.0001, ns= not significant. (**P**) Functional assays show AKT isoform rebalancing favors insulin-stimulated glucose uptake in palmitate mediated insulin resistant skeletal muscle L6 Rat myotubes. L6 rat myotubes were treated with varying concentrations of nAKT2i (100nM, 300nM, 500nM) in the presence or absence of 100nM Insulin (INS). Glucose uptake was measured via Glucose uptake Glo-Luminescence (RLU). Wortmannin (100nM) and Cytochalasin B (10 μM) was used as a negative control for PI3K/AKT and transporter-mediated uptake. (**Q**) Representative confocal images of retinal cross-sections from wild-type, *Nuc1*, and *Nuc1* + nAKT2 inhibitor–treated rats. Sections were stained with Hoechst (blue, nuclei), rhodopsin (green), and EBP50 (red). *Nuc1* rats carry a *Cryba1* mutation that causes RPE dysfunction and retinal degeneration. After 2 months, vehicle‑treated *Nuc1* retinas show reduced rhodopsin signal (red arrows), whereas nAKT2 inhibitor–treated animals display increased rhodopsin (white arrows), consistent with improved photoreceptor integrity. EBP50 expression and apical localization are also restored following Akt2 inhibition, indicating recovery of RPE polarity. Scale bar = 20μm. Data represent mean ± SD; individual biological replicates are shown as dots. Statistical significance was determined by two-way ANOVA; **p* < 0.05, ***p* < 0.01, ****p* < 0.001; ns=no significance; ^###^ *p*<0.001 denotes significance relative to insulin-only control.

**Table 1:**

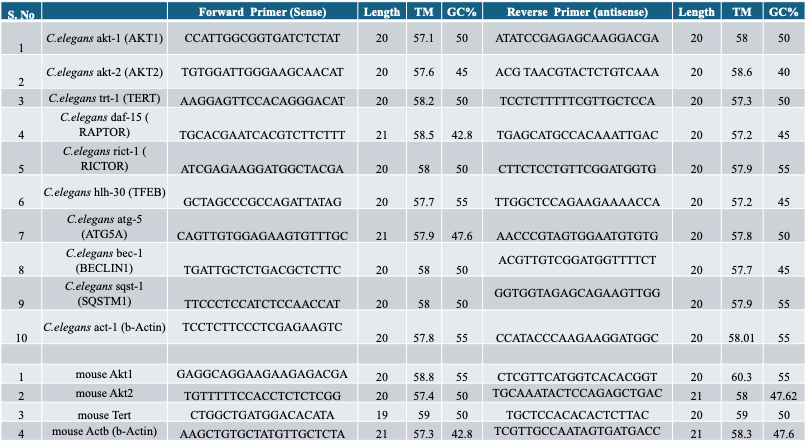

**Table 2:**

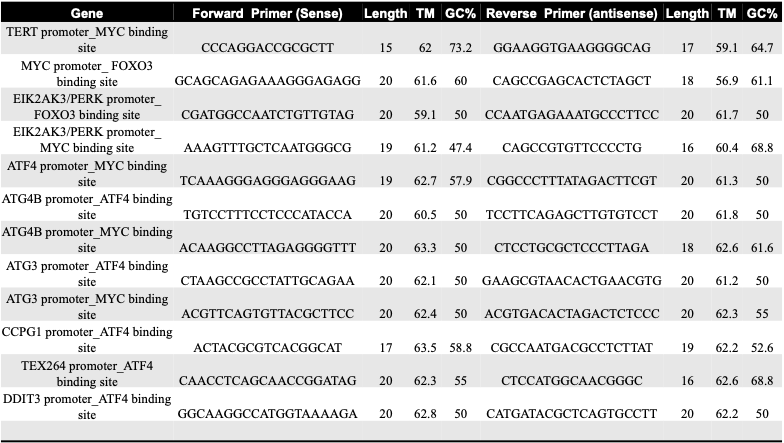
